# E2 displacement of CIP2A from TOPBP1 activates the DNA damage response during papillomavirus life cycles

**DOI:** 10.1101/2025.10.15.682653

**Authors:** Apurva T. Prabhakar, Claire D. James, Allison Q. Nguyen, Phoebe V. Bridy, Xu Wang, Ryan J. Zoldork, Jenny D. Roe, Austin J. Witt, Srimanya S. Panidepu, Molly L. Bristol, Krista Dalton, Andreas Wieland, Ahmed Diab, Anthony C. Faber, Elliot J. Androphy, Devraj Basu, Paul F. Lambert, Megan E. Spurgeon, Iain M. Morgan

## Abstract

The papillomavirus life cycle is intricately linked to epithelial differentiation, and the virus manipulates the differentiation process to facilitate viral production. One such manipulation is activation of the DNA damage response (DDR) which promotes viral replication via homologous recombination. This report demonstrates that the papillomavirus transcription/replication/segregation factor E2 activates the DNA damage response (DDR). During differentiation, E2 displacement of CIP2A from TOPBP1 causes CIP2A to bind and inhibit PP2A resulting in DDR activation via ATM phosphorylation. The DDR promotes inhibitory interaction of DBC1 with the class III deacetylase SIRT1, which further boosts the DDR via increased acetylation and stability of viral and host proteins. E2 forms a complex with TOPBP1 and ATM, while preventing ATR activation by blocking TOPBP1-ATR interaction. This “ATM up ATR down” phenotype promotes viral replication via ATM promotion of homologous recombination, and cell proliferation via inhibition of ATR. We demonstrate this mechanism of DDR activation in multiple systems: keratinocytes expressing only E2, in foreskin keratinocytes immortalized by HPV16, in HPV16 positive keratinocytes derived from a cervical lesion, in pre-neoplastic lesions induced by mouse papillomavirus MmuPV1, in head and neck cancer cell lines that retain E2 expression, and in HPV16 positive oropharyngeal patient derived xenografts that retain E2 expression. ATM inhibition preferentially killed cells expressing E2, presenting a novel strategy for treating HPV early preneoplasia and a large subset of HPV+ oropharyngeal cancers retaining E2 expression and episomal genomes.

## Introduction

Human papillomaviruses (HPV) are causative in around 5% of all cancers, with HPV16 being the most prevalent (zur Hausen, 2009). The HPV life cycle occurs in differentiating epithelium and viral production is intricately connected to squamous epithelial differentiation. The virus manipulates differentiation to produce infectious particles that egress from the upper layers of differentiated epithelium (Schiffman et al., 2016). During the viral life cycle the DNA damage response (DDR) is activated (Moody and Laimins, 2009). HPV activation of the DDR promotes the use of homologous recombination (HR) to replicate the viral genome, and HR factors are required for optimal viral DNA replication and the viral life cycle (Akagi et al., 2023; Anacker et al., 2014; Chappell et al., 2015; Das et al., 2019; Das et al., 2017; Moody, 2022; Sakakibara et al., 2013). The exact mechanism of HPV activation of the DDR is unknown; while viral oncogenes E6 and E7 can induce DNA damage when overexpressed in human cells, they are unable to significantly activate the DDR (Duensing et al., 2001; Duensing and Munger, 2002; Hoppe-Seyler et al., 2018; James et al., 2020b; McLaughlin-Drubin and Munger, 2009; Munger et al., 2001). Overexpression of the viral helicase E1 activates the DDR, presumably by non-specifically unwinding host DNA that would be recognized by host factors as DNA damage triggering the DDR (Kadaja et al., 2009; Kadaja et al., 2007; Sakakibara et al., 2011).

Recently, we reported a mitotic activation of the DDR in keratinocytes expressing HPV16 E2 (Prabhakar et al., 2024). E2 has two major domains: a carboxyl-terminal DNA binding and dimerization domain that interacts with 12bp palindromic sequences in the viral long control region (LCR), and an amino-terminal protein interaction platform (McBride, 2013; Morgan, 2025). During mitosis, E2 acts as a bridge between the viral and host genomes allowing the viral genome to hitchhike onto host DNA to ensure segregation to daughter nuclei. Binding of E2 with TOPBP1 is required for interaction with mitotic chromatin and plasmid segregation, and is promoted by CK2 phosphorylation of E2 at serine 23 (Prabhakar et al., 2022; Prabhakar et al., 2021; Prabhakar et al., 2023; Prabhakar and Morgan, 2025). E2 induced mitotic activation of the DDR requires interaction with TOPBP1 (Prabhakar et al., 2024). TOPBP1 is a scaffold protein containing 9 BRCT (BRCA1 carboxyl terminal) domains that act as interacting modules for phosphorylated proteins and damaged DNA structures (Wardlaw et al., 2014). TOPBP1 is involved in all aspects of DNA metabolism including transcription, replication, DNA damage recognition, signaling and repair (Wardlaw et al., 2014). TOPBP1 is active during mitosis to promote genome stability and DNA replication, while DNA damage during mitosis promotes TOPBP1 recruitment to host chromatin via MDC1 interaction; MDC1 binds to γH2AX at sites of DNA damage (Adam et al., 2021; Bagge et al., 2020; Bang et al., 2013; Bang et al., 2011; Broderick et al., 2015; Germann et al., 2014; Leimbacher et al., 2019; Pedersen et al., 2015).

HPV induces a G2/M-like cellular environment in differentiating epithelium that promotes the viral life cycle (Davy and Doorbar, 2007). This provides an environment that allows viral replication due to availability of host polymerases, while restricting host replication. This enhances the nucleotide pool available to the virus for genome replication. The link between E2 activation of a mitotic DDR and the ability of HPV to generate the G2/M phenotype, and the requirement of the DDR for the viral life cycle, prompted us to investigate whether E2 activates the DDR in differentiating keratinocytes. Here we describe a novel mechanism that E2 uses to activate the DDR during differentiation in a TOPBP1 interaction dependent manner, generating an “ATM up ATR down” DDR signaling phenotype. This activity of E2 persists in premalignant HPV-induced lesions and human-derived models of HPV-related head and neck cancers, which, in contrast to cervical cancers, often maintain episomal HPV genomes. The vulnerability to ATM inhibition created by this E2 activity defines a novel approach for combating papillomavirus infections and certain HPV-induced cancers.

## Results

### E2 generates an “ATM up ATR down” phenotype in differentiating keratinocytes

During mitosis E2 interaction with TOPBP1 is required for the plasmid segregation function of E2; E2 also enhances CHK2 phosphorylation during mitosis and activates the DDR (Prabhakar et al., 2024). Figure 1A demonstrates that in N/Tert-1 cells (hTERT immortalized human foreskin cells) stably expressing wild type E2 (N/Tert-1+E2-WT), ATM levels and its phosphorylation increased following 3 days of calcium-induced differentiation (Day 3), with a corresponding increase in CHK2 and pCHK2 levels (compare lane 6 with lane 3 in the empty vector N/Tert-1 control cells). Strikingly, there is a reduction of ATR levels in the differentiated N/Tert-1+E2-WT cells at day 3 compared with both growing (G, lane 4) and cells immediately prior to calcium addition (Day 0, lane 5), and with the differentiated N/Tert-1+Vec cells (lane 3). CHK1 and pCHK1 levels were also reduced following differentiation of N/Tert-1+E2-WT cells, consistent with reduced ATR signaling, as CHK1 is a direct downstream effector of ATR. E2 and CHK2 levels increase following differentiation, as they do during mitosis (Prabhakar et al., 2024). Therefore, E2 induces an “ATM up ATR down” phenotype in differentiating keratinocytes. We compared these findings to those with cells expressing an E2 mutant, E2-S23A, deficient in CK2 phosphorylation at Ser23 and unable to interact with TOPBP1 (Prabhakar et al., 2021). We did not observe the “ATM up ATR down” phenotype following differentiation of N/Tert-1+E2-S23A cells (lane 9). Therefore, the E2-TOPBP1 interaction is required for the “ATM up ATR down” DDR activation induced by the E2 protein.

**Figure 1.**
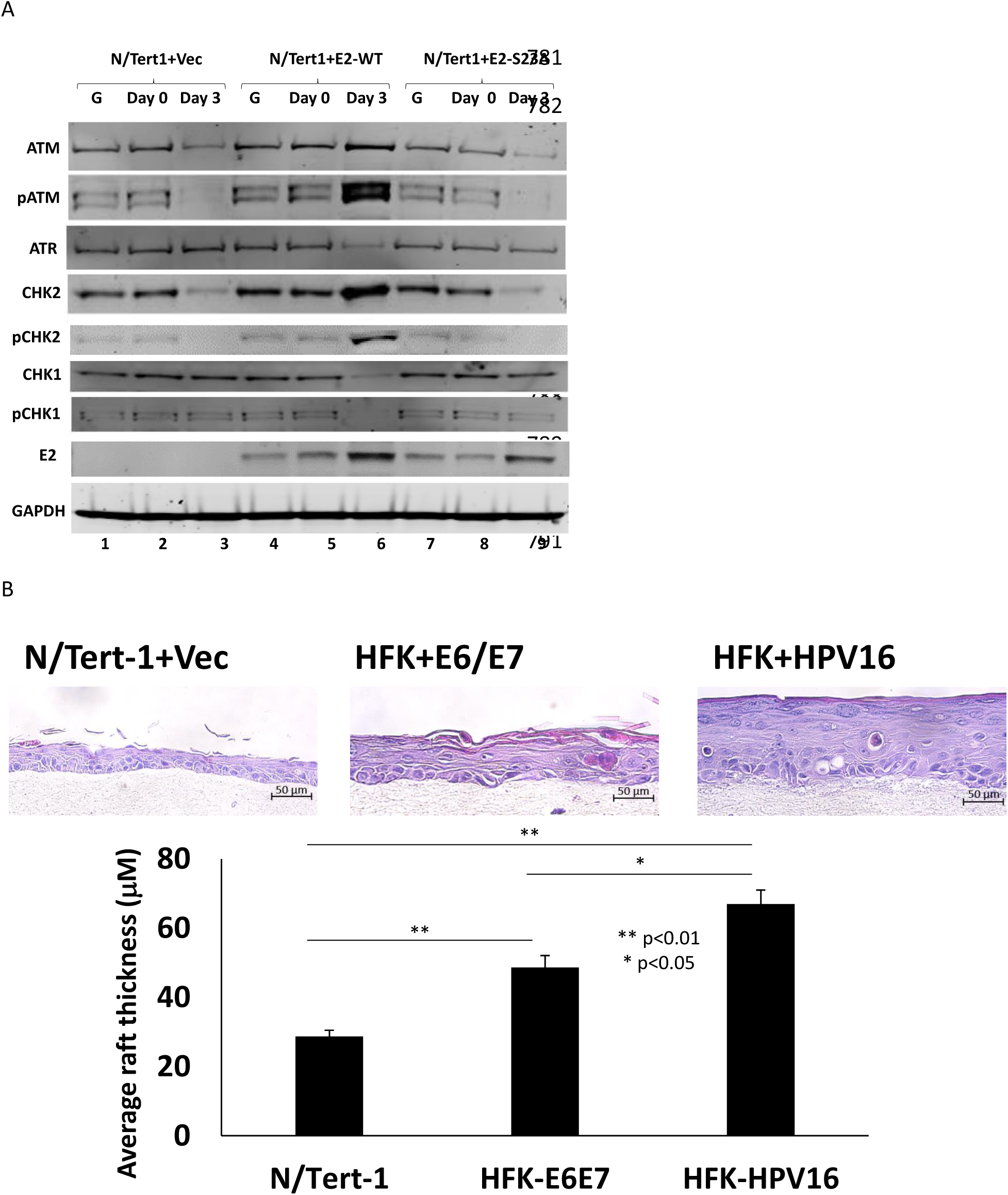
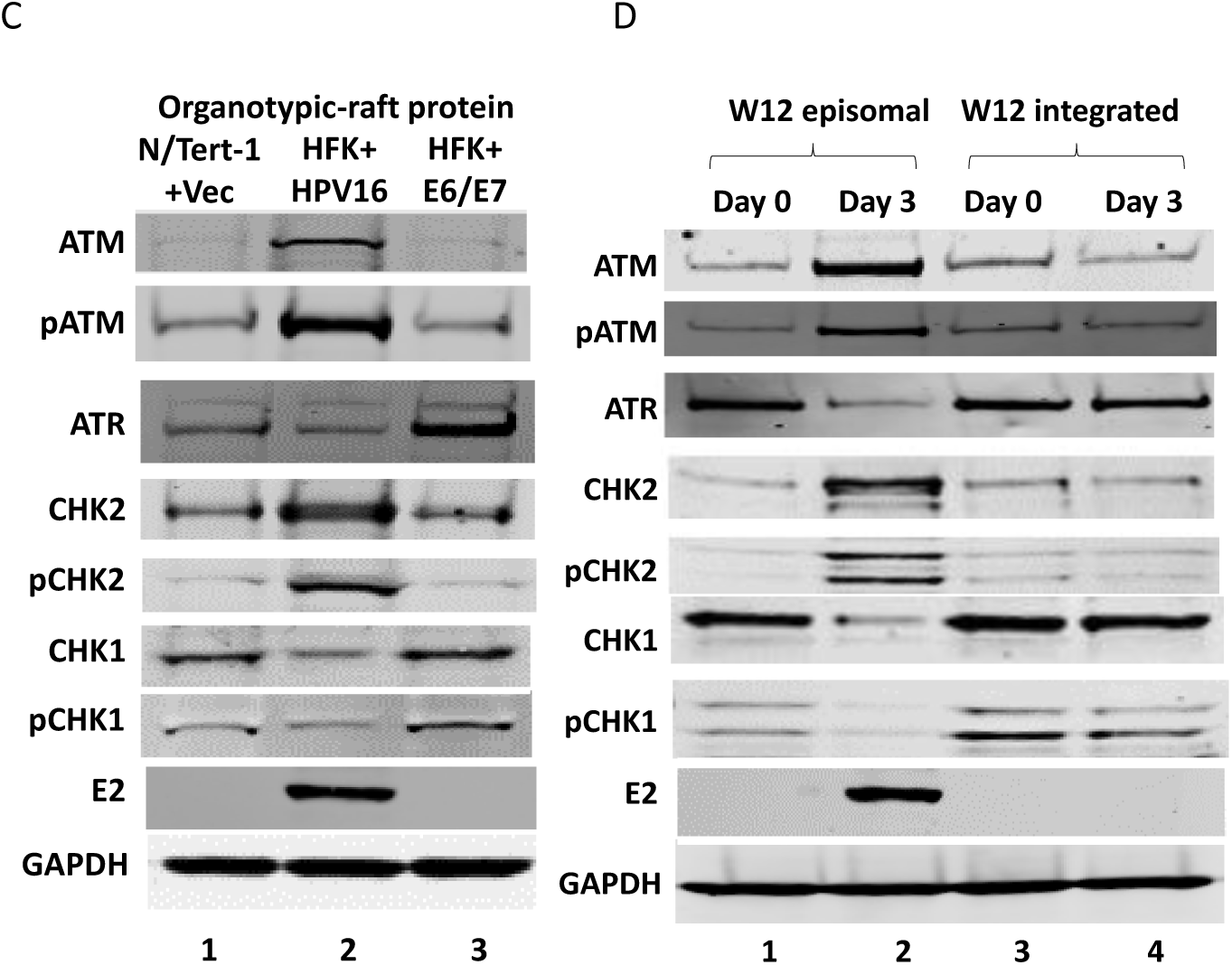
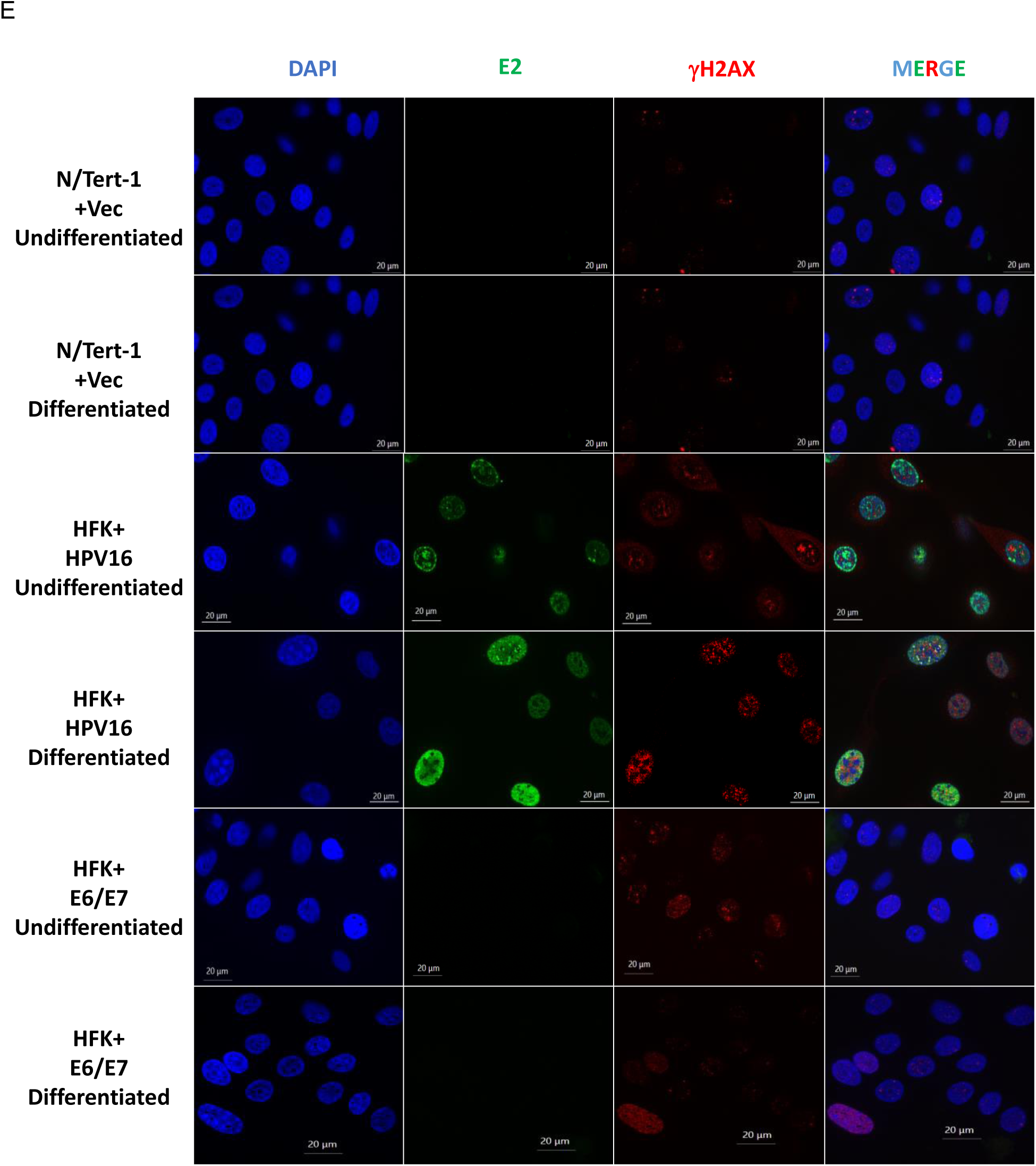
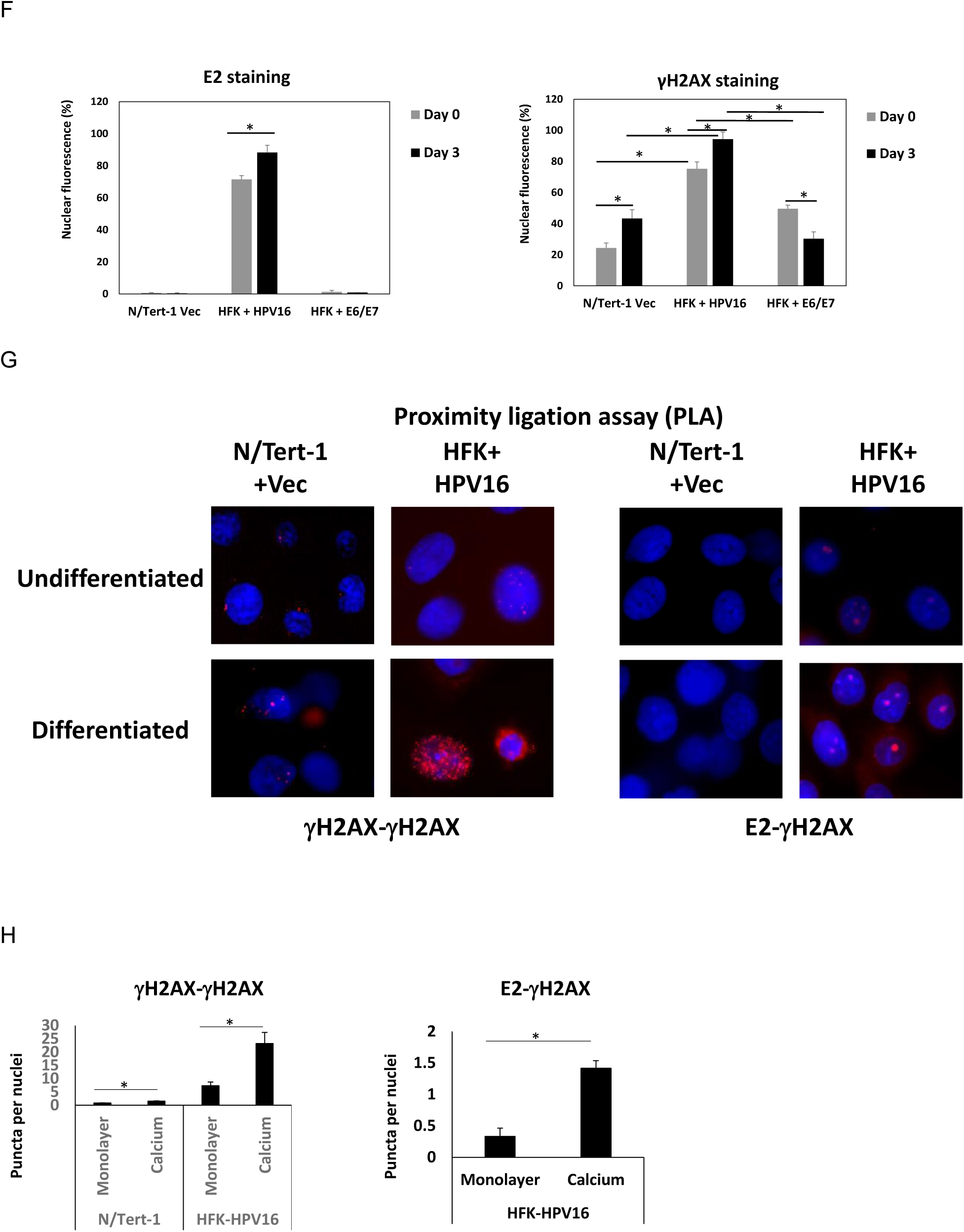
E2 induces an “ATM up ATR down” phenotype in differentiating epithelium. A. Proteins were extracted from N/Tert-1 cells with an empty vector control (N/Tert-1+Vec), expressing wild type E2 (N/Tert-1+E2-WT) or expressing an E2 that fails to interact with TOPBP1 (N/Tert-1+E2-S23A) that were growing (G), plated for differentiated with no calcium added (Day 0) or were induced to differentiate for 3 days following calcium treatment (Day 3). Western blotting with the indicated antibodies is shown. B. H&E staining of the indicated organotypic rafts. Thickness was quantitated from two independent rafts and the results shown below. C. Proteins were extracted from organotypic raft tissue generated with N/Tert-1+Vec cells, human foreskin keratinocytes immortalized by the entire HPV16 genome (HFK+HPV16) or human foreskin keratinocytes immortalized by the HPV16 E6 and E7 oncogenes (HFK+E6/E7). Western blotting with the indicated antibodies is shown. D. Proteins were extracted from W12 cells with episomal (W12 episomal) or integrated (W12 integrated) viral genomes just prior to differentiation induction (Day 0) and three days following calcium treatment (Day 3). Western blotting with the indicated antibodies is shown. E. Cells were stained with the indicated antibodies before and after 3 days of calcium induced differentiation. F. The results in Figure 1D were scanned and quantitated for the number of cells that had positive staining for E2 or γH2AX. G. Proximity ligation assays (PLA) were carried out for γH2AX-γH2AX interaction and for E2-γH2AX interaction before and after 3 days of calcium induced differentiation. H. The results in 1G were quantitated. * Indicates significant differences between samples, p-value<0.05. ** Indicates significant differences between samples, p-value<0.01.

We next investigated DDR activation in more biologically relevant models containing the entire HPV16 genome. Human foreskin keratinocytes immortalized with the entire HPV16 genome (therefore expressing E2; HFK+HPV16), or with the viral oncogenes E6 and E7 only (HFK+E6/E7), were cultured in 3D organotypic rafts to induce differentiation alongside N/Tert-1+Vec control cells. Figure 1B demonstrates that HFK+E6/E7 cells have a more disorganized and dysplastic phenotype than HFK+HPV16 with a reduced number of cell layers in the differentiated epithelium. There is expression of E2 and γH2AX throughout the differentiating epithelium in HFK+HPV16 (Prabhakar et al., 2021). Proteins were extracted and analyzed by western blotting for the “ATM up ATR down” phenotype. We observed elevated ATM, pATM, CHK2 and pCHK2 in differentiated HFK+HPV16 cells compared with the other cell lines (Figure 1C; compare lane 2 with lanes 1 and 3). ATR levels were also reduced in HFK+HPV16 cells compared to other samples, as were CHK1 and pCHK1 levels, thus repeating the “ATM up ATR down” pattern observed in N/Tert-1+E2-WT cells. The E6/E7 oncogenes expressed alone did not activate ATM and although ATR levels were increased, there was no downstream activation as levels of CHK1 and pCHK1 were comparable with N/Tert-1+Vec cells (compare lanes 1 and 3), indicating that the viral oncogenes do not activate the DDR during differentiation.

To further enhance biological relevance, we next investigated W12 cell clones for their ATM and ATR signaling dynamics; W12 cells were isolated from a pre-malignant HPV16 cervical lesion and cloned to generate lines with episomal (W12 episomal) or integrated (W12 integrated) viral genomes (Jeon et al., 1995; Stanley et al., 1989). Integration of the viral genome is common in cervical cancer and this results in loss of E2 protein expression as integration occurs in the E2 gene. W12 episomal (W12e, has E2 expression) and W12 integrated (W12i, has no E2 expression) cells were calcium differentiated and the status of ATM and ATR signaling determined. Following differentiation, the “ATM up ATR down” phenotype is observed in W12e cells (Figure 1D, compare lane 2 with the others); but not in of W12i cells. This again demonstrates that only in the presence of E2 is the “ATM up ATR down” pattern observed, and that the viral oncogenes do not substantially affect DDR during differentiation. E2 expression is only detectable in W12e cells following differentiation (compare lane 2 with lane 1), suggesting that it is stabilized during differentiation as it is during mitosis and in differentiating N/Tert-1 cells, Figure 1 (Prabhakar et al., 2024). All of the experiments in Figures 1A, C and D were repeated and quantitated and demonstrate significant changes in E2, ATM, pATM, ATR, CHK2, pCHK2, ATR, CHK1, and pCHK1 proteins following differentiation in all lines expressing E2-WT (Figure S1 A-C). These results establish E2 expression, but not E6 or E7 expression, as the key driver of the “ATM up ATR down” phenotype. None of the changes in the DDR proteins induced by E2 during differentiation were due to changes in RNA levels (Figure S1 D-F). Involucrin, a marker of differentiation, was induced in all of the calcium treated keratinocytes demonstrating an efficient induction of differentiation; western blots are shown in Figure 2 and RNA levels are shown in Figure S2.

**Figure 2.**
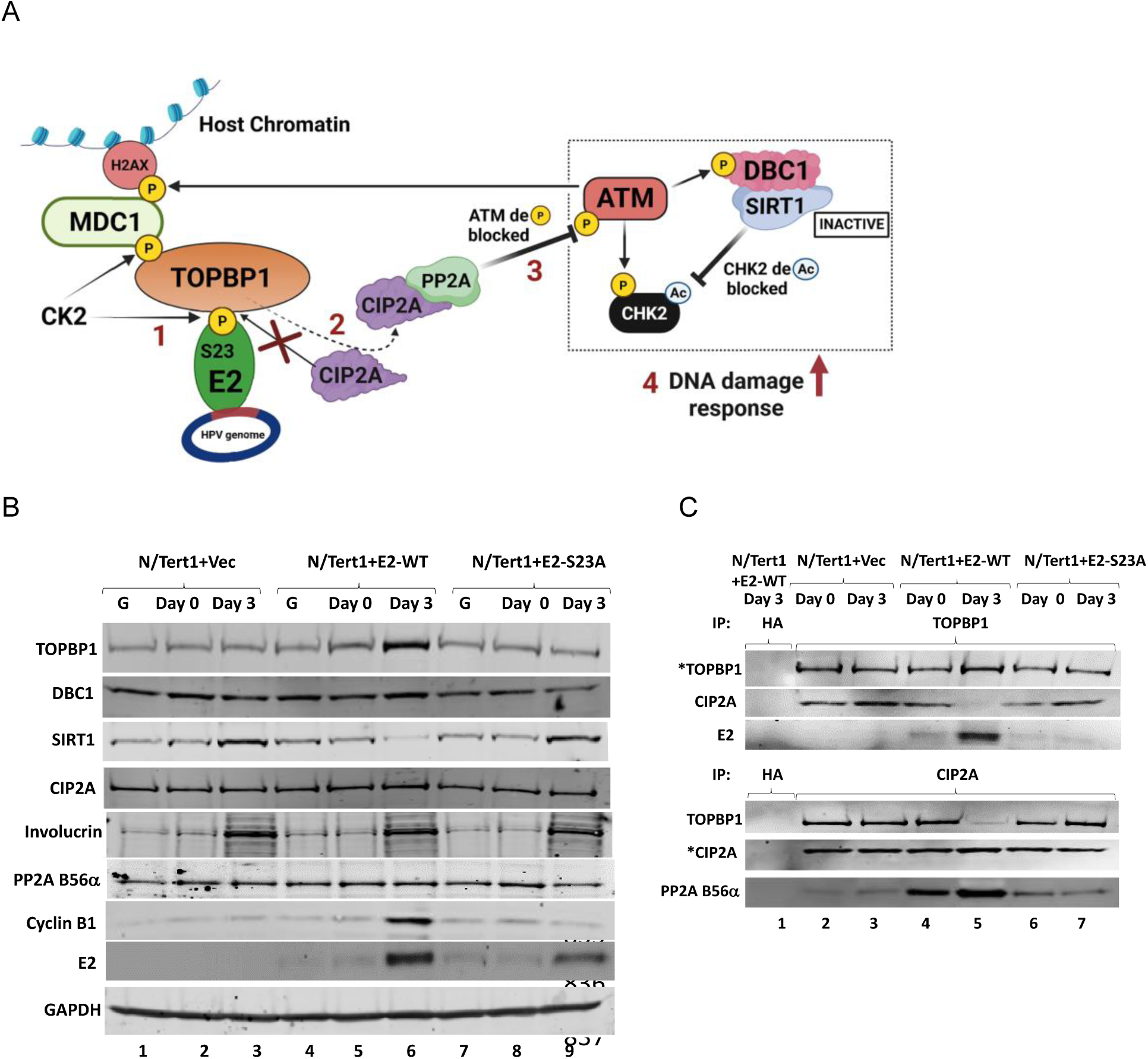
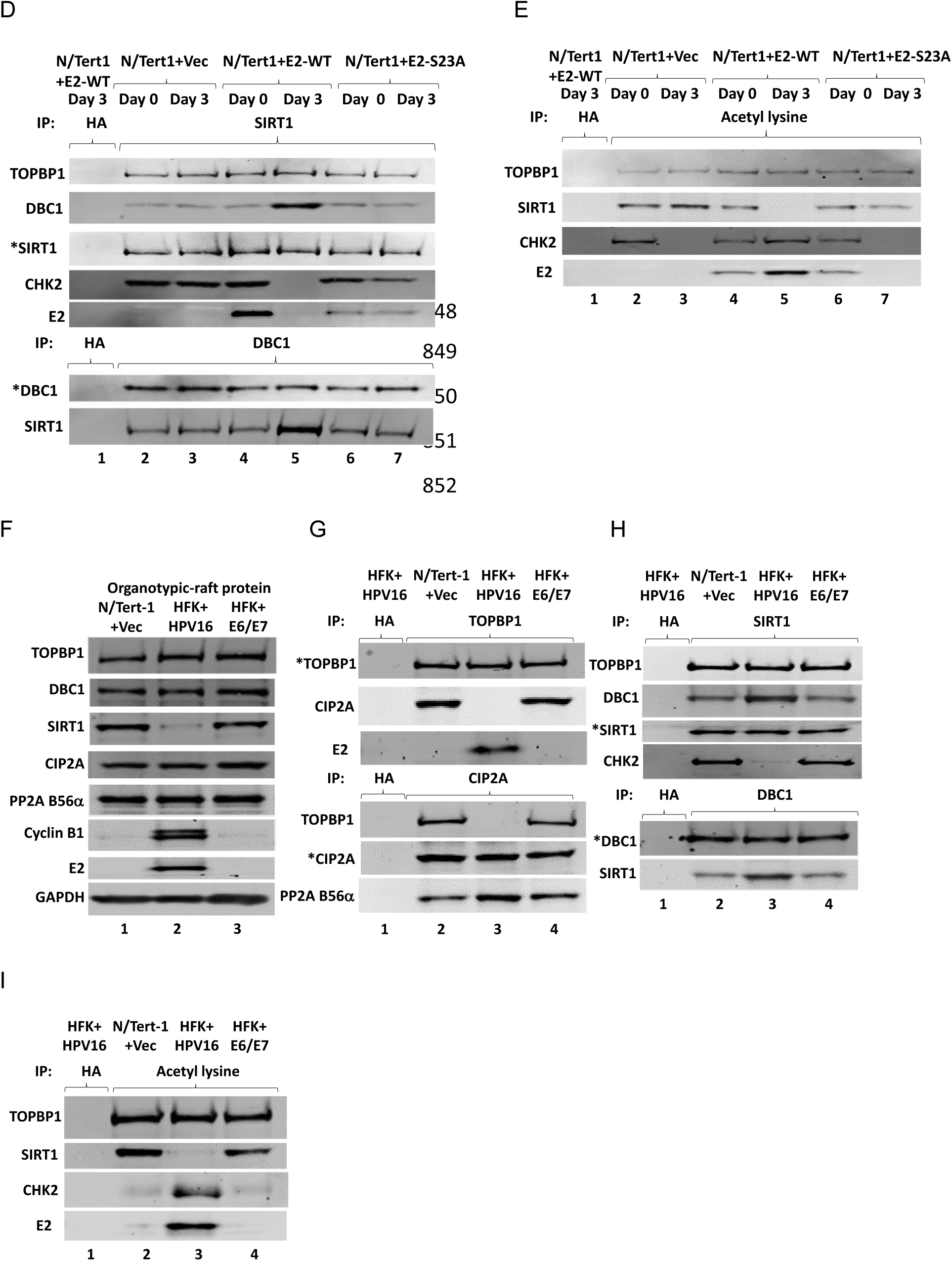
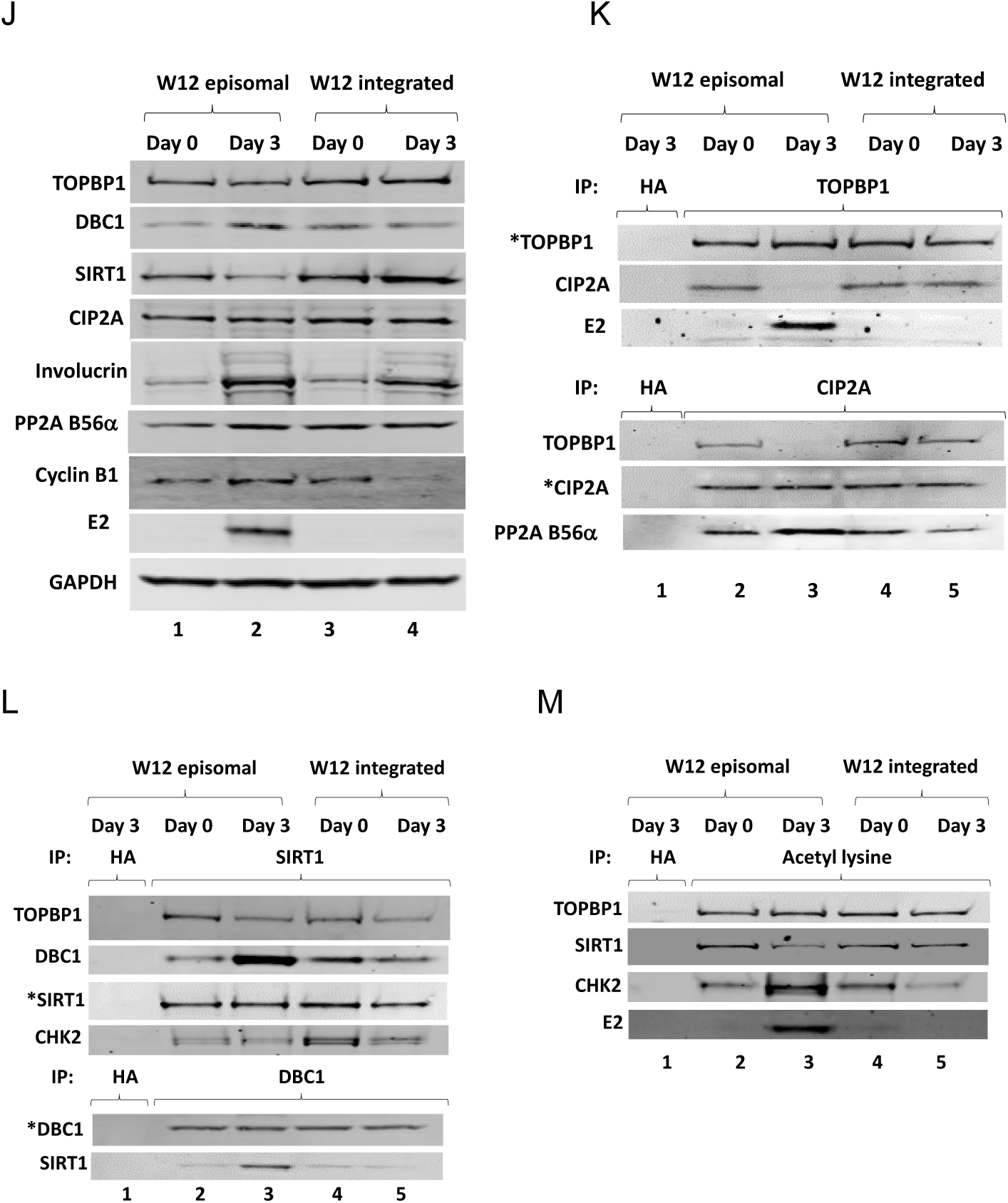
How E2 activates the DNA damage response during differentiation. A. A model that explains the biochemistry in Figure 2. In step 1 E2 complexes with TOPBP1 and removes CIP2A following differentiation. The “free” CIP2A has enhanced interaction with the PP2A subunit B56α, step 2, which inhibits PP2A function triggering an increased phosphorylation of ATM causing its activation and phosphorylation of CHK2, step 3. ATM activation enhances DBC1-SIRT1 interaction resulting in SIRT1 inhibition promoting increased acetylation of CHK2, a known activator of CHK2 activity, step 4. B. Proteins were extracted from N/Tert-1 cells with an empty vector control (N/Tert-1+Vec), expressing wild type E2 (N/Tert-1+E2-WT) or expressing an E2 that fails to interact with TOPBP1 (N/Tert-1+E2-S23A) that were growing (G), plated for differentiated with no calcium added (Day 0) or were induced to differentiate for 3 days following calcium treatment (Day 3). C. The protein extracts shown in B were immunoprecipitated with TOPBP1 (top panels) or CIP2A (lower panels) and western blotted for the proteins shown. D. The protein extracts prepared in B were immunoprecipitated with SIRT1 (top panels) or DBC1 (lower panels) and western blotted for the proteins shown. E. The protein extracts prepared in B were immunoprecipitated with an acetyl lysine specific antibody and western blotted for the proteins shown. F. Proteins were extracted from organotypic raft tissue generated with N/Tert-1+Vec cells, human foreskin keratinocytes immortalized by the entire HPV16 genome (HFK+HPV16) or human foreskin keratinocytes immortalized by the HPV16 E6 and E7 oncogenes (HFK+E6/E7). Western blotting with the indicated antibodies is shown. G. The protein extracts shown in F were immunoprecipitated with TOPBP1 (top panels) or CIP2A (lower panels) and western blotted for the proteins shown. H. The protein extracts shown in F were immunoprecipitated with SIRT1 (top panels) or DBC1 (lower panels) and western blotted for the proteins shown. I. The protein extracts shown in F were immunoprecipitated with an acetyl lysine specific antibody and western blotted for the proteins shown. J. Proteins were extracted from W12 cells with episomal (W12 episomal) or integrated (W12 integrated) viral genomes just prior to differentiation induction (Day 0) and three days following calcium treatment (Day 3). Western blotting with the indicated antibodies is shown. K. The protein extracts shown in J were immunoprecipitated with TOPBP1 (top panels) or CIP2A (lower panels) and western blotted for the proteins shown. L. The protein extracts shown in J were immunoprecipitated with SIRT1 (top panels) or DBC1 (lower panels) and western blotted for the proteins shown. M. The protein extracts shown in J were immunoprecipitated with an acetyl lysine specific antibody and western blotted for the proteins shown.

As E2 is responsible for activating the DDR during the HPV16 life cycle, we investigated co-localization of E2 with γH2AX before and after differentiation. N/Tert-1+Vec, HFK+HPV16 and HFK+E6/E7 were stained with E2 and γH2AX. Representative images are shown in Figure 1E and quantitated in Figure 1F. There is residual γH2AX staining in N/Tert-1+Vec cells that showed limited but significant increase following differentiation (Figure 1F, right hand panel). There is more γH2AX staining in HFK+HPV16 cells than in either of the other lines, and this significantly increases following differentiation. As expected, HFK+E6/E7 cells have enhanced damage when compared with N/Tert-1+Vec cells consistent with previous findings that the oncogenes induce DNA damage (Hoppe-Seyler et al., 2018), although the levels of γH2AX staining decreased in differentiated HFK+E6/E7 cells. In HFK+HPV16 cells, the levels of E2 also increased during differentiation (Figure 1F, left hand panel). Notably, undifferentiated HFK+HPV16 cells exhibit cytoplasmic γH2AX staining, suggesting the presence of damaged DNA in the cytoplasm (Figure 1E). However, after differentiation γH2AX is exclusively nuclear, where partial co-staining with E2 became apparent; this was not observed in undifferentiated cells. The staining patterns of E2 and γH2AX is relatively ubiquitous making it difficult to determine whether they are truly co-localized during differentiation; to investigate co-localization proximity ligation assays (PLA) were utilized (Figure 1G, quantitated in Figure 1H). In N/Tert-1+Vec cells γH2AX activity as measured using γH2AX-γH2AX PLA was low in undifferentiated cells and increased slightly, but significantly, following differentiation. HFK+HPV16 cells showed a marked and significant rise in γH2AX– γH2AX signal following differentiation, consistent with the E2 specific DDR activation observed in in Figure 1A, C and D. During differentiation there is a significant increase in the detection of nuclear E2-γH2AX foci as detected using PLA with around 2 puncta per cell detected. This demonstrates that E2 and γH2AX co-localize to only a small subset of foci in differentiating HFK+HPV16 cells. It is possible that differentiation induces a DNA double strand break at a specific chromosome site and E2 exploits this to interact with γH2AX and initiate DDR signaling.

### E2 displacement of CIP2A from TOPBP1 activates the DDR during the HPV16 life cycle

Having determined that the E2-TOPBP1 interaction is required for DDR activation during the HPV16 life cycle, we next investigated the mechanism. A clue was presented in two manuscripts about how fragmented human DNA can be segregated (Lin et al., 2023; Trivedi et al., 2023). These manuscripts demonstrated that fragmented damaged host DNA is recruited to mitotic host chromatin via a complex between TOPBP1 and cancerous inhibitor of protein phosphatase 2A (CIP2A) resulting in retention of the damaged DNA in a single daughter nucleus following cell division. CIP2A is overexpressed in a number of cancers and is a known inhibitor of PP2A (protein phosphatase 2A) (Chen et al., 2023). PP2A can dephosphorylate ATM to control activation of the DDR (Goodarzi et al., 2004). We tested a model shown in Figure 2A, where E2 displaces CIP2A from TOPBP1 during differentiation resulting in inhibition of PP2A, therefore promoting ATM activation. ATM is a known activator of DBC1, which is a SIRT1 inhibitor (Kim et al., 2008; Zannini et al., 2012; Zhao et al., 2008). We investigated whether SIRT1 is inactivated by E2 during differentiation, as it is during mitosis (Prabhakar et al., 2024). Figure 2B shows the levels of the proteins involved in the Figure 2A model in our N/Tert-1 cell lines, as well as cyclin B1 levels, with quantitated replicates (Figure S2A). Following differentiation TOPBP1, E2 and cyclin B1 levels are significantly increased in the presence of E2-WT protein (Figure 2B, lane 6) when compared with all other conditions; quantitation of repeat experiments is shown in Figure S2A. The increase in cyclin B1 is notable as, following differentiation, it has been proposed that viral replication occurs in a G2/M like environment (Davy and Doorbar, 2007). The levels of SIRT1 are reduced following differentiation of the E2-WT cells as it is during mitosis (Prabhakar et al., 2024). The ability of TOPBP1 to co-immunoprecipitate with CIP2A is attenuated following differentiation only in the presence of E2-WT (Figure 2C, upper panel, TOPBP1 IP, lower panel CIP2A IP, lane 5). At the same time there is an enhanced interaction between CIP2A and the PP2A subunit B56α, and between SIRT1 and DBC1 following differentiation of E2-WT cells (Figure 2D, lane 5). SIRT1 is a known interactor and regulator of E2 (Das et al., 2017; Prabhakar et al., 2024), and E2-SIRT1 interaction is blocked following differentiation (as it is in mitosis), Figure 2D lane 5, and the interaction between SIRT1 and CHK2 is also disrupted following differentiation of E2-WT cells. E2-WT and CHK2 protein levels increase following differentiation (Figure 1 and Figure 2B) and there is an enhanced acetylation of both proteins following differentiation that likely contributes to these increased protein levels (Figure 2E, lane 5). In E2-S23A cells (non-TOPBP1 binding), differentiation abrogates acetylation of both E2-S23A and CHK2, similar to the loss of CHK2 acetylation in N/Tert-1 control cells, and there is no disruption of the TOPBP-CIP2A interaction (lanes 2 and 7). CHK2 input levels are shown in Figure 1. All immunoprecipitations were repeated with multiple, independent, extracts and quantitated to determine significance (Figure S2B). Overall, the results support the model presented in Figure 2A; E2 displaces CIP2A from TOPBP1 during differentiation, promoting enhanced interaction between CIP2A and PP2A. This promotes ATM activity, leading to DBC1 phosphorylation and increased binding to, and inhibition of, SIRT1.

In proteins extracted from organotypic rafts of N/Tert-1+Vec, HFK+HPV16 and HFK+E6/E7 there is a significant reduction of SIRT1 protein levels (Figure 2F, lane 2) in the presence of E2 when compared with the N/Tert-1+Vec and HFK+E6/E7 cells, and this was repeated and quantitated (Figure S2C). Cyclin B1 levels were also significantly increased in the HFK+HPV16 sample when compared with the other non-E2 expressing cells (Figure 2F, lane 2), TOPBP1-CIP2A interaction was abrogated with enhanced CIP2A-B56α (a PP2A subunit) interaction (Figure 2G, lane 3), and SIRT1-DBC1 interaction (Figure 2H, lane 3). There is also disruption of SIRT1 interaction with E2 and CHK2 (Figure 2H, lane 3), accompanied by increased E2 and CHK2 acetylation in the presence of E2 (Figure 2I, compare lane 3 with lanes 2 and 4). All immunoprecipitations were repeated and quantitated (Figure S2D). The cell lines used for organotypic rafting were also differentiated with calcium, and that “ATM up ATR down” phenotype is observed in HFK+HPV16 but not in HFK+E6/E7 or N/Tert1+Vec (Figure S2E), as is the downregulation of SIRT1 protein expression (Figure S2F). The TOPBP1-CIP2A interaction is disrupted in calcium differentiated HFK+HPV16 cells (Figure S2G), while there is an enhanced CIP2A-B56α and DBC1-SIRT1 interaction. SIRT1 interaction with E2 and CHK2 is disrupted following differentiation (Figure S2H). The disruption of the SIRT1 interaction with E2 and CHK2 results in enhanced acetylation of E2 and CHK2 following differentiation of HFK+HPV16 cells (Figure S2I). These results reproduce the phenotype observed in differentiated N/Tert-1+E2-WT cells and in HFK+HPV16 organotypic rafts.

In W12e cells, differentiation results in a significant reduction in SIRT1 levels and a significant increase in Cyclin B1 and E2 levels (Figure 2J, lane 2), which was not detected in W12i cells. This was repeated and quantitated (Figure S2J). Following differentiation, the TOPBP1-CIP2A interaction is disrupted and the CIP2A-B56α(PP2A) interaction is enhanced (Figure 2K, lane 3), the DBC1-SIRT1 interaction is increased (Figure 2L, lane 3) and CHK2 and E2 acetylation enhanced (Figure 2M, compare lane 3 with lane 5) in differentiated W12e cells but not in W12i cells. All immunoprecipitations were repeated with multiple, independent extracts and quantitated (Figure S2K).

These findings demonstrate that our results obtained from multiple experiments with differentiated keratinocytes expressing wild type E2 protein support the model presented in Figure 2A. None of the changes in protein levels following calcium induced differentiation in all cell types (apart from involucrin induction) were due to RNA changes (Figures S2 L-N).

### CIP2A and DBC1 are required for E2-mediated DDR activation in differentiating keratinocytes

If the model in Figure 2A is correct, then down regulation of CIP2A (preventing PP2A inhibition) or DBC1 (preventing inhibition of SIRT1) would abolish E2 DDR activation. We first investigated whether E2 alters the cellular localization of CIP2A following differentiation. We performed immunofluorescence for B56α (PP2A) and CIP2A in N/Tert-1+Vec, HFK+E6/E7, and HFK+HPV16 cells under undifferentiated (Figure 3A) and calcium-differentiated conditions (Figure 3B). Following differentiation, the levels of nuclear CIP2A and B56α are significantly reduced in HFK+HPV16 cells (images were scanned and quantitated, Figure 3C). The total level of these proteins are not reduced by E2 (Figure 2), but the CIP2A fluorescent signal is reduced in the presence of E2; this may be due to a change in CIP2A protein conformation induced by the interaction of E2 with TOPBP1 resulting in a reduction of native protein detection via epitope masking. This result supports the model in Figure 2A where E2 displacement of CIP2A from TOPBP1 results in CIP2A localization to the cytoplasm along with B56α which would prevent B56α de-phosphorylating nuclear ATM therefore promoting DDR activation.

**Figure 3.**
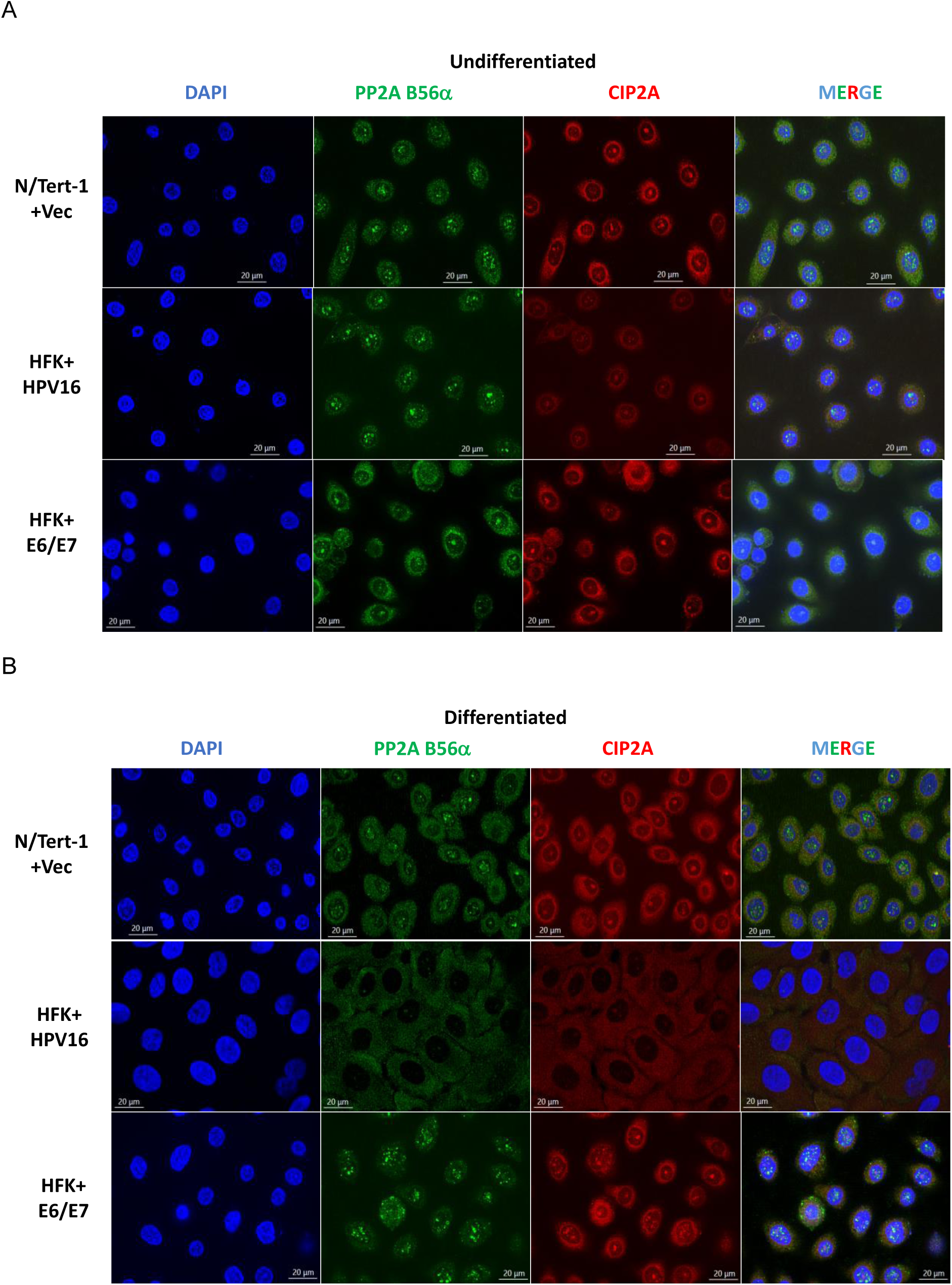
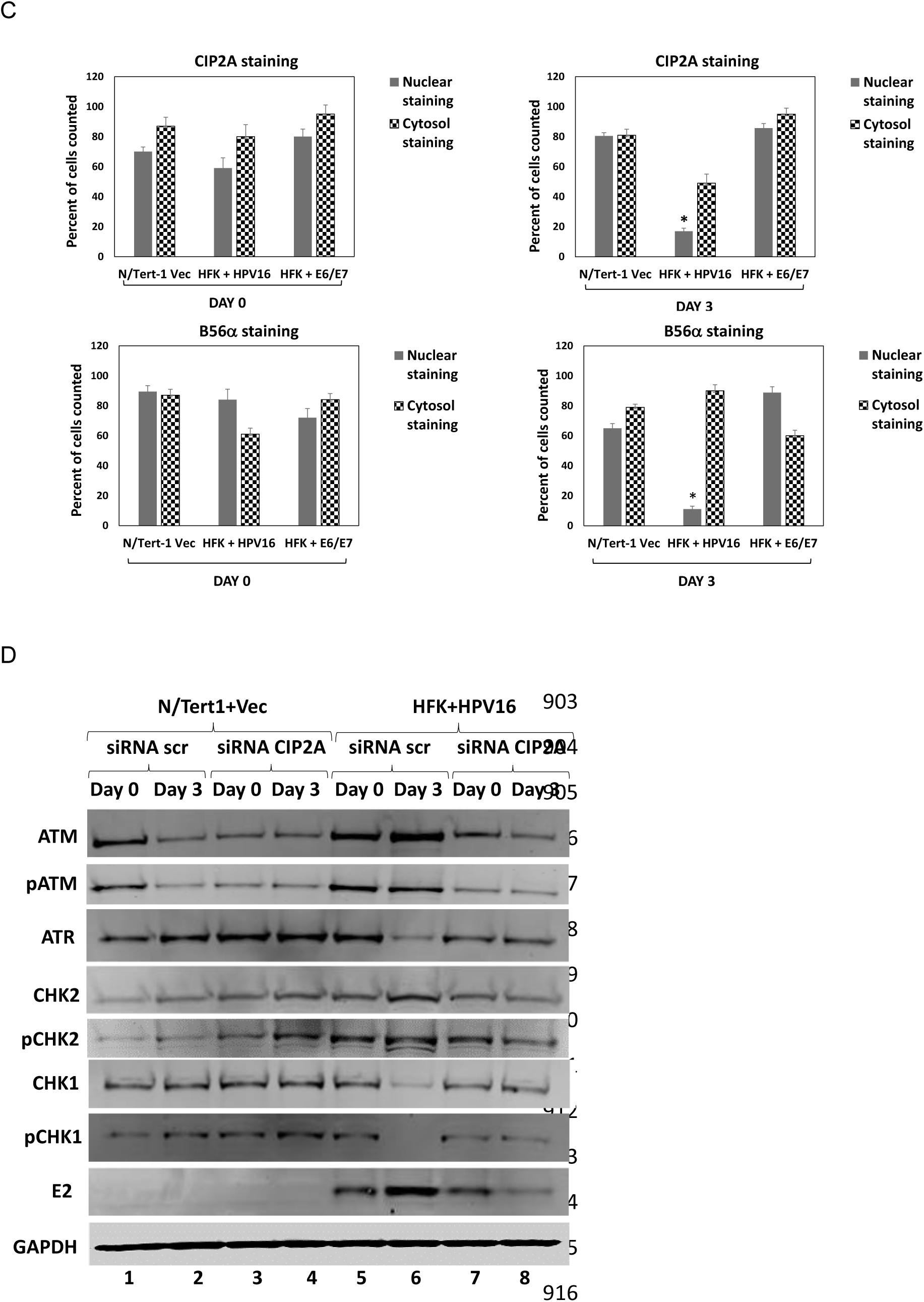
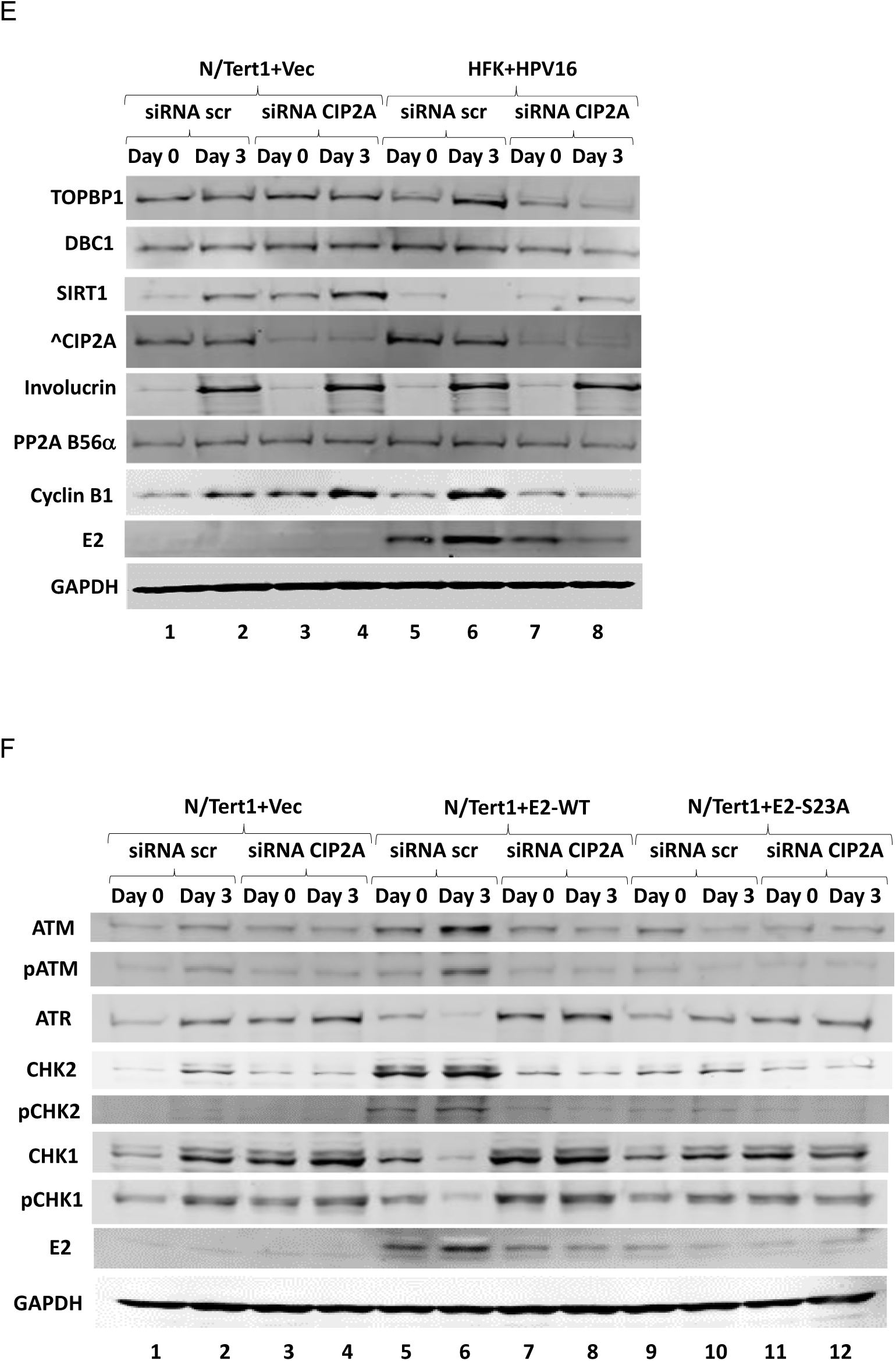
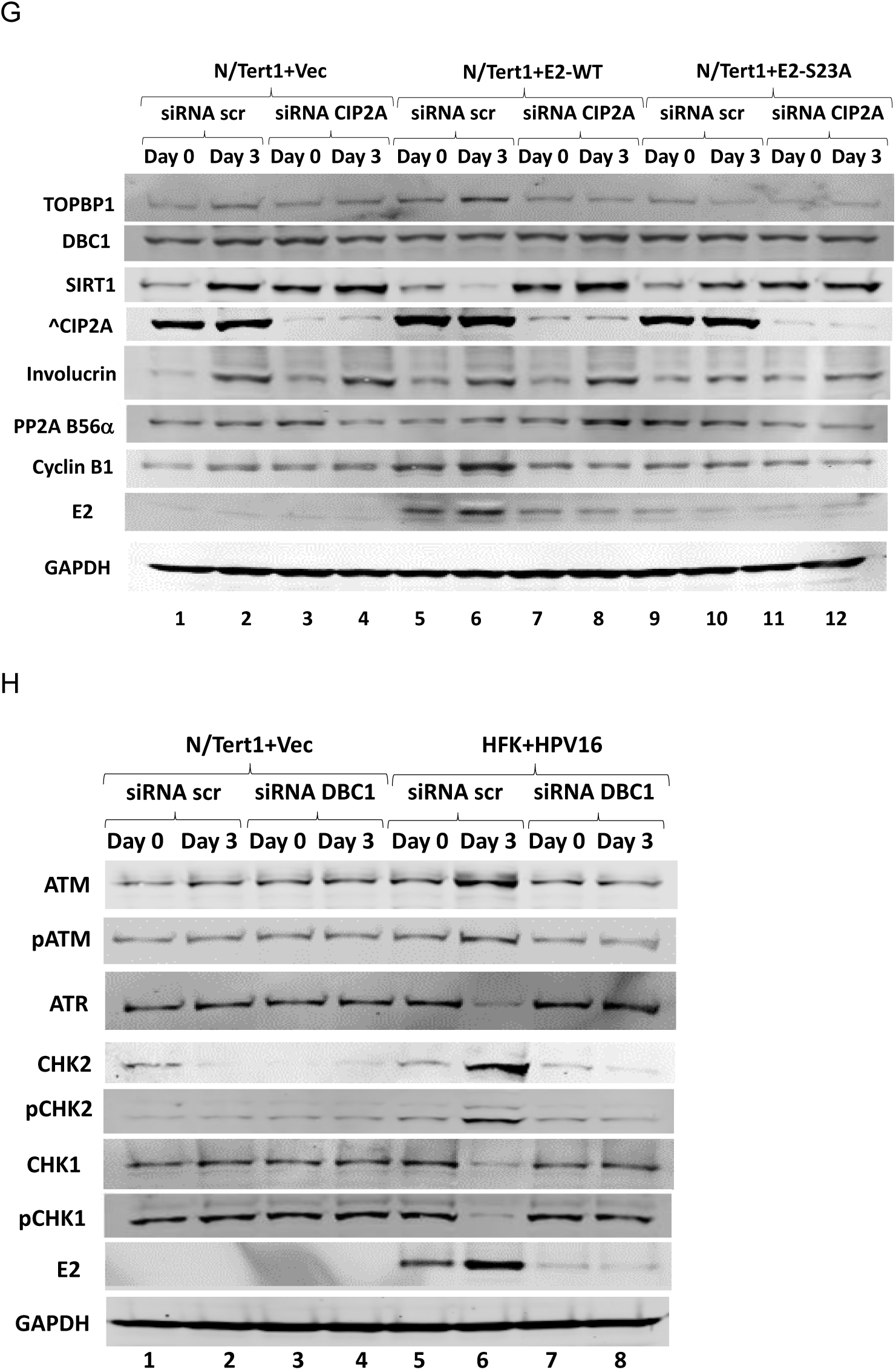
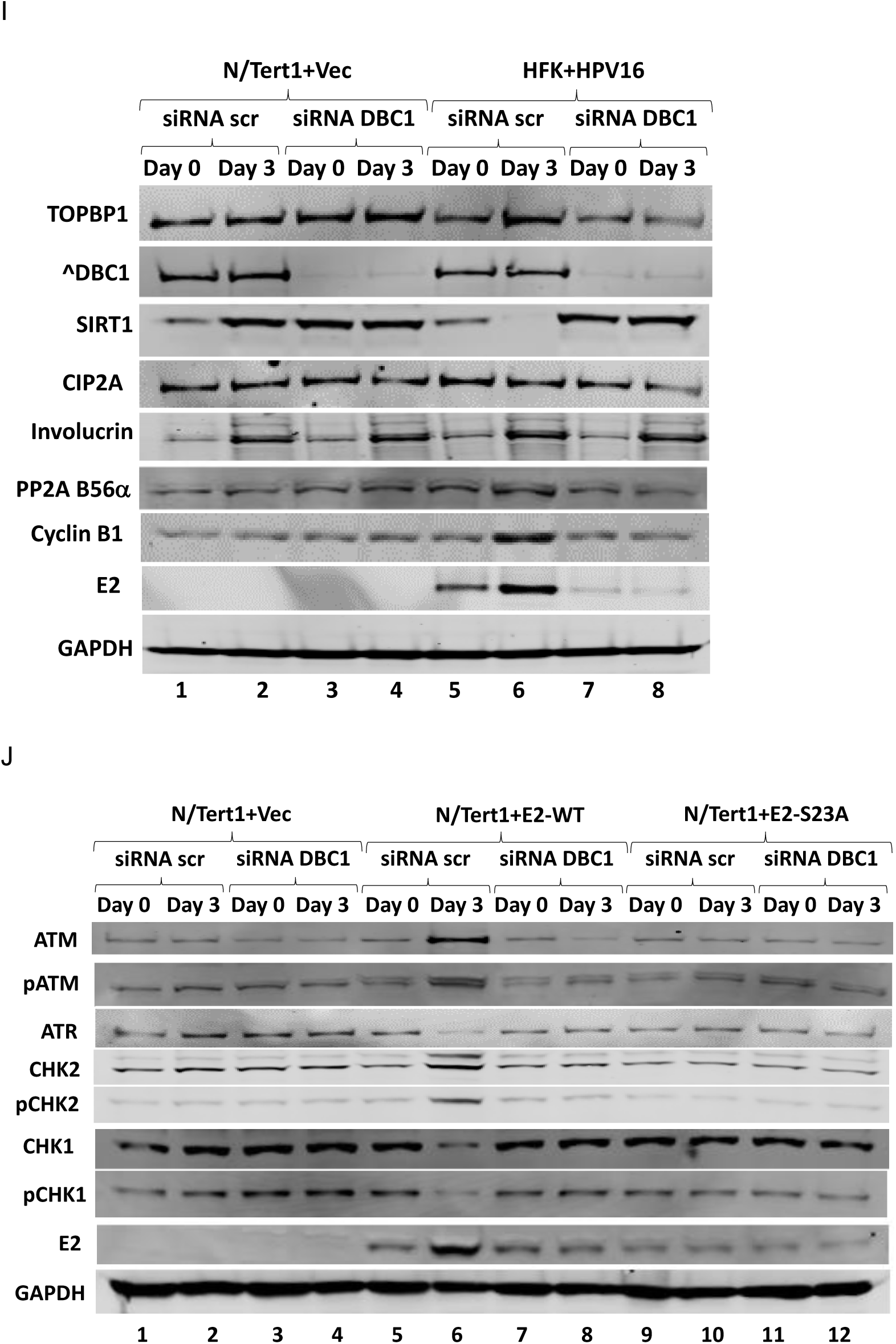
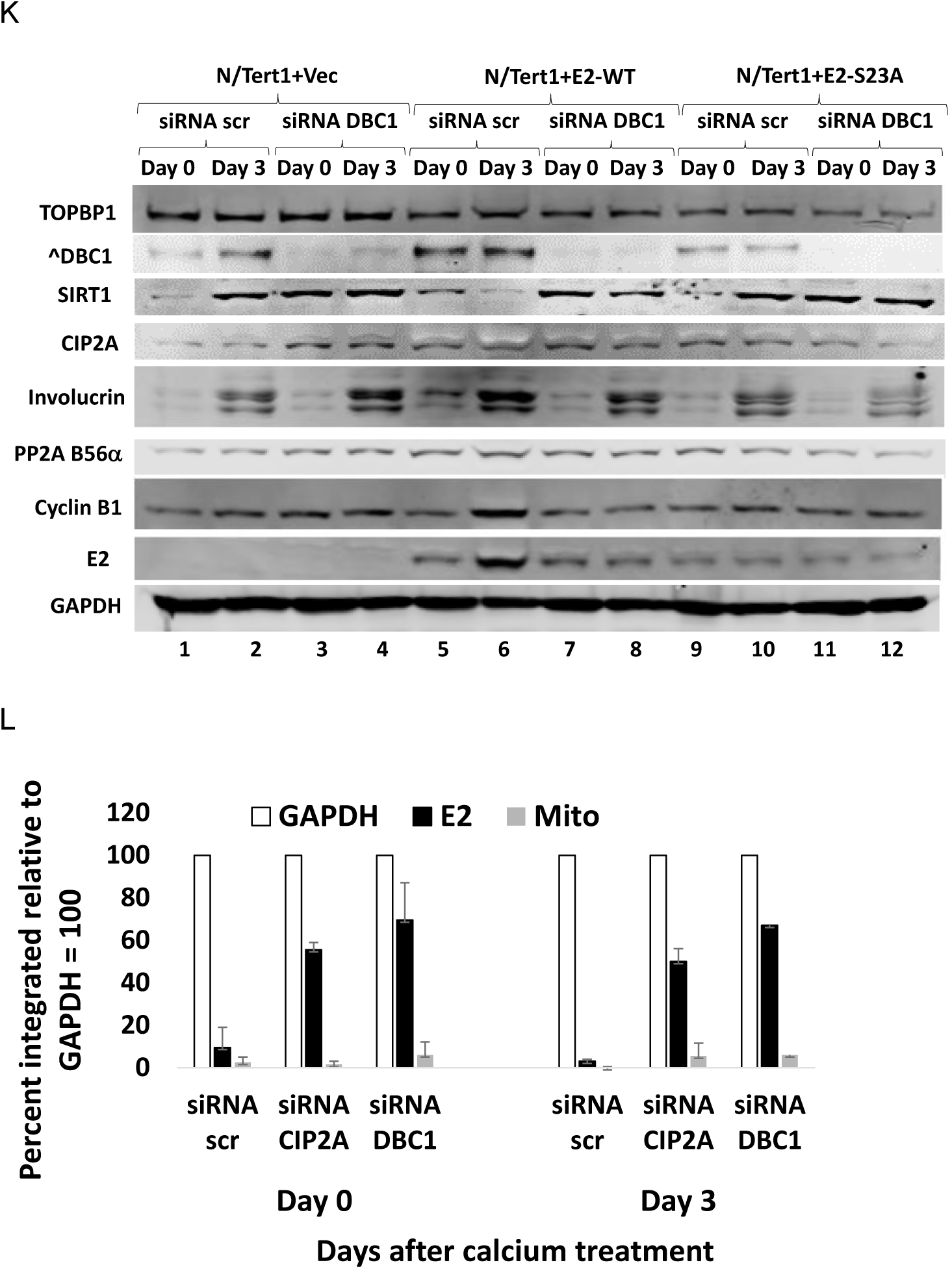
CIP2A and DBC1 are required for E2 activation of the DDR. A. Growing N/Tert-1+Vec (top panels), HFK+HPV16 (middle panels) and HFK+E6/E7 were stained with DAPI, B56α and CIP2A antibodies with a merge shown on the right panels. B. N/Tert-1+Vec (top panels), HFK+HPV16 (middle panels) and HFK+E6/E7 cells were differentiated by calcium treatment and three days later fixed and stained with DAPI, B56α and CIP2A antibodies with a merge shown on the right panels. C. Figures 3A and 3B are representative samples that were scanned and quantitated for the levels of nuclear and cytoplasmic B56α and CIP2A and the results are presented graphically. There is a statistically significant reduction in nuclear B56α and CIP2A following differentiation of HFK+HPV16 cells as indicated with *. D. siRNA knockdown of CIP2A prevents the “ATM up ATR down” phenotype in HFK+HPV16 cells following differentiation, and the increase in E2 protein expression. E. siRNA knockdown of CIP2A increases SIRT1 levels in HFK+HPV16 cells following differentiation but does not affect DBC1 or B56α expression and prevents the increase in cyclin B1 and TOPBP1. F. siRNA knockdown of CIP2A prevents the “ATM up ATR down” phenotype in N/Tert-1+E2-WT cells following differentiation, and the increase in E2 protein expression. G. siRNA knockdown of CIP2A increases SIRT1 levels in HFK+HPV16 cells following differentiation but does not affect DBC1 or B56α expression and prevents the increase in cyclin B1 and TOPBP1. H. siRNA knockdown of DBC1 prevents the “ATM up ATR down” phenotype in HFK+HPV16 cells following differentiation, and the increase in E2 protein expression. I. siRNA knockdown of DBC1 increases SIRT1 levels in HFK+HPV16 cells following differentiation but does not affect CIP2A or B56α expression and prevents the increase in cyclin B1 and TOPBP1. J. siRNA knockdown of DBC1 prevents the “ATM up ATR down” phenotype in N/Tert-1+E2-WT cells following differentiation, and the increase in E2 protein expression. K. siRNA knockdown of DBC1 increases SIRT1 levels in HFK+HPV16 cells following differentiation but does not affect CIP2A or B56α expression and prevents the increase in cyclin B1 and TOPBP1. L. TV exonuclease assays demonstrate that CIP2A and DBC1 knockdown promotes viral genome integration irrespective of the differentiation status of the cells. Each panel pair (D-E, F-G, H-I, J-K) in Figure 3 was generated from the same blot, stripped and re-probed for different targets. E2 and GAPDH controls are re-used within each pair.

To determine whether CIP2A is required for E2-mediated DDR activation, N/Tert-1+Vec and HFK+HPV16 cells were treated with CIP2A siRNA. Figure 3D demonstrates that treatment of HFK+HPV16 with scrambled control siRNA (siRNA scr) does not affect the “ATM up ATR down” phenotype (lane 6). There are increased levels of ATM, CHK2 and pCHK2 with reduced ATR, CHK1 and pCHK1 levels. However, following treatment of cells with siRNA targeting CIP2A there is an abrogation of these changes. Figure 3E confirms the knockdown of CIP2A expression (lanes 3, 4, 7 and 8), which prevented the reduction in SIRT expression, and induction of E2 expression, by HFK+HPV16 following differentiation. These experiments were repeated in N/Tert-1+Vec, N/Tert-1+E2-WT and N/Tert-1+E2-S23A with similar results (Figures 3F and 3G). siRNA scr had no effect on the “ATM up ATR down” phenotype in E2-WT cells (Figure 3F, lane 6), while treatment with siRNA targeting CIP2A abolished this phenotype (Figure 3F, lane 8). CIP2A knockdown also blocked downregulation of SIRT1 by E2-WT (Figure 3G, compare lane 8 with lane 6) and the induction of E2 protein levels following differentiation. These experiments were repeated with an additional CIP2A siRNA with similar results and quantitated for both siRNAs to demonstrate the significance of the results (Figures S3A-S3H). RNA levels were also determined for one of the siRNA treatments carried out in duplicate and no changes in RNA levels, outside of CIP2A, were observed for the proteins under study (Figures S3I and S3J).

Similarly to CIP2A knockdown, DBC1 knockdown also abolished E2-mediated DDR activation during differentiation (Figures 3H-3K), the results were reproducible with an alternative DBC1 siRNA confirming significance (Figures S3K to S3R) and DBC1 downregulation did not change the RNA expression of any of the proteins under study outside of DBC1 (Figures S3S and S3T).

Collectively, the results from Figures 2 and 3 support the model in which E2 disruption of the TOPBP1-CIP2A interaction during differentiation initiates a cascade that ultimately activates the DDR. Downregulation of either CIP2A or DBC1 enhanced viral genome integration before and after differentiation emphasizing the critical role for these factors in controlling the HPV16 life cycle (Figure 3L).

### The E2-TOPBP1 interaction promotes complex formation with ATM following differentiation and sensitizes cells to ATM inhibition

Following DNA damage, TOPBP1 association promotes ATR activation resulting in CHK1 phosphorylation (Kumagai et al., 2006). The activation of ATR by TOPBP1 requires the formation of TOPBP1 condensates and such condensates are recognized regulators of various cellular processes, including cell signaling (Egger et al., 2024; Frattini et al., 2021; Hadarovich et al., 2025; Qin and Shi, 2024; Velichko et al., 2025). It has been proposed that ATR activation can occur via ATM phosphorylation of TOPBP1, therefore the failure of ATR activation following DDR induction during the differentiation of E2 containing keratinocytes could be due to a failure of TOPBP1 to interact with ATR in the presence of E2 (Yoo et al., 2007). Using the cell extracts described in Figures 1-2, TOPBP1 immunoprecipitations revealed formation of a TOPBP1-ATM complex following differentiation of E2 containing keratinocytes, with a failure of TOPBP1 to complex with ATR (Figure 4A). The top panels are the N/Tert-1+E2 cells, the middle panels the HFK, and the bottom panels W12 cells. In all cases, following differentiation, there is a strong interaction between TOPBP1 and ATM and a reduced interaction with ATR in cell lines expressing E2-WT (lane 5). In all other samples there is an interaction between TOPBP1 and ATR, demonstrating that E2 disrupts the TOPBP1-ATR interaction and promotes the TOPBP1-ATM interaction only following differentiation. Immunoprecipitation with an E2 antibody demonstrates that E2 can complex with both ATM and ATR (as well as TOPBP1) in non-differentiated keratinocytes expressing E2-WT (Figure 4B, lane 4). The non-TOPBP1 binding E2-S23A mutant did not complex with either ATM or ATR suggesting that the ability of E2 to complex with ATM and/or ATR is mediated via an E2-TOPBP1 interaction (lanes 6 and 7). In E2-WT cells induced to differentiate E2 retains the ability to complex with TOPBP1 and ATM but fails to interact with ATR (Figure 4B, lane 5). The immunoprecipitations were repeated with different protein extracts and quantitated to demonstrate the significance of the results (Figures S4A and S4B).

**Figure 4.**
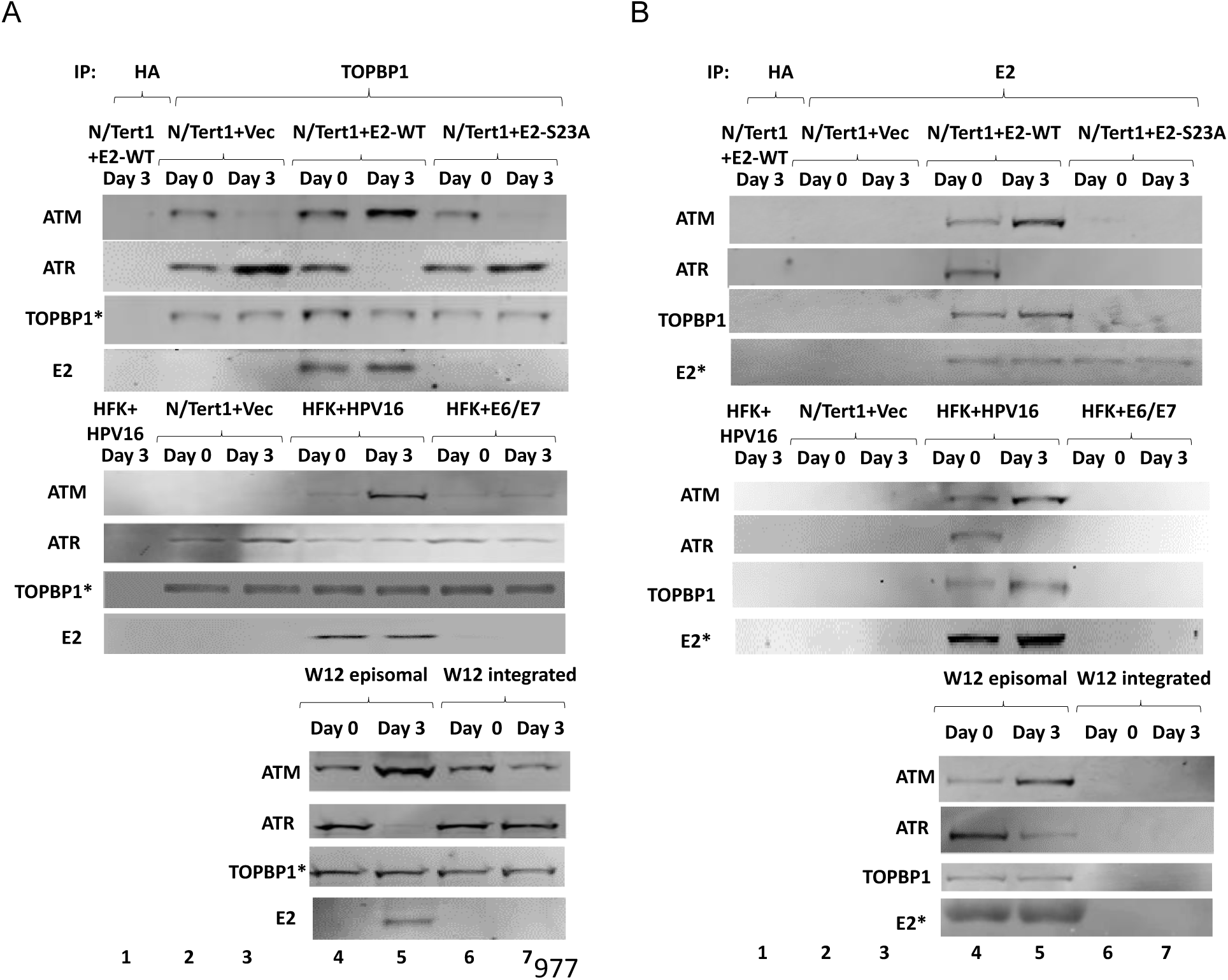
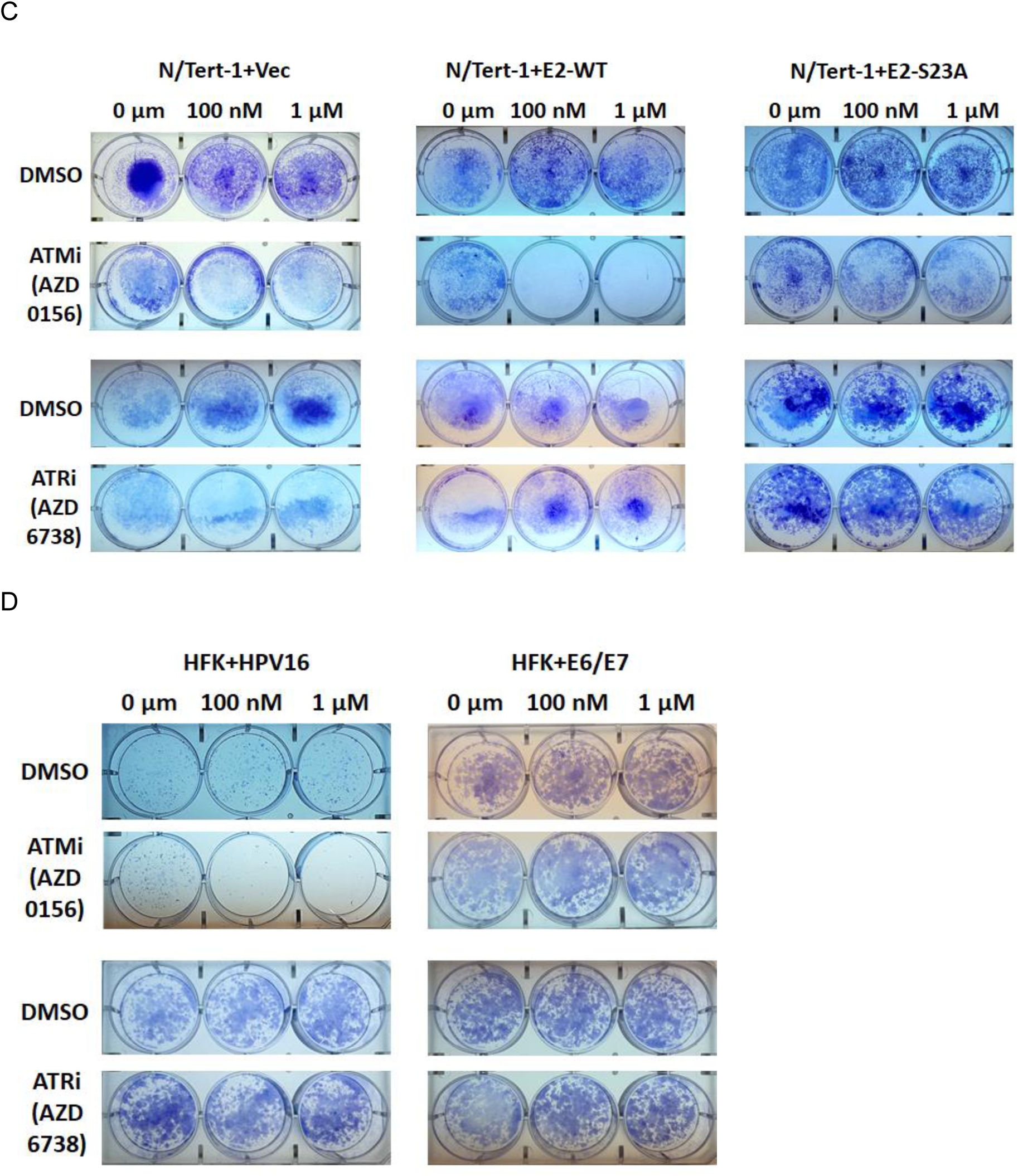
E2 promotes TOPBP1-ATM complex formation during differentiation and prevents TOPBP1 interaction with ATR. A. Immunoprecipitation with a TOPBP1 antibody in the presence of wild type E2 demonstrates a preferential formation of a TOPBP1-ATM complex following differentiation. The top panels show the results in N/Tert-1 cell lines, the middle panels in the HFK cell lines, and the bottom panels the W12 cell lines. B. Immunoprecipitation with an E2 antibody demonstrates that, following differentiation, wild type E2 complexes with ATM and TOPBP1 but not ATR. This phenotype depends upon E2 interaction with TOPBP1 as E2-S23A cannot interact with ATM or ATR. The top panels show the results in N/Tert-1 cell lines, the middle panels in the HFK cell lines, and the bottom panels the W12 cell lines. The input protein levels for these experiments are shown in Figure 1. C. The N/Tert-1 cell lines were treated with the ATM inhibitor AZD0156 (ATMi), top two panels, or the ATR inhibitor AZD6738 (ATRi), bottom two panels. Cells were treated with inhibitor for 48 hours and then seeded for colony formation. Colonies were fixed and stained with crystal violet after 10-14 days. D. The HFK cell lines were treated with the ATM inhibitor AZD0156 (ATMi), top two panels, or the ATR inhibitor AZD6738 (ATRi), bottom two panels. Cells were treated with inhibitor for 48 hours and then seeded for colony formation. Colonies were fixed and stained with crystal violet after 10-14 days.

The “ATM up ATR down” phenotype has the potential to sensitize cells to either ATM inhibition (as the cells have become addicted to ATM) or ATR inhibition (targeting the residual ATR activity could kill cells preferentially). To test these possibilities, we inhibited ATM and ATR in E2-expressing cells which we have shown activate DDR during mitosis, potentially creating a vulnerability to ATM or ATR inhibition (Prabhakar et al., 2024). We found that inhibition of ATM using AZD0156 preferentially killed N/Tert-1+E2-WT cells with no effect on N/Tert-1+Vec or N/Tert-1+E2-S23A cells (Figure 4C; upper two panels). Treatment with the ATR inhibitor AZD6738 had no effect on any of the cell lines. HFK+HPV16 and HFK+E6/E7 cells were treated with AZD0156 or AZD6738 and the HFK+HPV16 cells, that contain E2, were preferentially sensitive to the ATM inhibitor (Figure 4D). These results demonstrate that E2 mitotic DDR activation creates a vulnerability to ATM inhibition. The cell killing assays were repeated and quantitated (Figure S4C).

### The mechanism of E2-mediated DDR activation persists in papillomavirus infected tissue

Papillomaviruses infect and cause disease in differentiating epithelium. To investigate whether the E2 mechanism of DDR activation persists in an *in vivo* papillomavirus infection model, we evaluated murine tissues infected with MmuPV1 (Mus musculus papillomavirus 1). MmuPV1 causes disease in multiple anatomical sites including the mouse female reproductive tract, FRT (Spurgeon and Lambert, 2020; Spurgeon et al., 2019). For the sake of clarity, the letter m will be attached to all murine protein names (e.g., mE2 for murine E2). To confirm that mE2 interacts with mTOPBP1 via serine 23 as with HPV16 E2, stable NIH-3T3 cell lines were generated expressing mE2-WT and mE2-S23A (Figure 5A). Expression of mE2-WT (lane 2) or mE2-S23A (lane 3) did not change mTOPBP1 levels (Figure 5A, compare lanes 2 and 3 with lane 1). Immunoprecipitation of mTOPBP1 co-precipitated mE2-WT and not mE2-S23A (Figure 5B, compare lanes 3 and 4), confirming that mE2 interacts with mTOPBP1 via the same residue HPV16 E2 does (Prabhakar et al., 2021). Figure 5C demonstrates induction of dysplasia in the FRT of MmuPV1 infected mice and not in mock infected mice. Using FRT tissue from non-infected and MmuPV1-infected mice, proteins were extracted and mATM and mATR signaling investigated (Figure 5D). The infected FRT tissue had increased mATM, mpATM, mCHK2 and mpCHK2 and a reduced level of mATR, mCHK1 and mpCHK1 when compared with the non-infected control tissue (compare lane 2 with lane 1). This experiment was repeated on an additional FRT infected sample that exhibited an identical “ATM up ATR down” phenotype (Figure S5A). Quantitation demonstrated a statistically significant increase in mATM, mpATM, mCHK2 and mpCHK2 and reduced mATR, mCHK1 and mpCHK1 in infected tissue versus control tissue (Figure S5B). Figure 5E shows the expression levels of the mouse proteins involved in the model proposed and described in Figure 2 for E2 activation of the DDR. Unlike in the models in Figure 2, there is no significant change in SIRT1 protein levels in the infected versus non-infected FRT tissue (compare lane 1 with lane 2 in Figure 5E). This was repeated in an additional FRT tissue and the cumulative results quantitated (Figures S5C and S5D). The interaction between mTOPBP1 and mCIP2A was disrupted in MmuPV1 infected tissue versus control with an increase in mCIP2A-mB56α interaction (Figure 5F, compare lane 3 with lane 2), and there was an enhanced interaction between mSIRT1 and mDBC1 (lane 3, Figure 5G). Figure 5H demonstrates an enhanced acetylation of mTOPBP1 and mCHK2 in the lesion tissue (lane 3), with a reduction in mSIRT1 acetylation. mE2 is also acetylated. This is the same pattern observed in our human pre-cancerous model cell lines described in Figures 1-3. These experiments were repeated in an additional lesion tissue with comparable results (Figure S5E-G). The immunoprecipitations were repeated and quantitated to demonstrate the significance of the results (Figure S5H). DNA was extracted from the lesions and TV exonuclease assays confirmed the presence of 100% episomal MmuPV1 genomes in these lesions (Figure S5I). In MmuPV1 muzzle lesions the viral genome is predominantly integrated, indicating there may be distinct differences between MmuPV1 genome status in different anatomical sites of infection (Yu et al., 2021). The results from the MmuPV1 FRT infected tissue demonstrate that the “ATM up ATR down” phenotype persists *in vivo* and that this is likely due to E2 disruption of the mTOPBP1-mCIP2A interaction. This is similar to the phenotypes observed in differentiating human cells *in vitro* (Figures 1-2).

**Figure 5.**
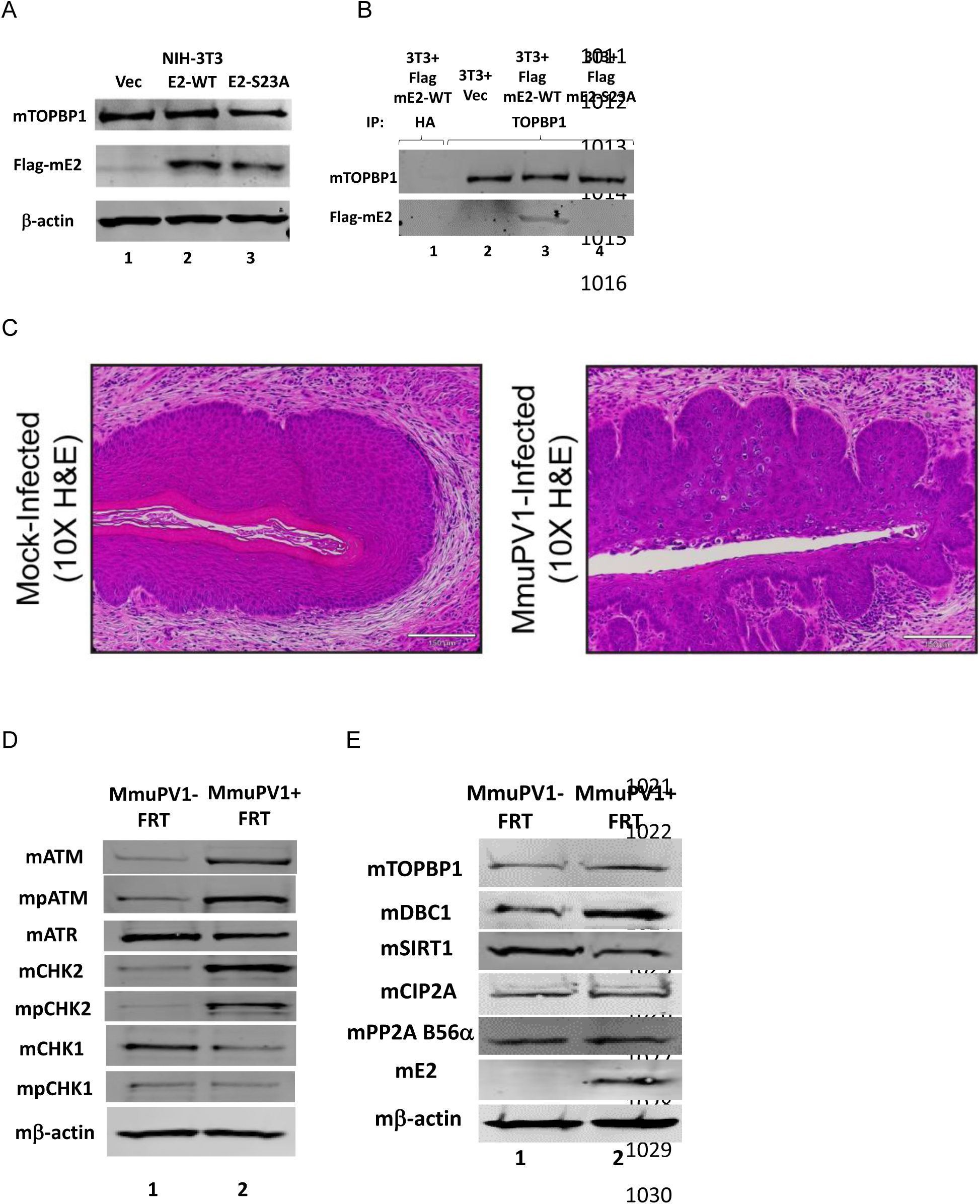
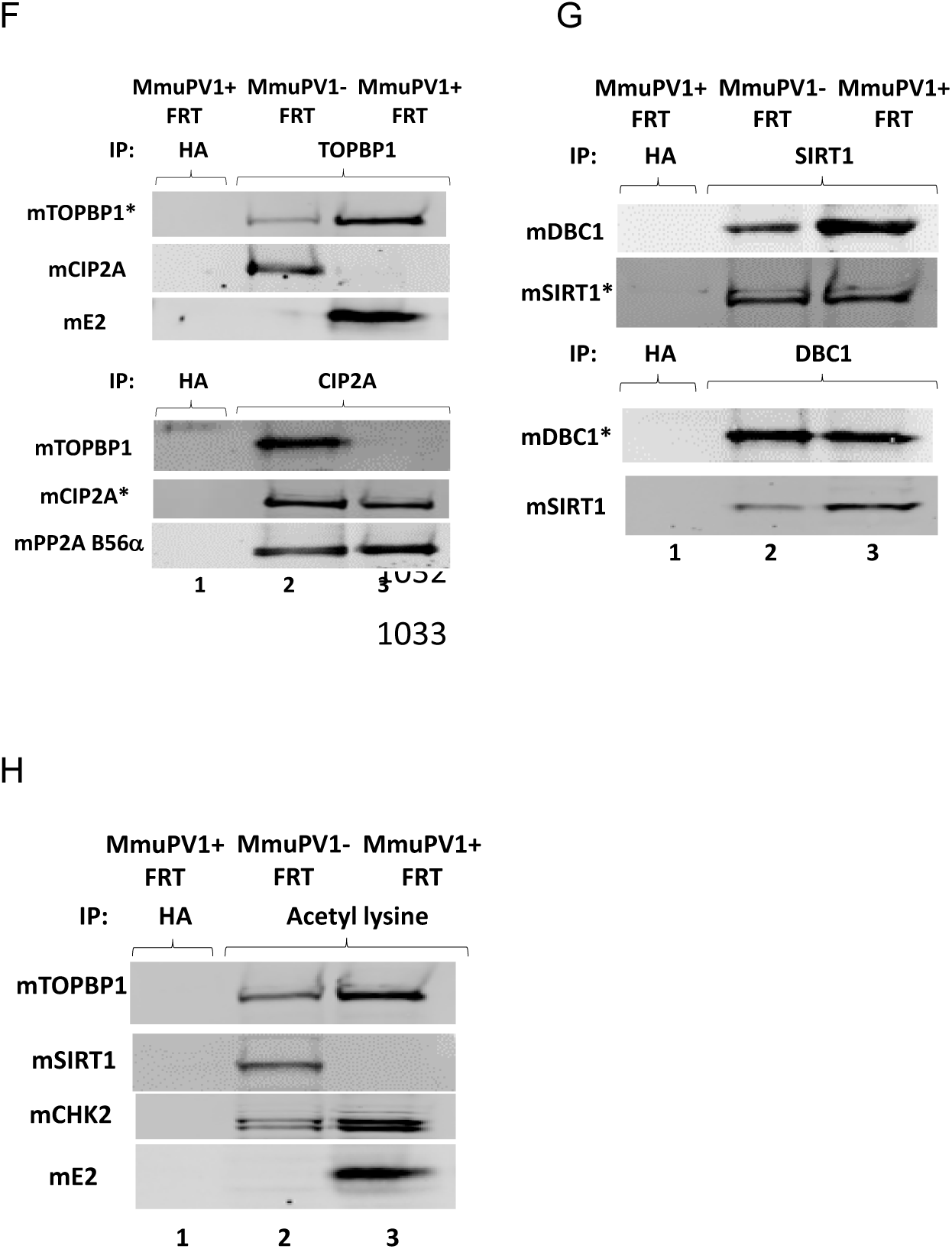
MmuPV1 lesions retain the E2 mechanism of DDR activation. A. NIH-3T3 cells were generated stably expressing empty vector, wild-type mE2 (E2-WT) or mE2-S23A (serine 23 mutated to alanine). B. The cell extracts from A were immunoprecipitated with a TOPBP1 antibody and the results demonstrate that mE2-WT can interact with mTOPBP1, but mE2-S23A cannot. C. Tissues from mock (left) and MmuPV1 (right) infected tissue were fixed and H&E stained. MmuPV1 dysplasia is evident in the disorganized epithelium with an increase in koilocytes. D and E. Protein was extracted from the female reproductive tract (FRT) of non-infected (MmuPV1-) or infected (MmuPV1+) mice and western blotting carried out with the indicated antibodies. F-H.

### The mechanism of E2-mediated DDR activation persists in HPV16 cancers

HPV16 cancers originate from infected, differentiating epithelium that continue to proliferate rather than terminally differentiate (Schiffman et al., 2016). As E2 activates the DDR during differentiation, we investigated whether the E2 induced DDR persists in HPV16 cancers that avoid terminal differentiation due to persistent proliferation. The majority of HPV16+HNSCC (head and neck squamous cell carcinomas) retain episomal viral genomes, indicating they likely retain E2 expression which is required for episomal maintenance (Nulton et al., 2018; Nulton et al., 2017). UMSCC-104 cells have episomal genomes and they express the E2 protein to maintain these genomes (Witt, 2025). UMSCC-47 is an example of an integrated cell line in which E2 expression has been lost. UMSCC-104 cells have an “ATM up ATR down” phenotype as there is increased ATM, pATM, CHK2 and pCHK2 and reduced ATR, CHK1 and pCHK1 when compared with UMSCC-47 (Figure 6A, compare lanes 1 and 2). This was repeated and quantitated to demonstrate significant differences in ATM and ATR signaling pathways between the samples (Figure S6A). The disruption of the TOPBP1-CIP2A interaction was investigated, Figure 6B shows the protein levels of the complex. As with the mouse FRT MmuPV1 lesions, the levels of SIRT1 (or any of the other complex proteins) are not different between UMSCC-104 and UMSCC-47 cells; this was repeated and quantitated (Figure S6B). In UMSCC-104 cells (which retain E2 expression) the TOPBP1-CIP2A interaction is disrupted with an enhanced CIP2A-B56α and DBC1-SIRT1 interaction with increased CHK2 and E2 acetylation compared to UMSCC-47 cells (Figure 6C-6E, compare lanes 2 and 3). The increased CHK2 acetylation in UMSCC-104 cells is likely due to a decreased interaction with SIRT1 (Figure 6D). Although UMSCC-104 and UMSCC-47 cells express comparable levels of SIRT1 (Figure 6B), SIRT1 is significantly more acetylated in UMSCC-47 cells (Figure 6E) suggesting that SIRT1’s acetylation status may regulate its enzyme function. In UMSCC-104 TOPBP1 complexes with ATM and E2 but not ATR, and in UMSCC-47 cells TOPBP1 can interact with both ATM and ATR, explaining the differential regulation of these two kinases between the two cell lines (Figure 6F). The immunoprecipitation experiments were repeated with additional cell extracts and quantitated (Figure S6C). When cells were treated with the ATM inhibitor AZD0156, UMSCC-104 are preferentially sensitive compared with UMSCC-47 cells (Figure 6G). This experiment was repeated and quantitated (Figure S6D). This data demonstrates that the model for DDR activation by E2 described in Figure 2 persists in HPV16 positive cancers.

**Figure 6.**
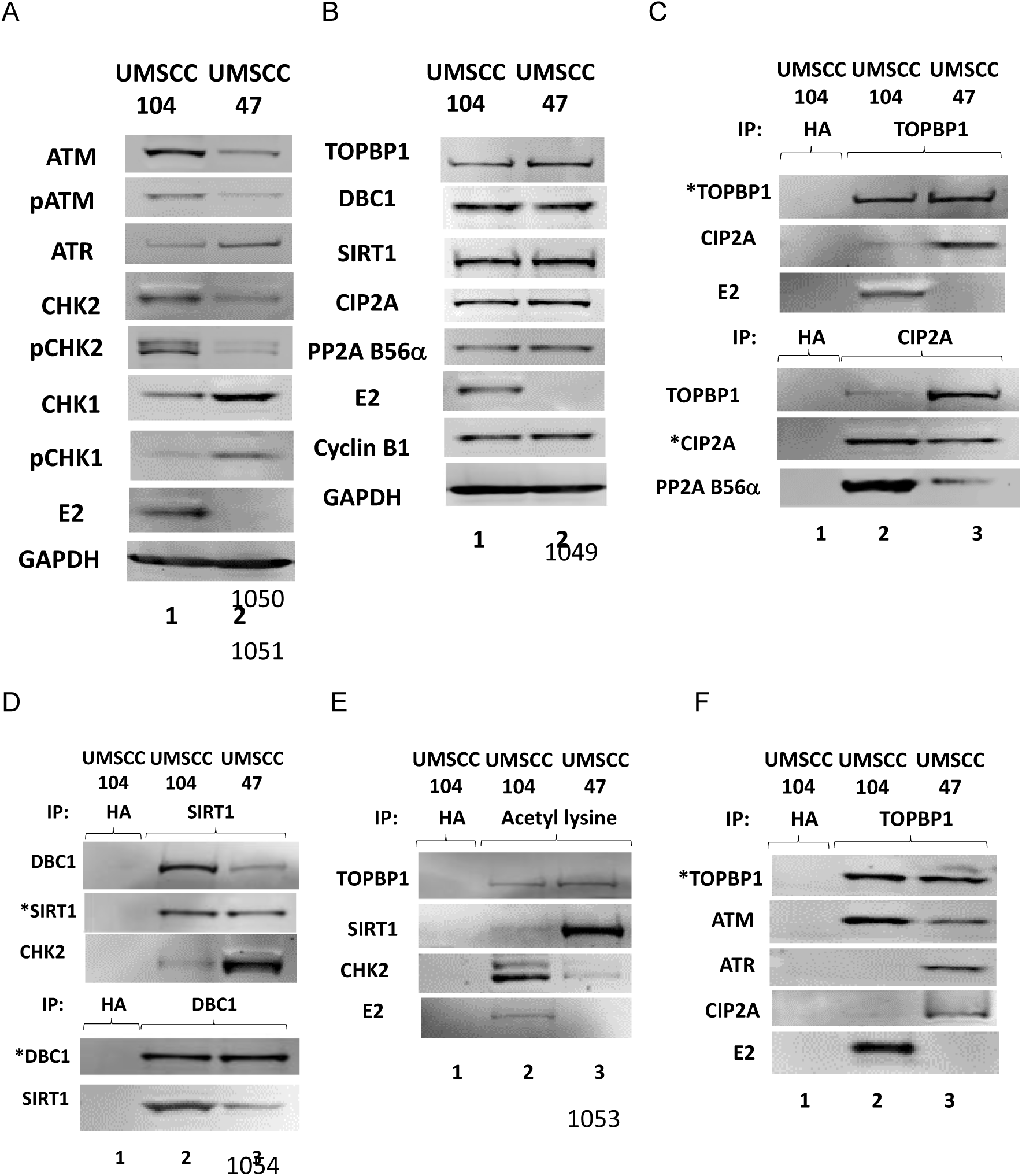
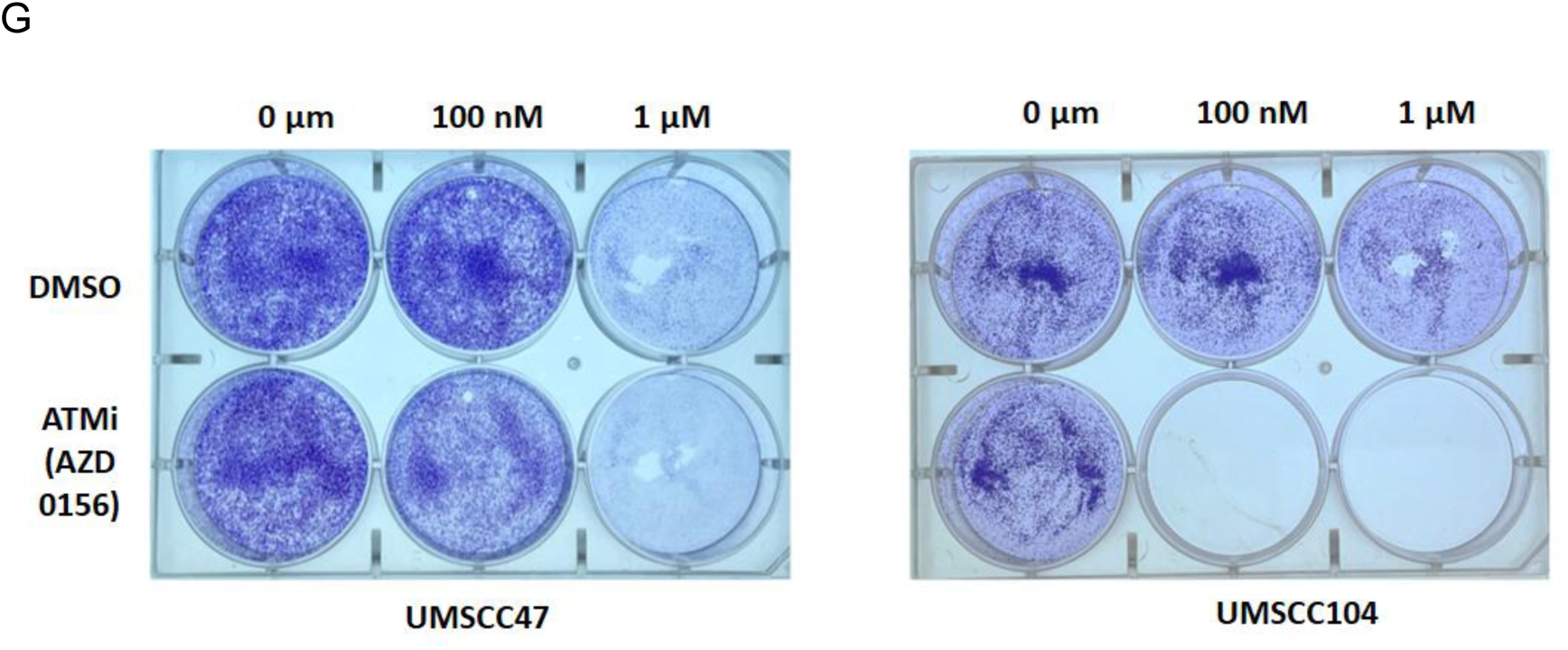
The “ATM up ATR down” phenotype persists in an HPV16 positive head and neck cancer cell line. A and B. Proteins were extracted from UMSCC-104 and UMSCC-47 cell lines and western blotting used to determine expression of the indicated proteins. C. The protein extracts shown in B were immunoprecipitated with TOPBP1 (top panels) or CIP2A (lower panels) and western blotted for the proteins shown. D. The protein extracts prepared in B were immunoprecipitated with SIRT1 (top panels) or DBC1 (lower panels) and western blotted for the proteins shown. E. The protein extracts prepared in B were immunoprecipitated with an acetyl lysine specific antibody and western blotted for the proteins shown. F. Immunoprecipitation with a TOPBP1 antibody demonstrates a preferential formation of a TOPBP1-ATM complex in UMSCC-104 cells and a failure of TOPBP1 to interact with ATR. This is not observed in UMSCC-47 cells that do not express E2. G. The UMSCC-104 and UMSCC-47 cell lines were treated with the ATM inhibitor AZD0156 (ATMi) for 5 days and then seeded for colony formation. Colonies were fixed and stained with crystal violet after 14-days.

We were unable to detect E2 expression in other HPV16+HNSCC cell lines and this is potentially related to continued growth in culture promoting viral genome integration. To expand the studies of the E2-TOPBP1 complex in cancers, HPV16 positive oropharyngeal cancer (HPV16+OPC) patient derived xenograft (PDX) models were used. The genomic status of these PDXs has been characterized with regard episomal or integrated HPV16 genomes (CD. James, 2024). Figure S6E demonstrates that, when E2 is expressed in cancers with episomal HPV, there is an “ATM up ATR down” phenotype; this western blot was repeated and the results quantitated (Figure S6F). The proteins present in the complex that E2 uses to activate the DDR was investigated in the PDXs. We observed decreased SIRT1 expression and increased cyclin B1 levels in the E2 positive lines (Figure S6G). This was similar to the pre-cancer lines used in Figures 1 and 3 and distinct from the MmuPV1 lesions (Figure 5) and the UMSCC-104 cells (Figure 6B). This western blot was repeated and the protein levels quantitated (Figure S6H). In the E2 positive tumors there was a disruption in the TOPBP1-CIP2A complex, enhanced SIRT1-DBC1 interaction, reduced SIRT1 acetylation and enhanced CHK2 acetylation as well as E2 acetylation (Figure S6I). Tumor OPT61 is an outlier in that there is an “ATM up ATR down” phenotype but there is not an increased CHK2 phosphorylation, even though CHK2 levels are increased, likely via increased acetylation. In this tumor there is also significant residual TOPBP1-CIP2A interaction when compared with the other E2 positive tumors. This demonstrates that in HPV16 positive cancers the presence of E2 by itself is not sufficient to activate ATM signaling in all cases.

## Discussion

HPV are causative agents in around 5% of all cancers worldwide with HPV16 being responsible for the majority of these (zur Hausen, 2009). A seminal paper from Moody and Laimins demonstrated that the DNA damage response is activated during the HPV life cycle, and that ATM inhibition blocks the viral life cycle (Moody and Laimins, 2009). However, the mechanism of how HPV activates the DDR was not fully elucidated. In this study we demonstrate that the papillomavirus transcription/replication/segregation factor E2 is required for DDR activation in differentiating keratinocytes. Our recent work demonstrated that an interaction between HPV16 E2 and the host DDR signaling and repair protein TOPBP1 is required for segregation of viral genomes during the viral life cycle (Prabhakar et al., 2023). During these studies mitotic enriched cells were generated and wild type E2 containing cells had an active DDR, increased protein acetylation and diminished SIRT1 activity which results in enhanced acetylation of viral and cellular proteins via p300 (Prabhakar et al., 2024). HPV infection models have indicated an enriched G2/M environment in differentiating cells (Davy and Doorbar, 2007) which, combined with the requirement of DDR activation during the viral life cycle, prompted us to investigate whether E2 can activate the DDR during differentiation of wild type E2-containing keratinocytes. In non-differentiating cells neither the full genome nor keratinocytes expressing only the viral oncogenes have a significantly different DDR from N/Tert-1+Vec control cells. However, following differentiation induced by calcium addition, or organotypic rafting, E2 activates ATM signaling and suppresses ATR signaling in a TOPBP1 dependent manner. This was reproducible in N/Tert-1 cells expressing wild-type E2, in HFK+HPV16 cells and in W12e cells. We also confirmed, using MmuPV1 FRT lesions, that this “ATM up ATR down” phenotype persists in papillomavirus infected tissues *in vivo*. The preferential activation of ATM over ATR is likely due to the presence of E2 promoting an ATM-TOPBP1 complex that prevents TOPBP1 interaction with ATR. Homologous recombination factors are required for the viral life cycle and ATM activation would promote HR

(Anacker et al., 2014; Chappell et al., 2015; Das et al., 2019; Gillespie et al., 2012; James et al., 2020a; James et al., 2019; James et al., 2024). Activation of ATR, on the other hand, can sense replication stress and promote cell cycle arrest and DNA repair (Kumagai et al., 2006; Nam and Cortez, 2013; Qu et al., 2013; Sowd et al., 2013; Vidal-EycheniÃ© et al., 2013; Xu and Leffak, 2010). Such an arrest would block the viral life cycle. Therefore, the ability of E2 to switch ATM on and prevent ATR activation is an exquisite tuning of the DDR by the virus.

TOPBP1 can segregate fragmented human DNA during mitosis via an interaction with CIP2A, an inhibitor of PP2A that is over expressed in cancer cells (Chen et al., 2023; De Marco Zompit et al., 2022; Lin et al., 2023; Nagelli and Westermarck, 2024; Trivedi et al., 2023). As E2 is required to segregate non-centromeric viral genomes we investigated the ability of E2 to disrupt the TOPBP1-CIP2A interaction during the viral life cycle. CIP2A is a partner protein for the PP2A subunit B56α (Pavic et al., 2023) and there is an enhanced interaction between CIP2A and B56α following CIP2A displacement from TOPBP1. Not only that, CIP2A and B56α are predominantly relocated to the cytoplasm following differentiation of HFK+HPV16 cells, thus disabling the ability of B56α to function in ATM dephosphorylation. Reduced PP2A function enhances ATM phosphorylation, thereby helping activate the DDR in differentiating E2-expressing cells (Goodarzi et al., 2004). It is possible that additional CIP2A partner proteins are contributing to activation of the DDR by E2, but we demonstrate that CIP2A is necessary for DDR activation as CIP2A knockdown abolishes this signaling pathway in E2 cells. ATM promotes the SIRT1-DBC1 interaction (Kim et al., 2008; Zannini et al., 2012; Zhao et al., 2008) and we also observe this in E2 expressing differentiating keratinocytes. This interaction inhibits SIRT1 function, promoting enhanced acetylation of E2 and CHK2 which would, in addition to ATM phosphorylation, boost CHK2 function (Zhang et al., 2020). Knockdown of DBC1 also prevents activation of the DDR during E2 keratinocyte differentiation. Our results reveal that multiple interactions are triggered and required for the sustained E2-induced DDR.

There is a strong correlation between what is observed in mitotic and differentiating E2 cells. There is elevated cyclin B1 (a marker for mitotic cells), increased E2 protein expression, and an active DDR, as well as reduced SIRT1 activity and reduced ability of SIRT1 to interact with substrates including E2 and CHK2. None of these changes are observed in differentiating cells expressing only E6 and E7, demonstrating that these are E2 driven phenotypes. There is no change in the RNA levels of the mentioned proteins during either mitosis or differentiation suggesting E2 has a post-transcriptional mechanism for reprograming mitotic and differentiating cells. The block of SIRT1 activity that occurs in an E2-TOPBP1 interaction dependent manner contributes to the protein changes. Although it is not clear whether cyclin B1 is regulated by acetylation, there are increased cyclin B1 protein levels in HPV cervical lesions (Kostopoulou et al., 2009; Zhao et al., 2006).

While the viral oncogenes can induce DNA damage in infected cells, we show here they are incapable of activating the DDR. An HPV16 genome with stop codons in E6 and E7 increases DNA damage in human keratinocytes when compared with the wild-type genome, and DDR signaling is as active in the wild-type and mutant genome containing cells (James et al., 2020b). This suggests that a potential role for the viral oncogenes during the viral life cycle may be to help the host genome manage the DDR initiated due to E2 expression, to allow host cell replication and therefore proliferation in the presence of the DDR induced by E2. Further work is required to explore this possibility. It is also of interest that, following differentiation, E2-γH2AX PLA identifies 2 large foci in HFK+HPV16 cells even although there are many individual E2 and γH2AX foci. One possible role for the E2-γH2AX proximity could be the recognition of double strand breaks that occur on a specific region of paired chromosomes following differentiation. Such localization of E2 could “kickstart” the DDR signaling pathway that is observed only in E2 positive differentiating cells. Again, further work is required to test this hypothesis.

The similarity between the DDR and its mechanism of activation in E2 positive cells from pre-cancerous *in vitro* HPV16 keratinocyte models and MmuPV1 infected tissue extends to human cancer cells. This suggests that, following the initiation of differentiation, there is an almost immediate activation of the DDR via the E2 mechanism and that, in cancer, the infected cell “escapes” from the differentiation process and continues to proliferate with an active “ATM up ATR down” DDR phenotype. However, if the viral genome integrates with loss of E2 expression, the “ATM up ATR down” phenotype is lost. We show that E2 activation of “ATM up ATR down” creates a vulnerability to ATM inhibitors that require further investigation as potential anti-viral and anti-HPV cancer therapies.

Together, our findings reveal a previously unknown role for the papillomavirus E2 protein in orchestrating a finely tuned manipulation of the host DDR that leverages post-transcriptional control, protein acetylation, and dynamic protein interactions to reshape DDR signaling. This reprogramming is essential for viral infection, persistence and pathogenesis.

## Materials and methods

### Cell Lines and Primary Cultures

Low-passage N/Tert-1 keratinocytes stably expressing HPV16 E2 wild-type (E2-WT), E2-S23A (serine 23 mutated to alanine, abrogating interaction with TopBP1), or empty vector (Vec) were generated as previously described (Prabhakar et al., 2021). N/Tert-1 cells were cultured in keratinocyte serum-free medium (K-SFM; Invitrogen) supplemented with human recombinant epidermal growth factor (EGF; Invitrogen) and bovine pituitary extract (BPE; Invitrogen), 0.3 mM calcium chloride (Sigma-Aldrich), and 150 µg/mL G418 sulfate (Thermo Fisher Scientific). Primary human foreskin keratinocytes (HFKs) were immortalized with either HPV16 wild-type genome (HFK+HPV16) or HPV16 E6/E7 oncogenes (HFK+E6/E7) as previously described (Prabhakar et al., 2021) and cultured in Dermalife-K complete medium (Lifeline Cell Technology) along with mitomycin C-treated 3T3-J2 fibroblast feeders (Kerafast), plated 24 h prior to seeding. W12 episomal and W12 integrated cervical keratinocytes, harboring episomal and integrated HPV16 respectively, were maintained in K-SFM supplemented with EGF, BPE, and calcium chloride and cultured with mitomycin C-inactivated 3T3-J2 fibroblast feeders. UM-SCC-47 cells (male donor, HPV16 integrated) were cultured with 3T3-J2 fibroblast feeders in Dulbecco’s Modified Eagle Medium (DMEM; MilliporeSigma) supplemented with 10% charcoal-stripped fetal bovine serum (FBS; R&D Systems). UM-SCC-104 cells (male donor, HPV16 episomal) were maintained with mitomycin C-inactivated 3T3-J2 fibroblast feeders in Eagle’s Minimum Essential Medium (EMEM; Invitrogen) supplemented with 10% charcoal-stripped FBS and non-essential amino acids (Gibco).

NIH 3T3 mouse embryonic fibroblasts were transfected with pFLAG-CMV constructs encoding FLAG-tagged MmuPV1 E2 wild-type (E2-WT), E2-S23A, or empty FLAG vector (Vec), using calcium phosphate transfection as previously described (Gauson et al., 2014). Stable pools were selected using 750 µg/mL Geneticin® (G418 sulfate; Thermo Fisher Scientific) and maintained in Dulbecco’s Modified Eagle Medium (DMEM; MilliporeSigma) supplemented with 10% FBS.

Mitomycin C-treated 3T3-J2 fibroblast feeders were prepared by incubating 80-90% confluent 3T3-J2 cells (Kerafast) with 4 µg/mL mitomycin C (Sigma-Aldrich) for 4-6 hours at 37°C, followed by PBS washes and trypsinization. Cells were counted, resuspended to 2 × 10⁶ cells/mL in DMEM + 10% FBS or Bambanker freezing media (Bulldog-Bio), and plated 24 hours prior to keratinocyte seeding. Quality control was performed by monitoring treated feeders for proliferation over 5 days to confirm complete inactivation.

All cell lines and primary cultures were maintained at 37°C in a humidified 5% CO₂/95% air atmosphere and were routinely screened for mycoplasma contamination using the LookOut® mycoplasma PCR detection kit (Sigma-Aldrich), following the manufacturer’s protocol. All human cell lines were authenticated via genetic fingerprinting. Mouse 3T3-J2 feeder lines were not independently authenticated but were sourced from vendors with documented quality control procedures. Genomic status of HPV16 in W12 and UM-SCC cell lines was validated using exonuclease V digestion , followed by qPCR to distinguish episomal from integrated viral DNA based on selective degradation of linear genomes and retention of circular episomes (Myers et al., 2020; Witt, 2025).

### Growth of patient derived xenografts

Experiments were performed under Wistar Institute IACUC protocols 201166 and 201178. PDXs were established and passaged in the subcutaneous flank of NSG mice as described (Facompre et al., 2020). The PDXs were harvested in triplicate upon reaching 1 cm^3^. Volumes were calculated as width 2 × length/2. Harvested tumors were flash-frozen in liquid nitrogen and stored at -80°C until processing.

### Generation of MmuPV1 female reproductive tract lesions

Inbred, immunocompetent *FVB/N* mice were purchased from Taconic and housed in the University of Wisconsin School of Medicine and Public Health, Association for Assessment of Laboratory Animal Care-approved, Animal Care Unit. All procedures were carried out in accordance with animal protocols (M005871 Lambert, M006872 Spurgeon) approved by the University of Wisconsin School of Medicine and Public Health Institutional Animal Care and Use Committee.

Female mice were infected with MmuPV1 as previously described (Spurgeon et al., 2019). Briefly, 6-8 week old female *FVB/N* mice were treated 4-7 days prior to infection with medroxyprogesterone acetate (brand name Depo-Provera) to synchronize mice in diestrus. On the day of infection, mice were treated with 4% nonoxynol-9 (VCF; vaginal contraceptive gel) to induce epithelial wounds that potentiate MmuPV1 infection. Approximately four hours after nonoxynol-9 treatment, mice were infected intravaginally with 10^8^ viral genome equivalents (VGE) of MmuPV1 resuspended in 25 µL 4% carboxyl methylcellulose (CMC) (C4888; Sigma). MmuPV1 viral stocks were generated by isolating virions from MmuPV1-induced cutaneous warts arising on immunodeficient *FoxN1^nu/nu^* mice. Tissues were harvested from mice and immediately frozen in liquid nitrogen before storing at -80C.

### Organotypic Raft Culture

N/Tert-1, HFK+HPV16, and HFK+E6/E7 cells were differentiated using organotypic raft culture, as previously described (ref). Briefly, 1 × 10⁶ cells were seeded onto type I rat tail collagen matrices (Advanced BioMatrix) embedded with mitomycin C-treated J2 3T3 fibroblast feeder cells. Once confluent, the cultures were transferred onto wire grids and maintained at the air-liquid interface in standard tissue culture dishes, with growth medium replaced every other day. After 13 days, rafts were harvested and processed for downstream applications. Part of the rafts were fixed in 4% formaldehyde (vol/vol) and embedded in paraffin blocks for histological analysis. Others were flash-frozen and stored at -80°C prior to homogenization and protein extraction.

### Calcium-Induced Differentiation

To induce keratinocyte differentiation, we employed a calcium-switch protocol adapted for monolayer culture (Moody et al., 2007). Mitomycin C-treated 3T3-J2 fibroblast feeders were plated one day prior to seeding keratinocytes, with 1 × 10⁶ cells distributed across six wells and cultured in Dermalife-K (for HFKs) or K-SFM supplemented with growth factors (for N/Tert-1 or W12 cells). On the following day, N/Tert-1, HFK, and W12 cells were seeded at 2.5 × 10⁵ cells per well and maintained in calcium-free growth medium. At 60-70% confluence (Day 0), cells were switched to low-calcium medium (0.03 mM CaCl₂; Sigma-Aldrich), prepared by supplementing M154 CF (Invitrogen) with Human Keratinocyte Growth Supplement (HKGS; Gibco). After 24 hours (Day 1), cultures were transitioned to high-calcium medium (1.5 mM CaCl₂; Sigma-Aldrich), prepared in M154 CF without HKGS. Media were refreshed on alternate days, and cells were maintained in high-calcium conditions for 72 hours. Cells were processed at Day 0 (growing cells/ undifferentiated) and Day 3 (72 hours post high-calcium treatment) for downstream analysis. Differentiation was confirmed by increased expression of involucrin, assessed by quantitative PCR and western blotting.

### Protein isolation

Protein lysates were prepared from monolayer cultures, organotypic raft cultures, patient-derived xenograft (PDX) tissues, and mouse tissues (uninfected or MmuPV1-induced ear warts or FRT lesions). For monolayer cultures, cells were trypsinized using 0.05% Trypsin-EDTA (Life Technologies), washed with 1× PBS, and resuspended in 2× pellet volume of lysis buffer containing 0.5% Nonidet P-40, 50 mM Tris (pH 7.8), and 150 mM NaCl, supplemented with protease inhibitor (Roche Molecular Biochemicals) and phosphatase inhibitor cocktail (Sigma-Aldrich). Lysates were incubated on ice for 20 minutes and centrifuged at 14,000 × g for 15 minutes at 4°C. Supernatants were collected for downstream analysis.

For organotypic raft cultures, mouse tissues, and PDX samples, tissues were flash-frozen in liquid nitrogen and stored at -80°C until processing. Samples were weighed and transferred into pre-chilled microcentrifuge tubes. T-PER Tissue Protein Extraction Reagent (Thermo Fisher Scientific) was added at a ratio of 1 mL per 50 mg of tissue, starting with 500 µL. Tissues were minced using sterile scissors and homogenized using a handheld rotor-stator homogenizer (Omni International), rinsed between samples with distilled water and 70% ethanol. Homogenates were transferred to fresh tubes and rinsed with an additional 500 µL of T-PER to recover residual lysate. Lysates were incubated on ice for 30 minutes and centrifuged at 14,000 × g for 15 minutes at 4°C. The soluble protein fraction was collected and quantified using the Bio-Rad Protein Assay, following the manufacturer’s instructions. Lysates were used immediately or stored at -80°C and used for immunoprecipitation and western blotting.

### Immunoblotting

Protein lysates (100 µg) were mixed with 4× Laemmli sample buffer (Bio-Rad) and denatured at 95°C for 5 minutes. Samples were resolved on Novex 4-12% Tris-glycine gels (Invitrogen) and transferred to nitrocellulose membranes (Bio-Rad) using wet transfer at 30 V overnight at 4°C. Membranes were blocked for 1 hour at room temperature in Odyssey PBS blocking buffer (LI-COR; diluted 1:1 in 1× PBS) and incubated overnight at 4°C with primary antibodies diluted in blocking buffer. The following primary antibodies were used in this study: anti-ATM (D2E2) 1:1,000 (CST), anti-ATR 1:1,000 (CST), anti-ATR (C-1) 1:200 (SCBT), anti-Chk1 (2G1D5) 1:1,000 (CST), anti-Chk2 1:1,000 (CST), anti-CIP2A 1:1,000 (Abcam), anti-Cyclin B1 1:1,000 (CST), anti-DBC1 (3G4) 1:1,000 (CST), anti-DYKDDDDK tag (FLAG®) 1:1,000 (CST), anti-GAPDH (0411) 1:200 (SCBT), anti-HPV16 E2 (B9) 1:500 from 0.5mg/ml concentration stock (Wieland et al., 2020), anti-Involucrin 1:200 (SCBT), anti-MmuPV1 E2 (Clone#14, 1:100, a kind gift from Drs. Elliot Androphy and Neil Christensen), anti-Phospho-(Ser/Thr) ATM/ATR substrate 1:1,000 (CST), anti-Phospho-ATM (Ser1981) 1:500 (Thermo Fisher), anti-Phospho-CHK1 (Ser345) 1:1,000 (CST), anti-Phospho-CHK2 (Thr68) 1:500 (Thermo Fisher), anti-PP2A-B56-α (F-10) 1:200 (SCBT), anti-SIRT1 1:1,000 (Millipore), anti-TOPBP1 1:1,000 (Bethyl), and anti-β-Actin (C4) 1:200 (SCBT). After primary incubation, membranes were washed twice with PBS-Tween (0.1%) and once with 1× PBS, then probed with IRDye-conjugated secondary antibodies (goat anti-mouse IRDye 800CW or goat anti-rabbit IRDye 680CW; LI-COR) diluted 1:10,000 in Odyssey PBS blocking buffer. After final washes, membranes were imaged using the Odyssey CLx Imaging System (LI-COR). Band intensities were quantified using ImageJ (NIH) software, with GAPDH used as an internal loading control. Each panel pair (D–E, F–G, H–I, J-K) in Figure 3 was generated from the same blot, stripped with 0.2M NaOH buffer and reprobed for different targets. E2 and GAPDH controls are reused within each pair.

### Immunoprecipitation

Protein lysates (250 µg) from indicated cells (prepared as described above) were incubated with either the primary antibody of interest or an anti-HA tag antibody (used as a negative control). Antibody was added at a ratio of 1 µg per 100 µg of protein lysate. The lysate-antibody mixture was brought to a final volume of 500 µL using lysis buffer supplemented with protease inhibitors (Roche) and phosphatase inhibitor cocktail (Sigma-Aldrich), and rotated end-over-end overnight at 4°C. The following day, 40 µL of prewashed protein A agarose beads (Sigma-Aldrich) were added to each sample. Beads were prewashed in lysis buffer according to the manufacturer’s protocol. Samples were rotated for an additional 4 hours at 4 °C. Immunocomplexes were pelleted by centrifugation at 1,000 × g for 3 minutes and washed three times with 500 µL of lysis buffer. After the final wash, bead pellets were resuspended in 4× Laemmli sample buffer (Bio-Rad), heat-denatured at 95 °C for 5 minutes, and centrifuged at 1,000 × g for 3 minutes. Eluted proteins were separated by SDS-PAGE and transferred to nitrocellulose membranes using the wet transfer method. Membranes were probed for target proteins using the western blotting protocol described above.

### Small interfering RNA treatment

HFK cells or N/Tert-1 cells were plated on Mitomycin C-treated 3T3-J2 fibroblast feeders in 10 cm plates using their growth respective media. The following day, cells were transfected with 10 µM of the small interfering RNA (siRNA) to deplete endogenous CIP2A or DBC1. A 10 µM non-targeting control siRNA (MISSION® siRNA Universal Negative Control #1; Sigma-Aldrich) was included in all experiments. Transfections were performed using Lipofectamine™ 2000 transfection reagent (Invitrogen) according to the manufacturer’s protocol. Cells were harvested 48 hours post-transfection, and immunoblotting was performed as described above to confirm knockdown of the target protein. All siRNAs were custom-synthesized and purchased from Sigma-Aldrich. Sequences and target annotations are provided in the Key Resources Table.

### RNA Isolation and Quantitative Reverse Transcription PCR

Total RNA was extracted using the SV Total RNA Isolation System (Promega) following the manufacturer’s protocol. 2 µg of purified RNA was reverse-transcribed into complementary DNA (cDNA) using the High-Capacity cDNA Reverse Transcription Kit (Applied Biosystems). The resulting cDNA was used as input for quantitative reverse transcription PCR (qRT-PCR).

qRT-PCR was performed using Perfecta SYBR FastMix (VWR) and gene-specific primers on a 7500 Fast Real-Time PCR System (Applied Biosystems). Relative transcript levels were calculated using the 2^-ΔΔCt method, normalized to GAPDH expression. Primer sequences are listed in the Key Resources Table.

### Immunofluorescence

N/Tert-1 vector control, HFK HPV16, and HFK E6/E7 cells with mitomycin C–treated 3T3-J2 fibroblast feeders were seeded onto acid-washed, poly-L-lysine–coated coverslips placed in six-well plates at a density of 2 × 10⁵ cells per well per well in their respective growth media. Cells were either calcium-differentiated or maintained in undifferentiated conditions as described above. At the specified time point, all samples were rinsed twice with PBS and fixed in ice-cold methanol for 10 minutes. Fixed cells were permeabilized with 0.2% Triton X-100 in PBS for 15 minutes at room temperature. To reduce background staining, coverslips were blocked in 10% normal goat serum (Life Technologies). Primary antibodies used for staining in this study were as follows: anti-HPV16 E2 (B9) 1:200 from 0.5mg/ml concentration stock (Wieland et al., 2020), anti-CIP2A 1:500 (Abcam), anti-PP2A 1:50 (SCBT), anti-phospho-histone H2A.X (γH2AX) 1:200 (CST). Following primary incubation, cells were washed and incubated with secondary antibodies diluted 1:1,000: Alexa Fluor 488 goat anti-mouse (Thermo Fisher, catalog no. A-11001) and Alexa Fluor 594 goat anti-rabbit (Thermo Fisher, catalog no. A-11037). Wash steps were repeated after secondary incubation. Nuclei were counterstained with DAPI (Santa Cruz Biotechnology, catalog no. sc-3598), and coverslips were mounted using Vectashield mounting medium (Thermo Fisher). Fluorescence imaging was performed using a Zeiss LSM700 laser scanning confocal microscope. Image acquisition and quantification were conducted using Zen LE software (Zeiss) and the Keyence BZ-X810 analysis system.

### Proximity Ligation Assay (PLA)

Protein–protein interactions were visualized using the Duolink® In Situ Red Kit (MilliporeSigma) according to the manufacturer’s protocol. Cells were incubated with primary antibodies from different species (yH2AX mouse Cell Signaling Technology 80312; yH2AX rabbit, Cell Signaling Technology 9718; HPV16 E2 TVG261), followed by PLUS and MINUS probes, ligation, and rolling circle amplification. Nuclei were counterstained with DAPI. Negative controls omitting one or both primary antibodies were included. Images were acquired on a Keyence fluorescence microscope and quantified in ImageJ.

### ATM and ATR inhibition

To inhibit ATM or ATR kinase activity, cells were treated with the ATM inhibitor, AZD0156 (MedChemExpress) or the ATR inhibitor, AZD6738 (MedChemExpress), dissolved in DMSO. N/Tert-1 Vec, E2-WT, and S23A mutant cells; HFK+HPV16 and HFK+E6/E7; and UM-SCC-104 and UM-SCC-47 cells were seeded in in 6-well plates and allowed to adhere overnight, reaching ∼60-80% confluency at the time of treatment. The following day, cells were treated with AZD0156 or AZD6738 at final concentrations of 0 µM, 100 nM, or 1 µM. For each concentration, the corresponding volume of DMSO used to deliver the drug was matched in vehicle control wells to ensure consistency across conditions. Keratinocytes were treated for 48 hours, while UM-SCC-104 and UM-SCC-47 cells were treated continuously for 5 days. Following treatment, cells were trypsinized, counted, and replated at low density (1000-5,000 cells per well depending on cell line) in fresh drug-free medium for clonogenic survival assessment.

### Clonogenic Survival Assay

Following ATM treatment as described above, cells were harvested, counted, and replated at low density (1000-5,000 cells per well depending on cell line) in fresh drug-free medium. Cultures were maintained for 10–14 days to allow colony formation. Colonies were fixed in ice-cold methanol (−20 °C) for 10 minutes and stained with 0.5% crystal violet and imaged by photographing the plates using a standard digital camera under consistent lighting conditions. For quantification, crystal violet was eluted using 10% acetic acid, and absorbance was measured at 590 nm using a spectrophotometer. Relative survival was calculated by normalizing absorbance of treated cells to DMSO controls.

### Exonuclease V assay

PCR-based analysis of HPV16 genome status was performed as previously described by Myers et al. (Myers et al., 2020). Genomic DNA (20 ng) was incubated with Exonuclease V (RecBCD; New England Biolabs) or left untreated as a no-enzyme control. Digestion reactions were performed at 37 °C for 1 hour, followed by heat inactivation at 95 °C for 10 minutes. To validate enzyme activity, reactions containing linearized HPV16 DNA and circular plasmid DNA were processed in parallel with genomic DNA. Following digestion, 2 ng of each sample was subjected to quantitative PCR using SYBR Green PCR Master Mix (Applied Biosystems) and 100 nM of primer in a 20 μl reaction on a 7500 FAST Real-Time PCR System (Applied Biosystems). Details about the primer sets included are mentioned in the Key Resource Table. Thermal cycling conditions were as follows: 50 °C for 2 minutes, 95 °C for 10 minutes, followed by 40 cycles of 95 °C for 15 seconds, and a dissociation stage consisting of 95 °C for 15 seconds, 60 °C for 1 minute, 95 °C for 15 seconds, and 60 °C for 15 seconds. Ct values were analyzed in Microsoft Excel, and the difference between ExoV-treated and untreated samples (ΔCt = Ct_ExoV − Ct_control) was used to infer DNA susceptibility to digestion, indicating the relative abundance of linear versus circular DNA.

### Quantitation and statistical analysis

All data are presented as mean ± standard error of the mean (SEM), calculated from independent biological replicates. Statistical significance was assessed using a two-tailed Student’s *t* test, unless otherwise specified. A *p*-value < 0.05 was considered statistically significant. Statistical analyses and graphs were generated with Microsoft Excel software. Western blot band intensities were quantified using ImageJ, normalized to loading controls (e.g., GAPDH, β-actin), and expressed relative to control conditions. Immunofluorescence signal intensities were measured using Zen LE (Zeiss) or Keyence BZ-X Analyzer, with consistent exposure settings across replicates. For nuclear versus cytoplasmic signal comparisons, regions of interest (ROIs) were defined using Keyence BZ-X800 analysis software, which automatically segmented nuclei based on DAPI fluorescence and quantified signal intensity across channels. Measurements were performed across ≥5 fields per condition, with consistent exposure settings and threshold parameters applied to all replicates.

## Acknowledgements

We thank Zhi-Ming (Thomas) Chen, National Cancer Institute, for the gift of the Flag-tagged mouse E2 expression plasmid. We also thank Neil Christensen, Penn State, for the gift of the mouse E2 antibody. We acknowledge the NIH that supported this work: R01DE029471 and R21 AI178143 (IMM); P01CA022443 (PFL); R01DE034056 (DB); R00DE030194 (AD);R21AI174247 (EJA). MES is supported by startup funds provided by the John W. and Jeanne M. Rowe Center for Research in Virology and the Morgridge Institute for Research

