## Supplemental Figures for "E2 displacement of CIP2A from TOPBP1 activates the DNA damage response during papillomavirus life cycles"

Figure S1.

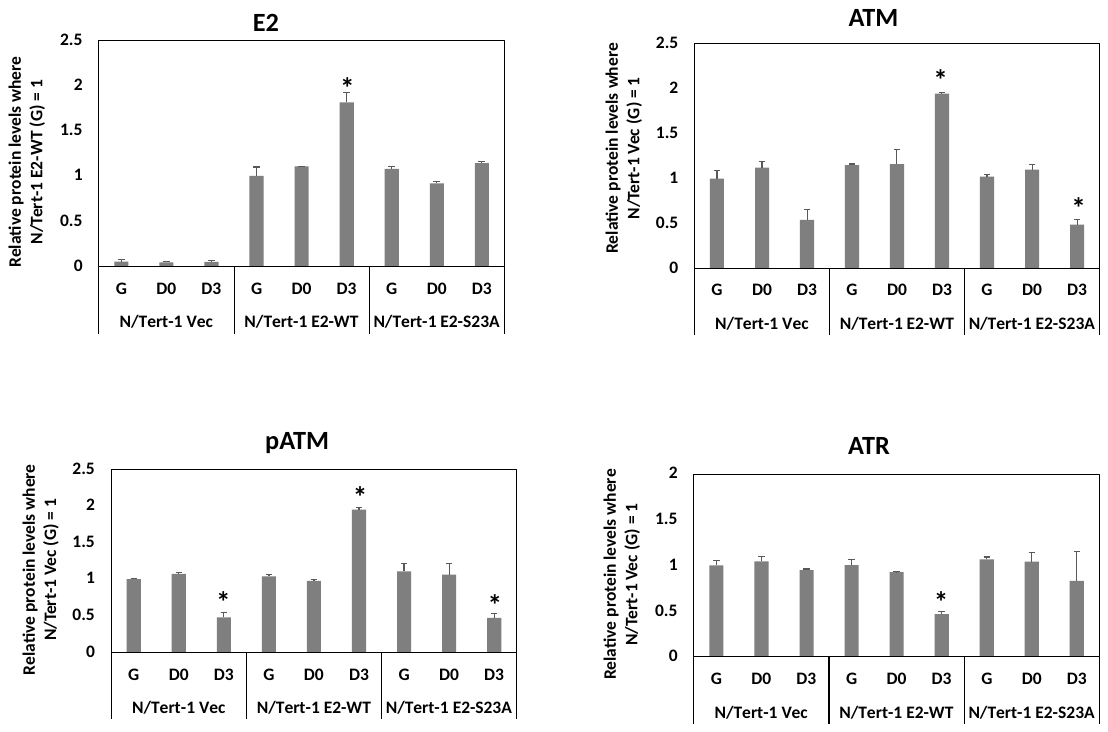
A

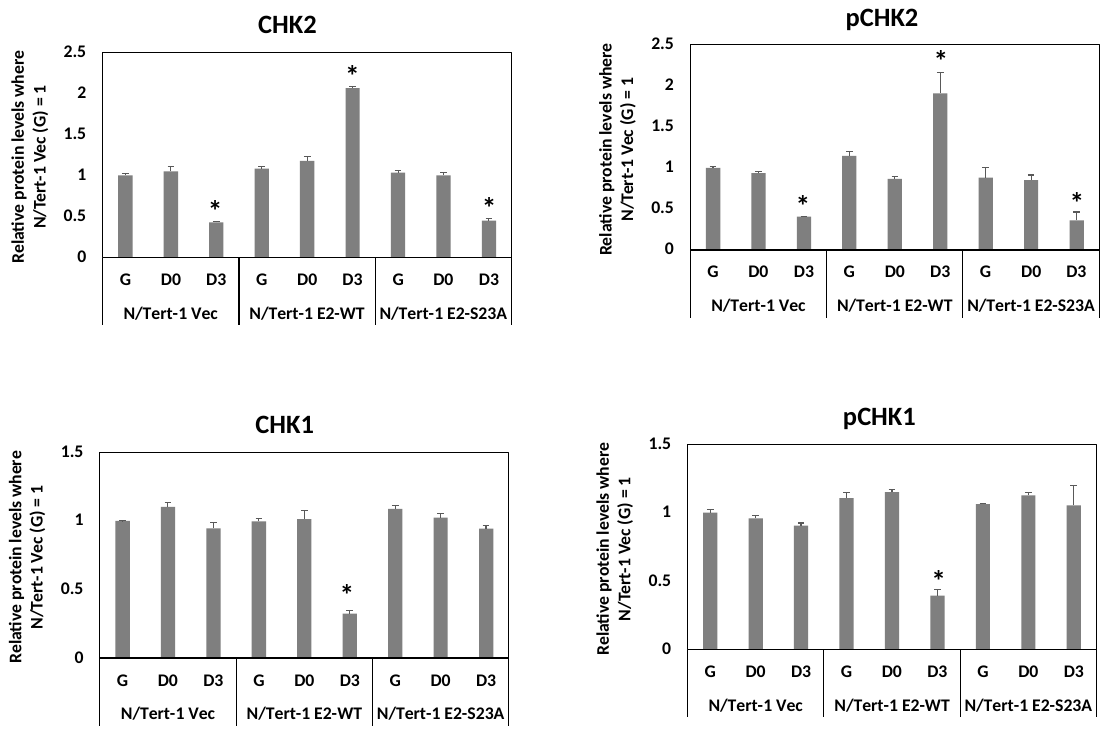

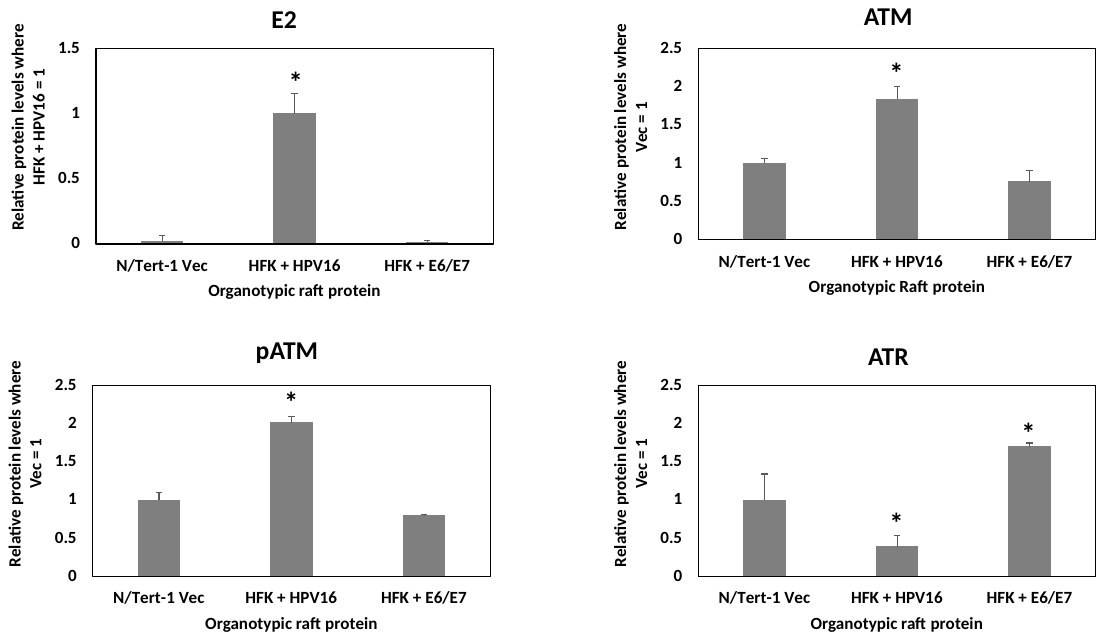
B

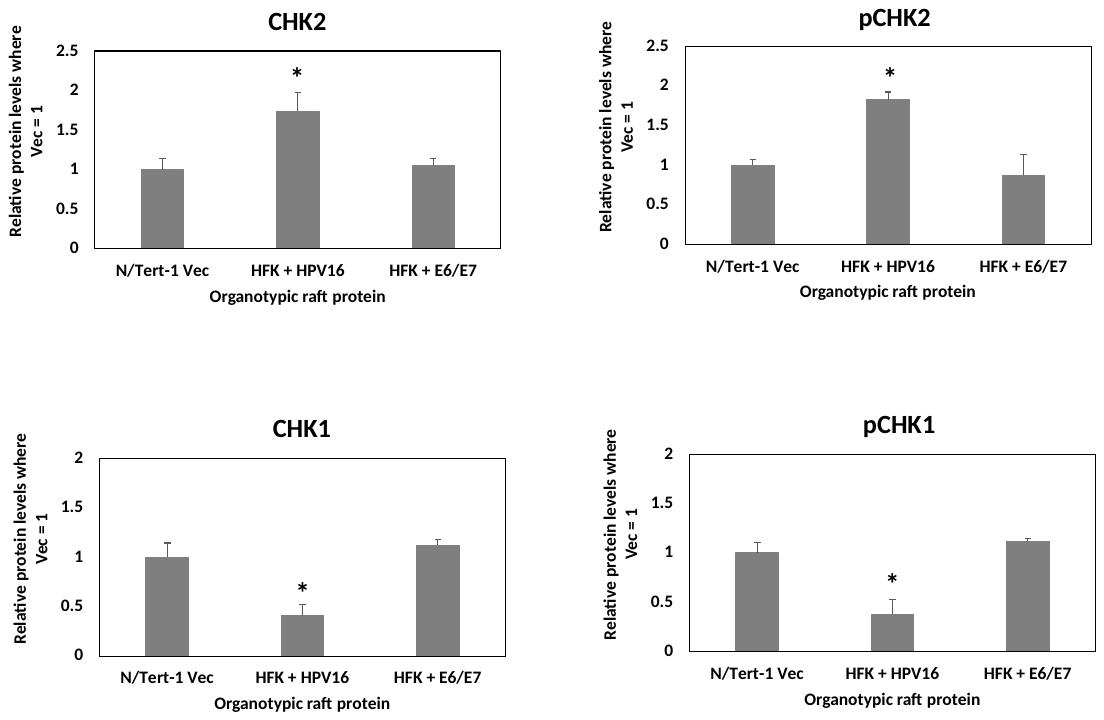

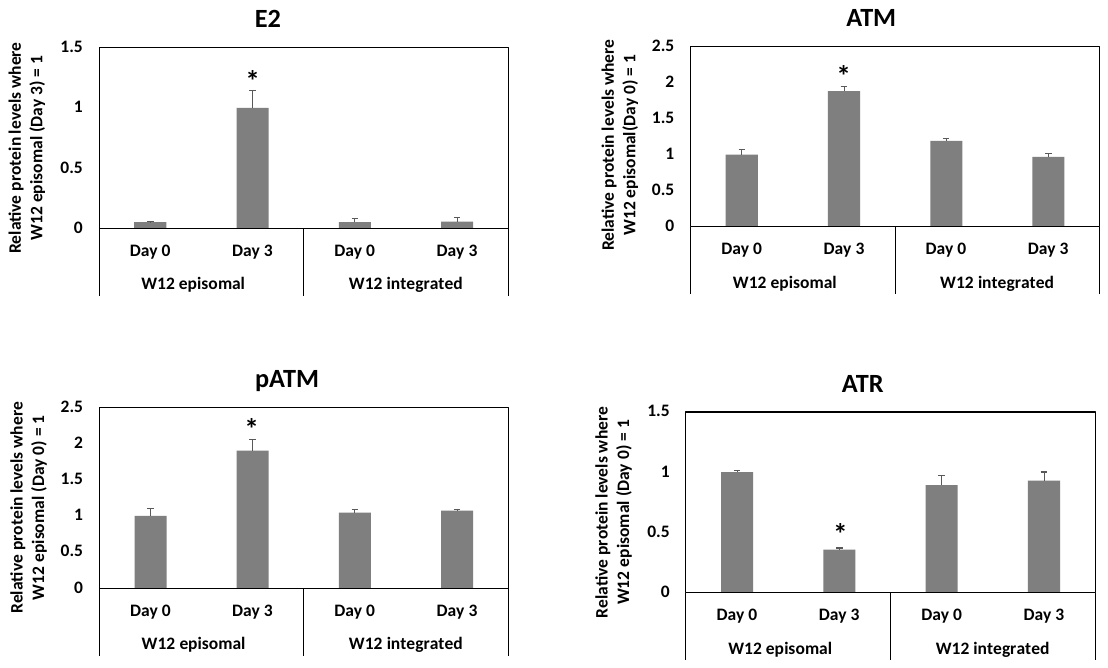
C

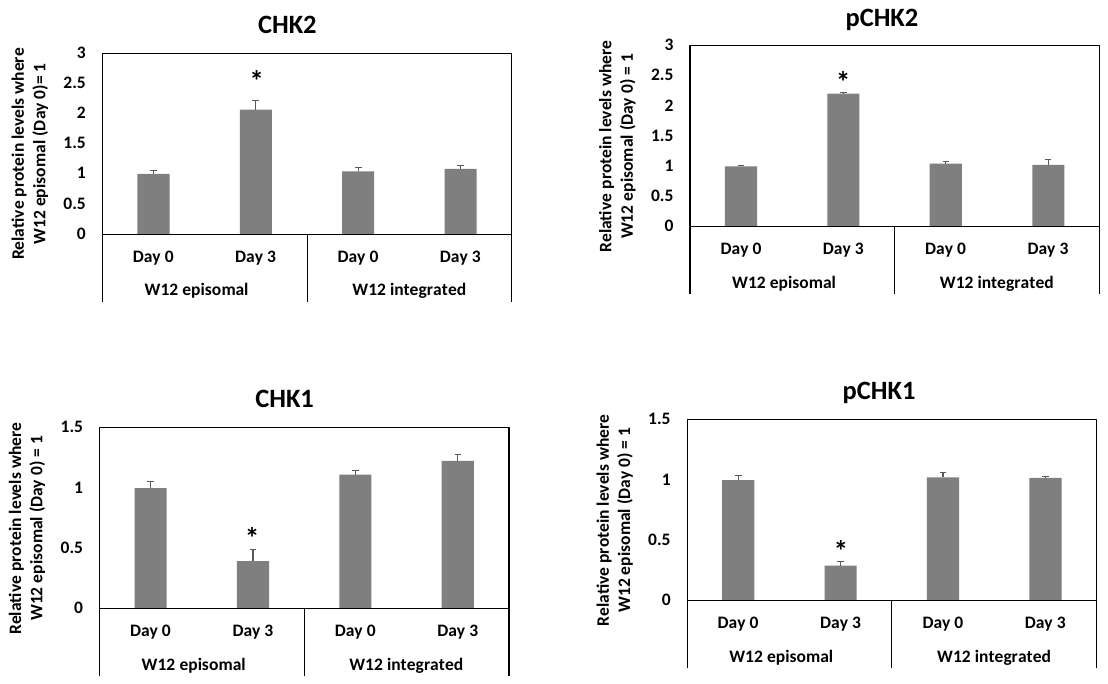

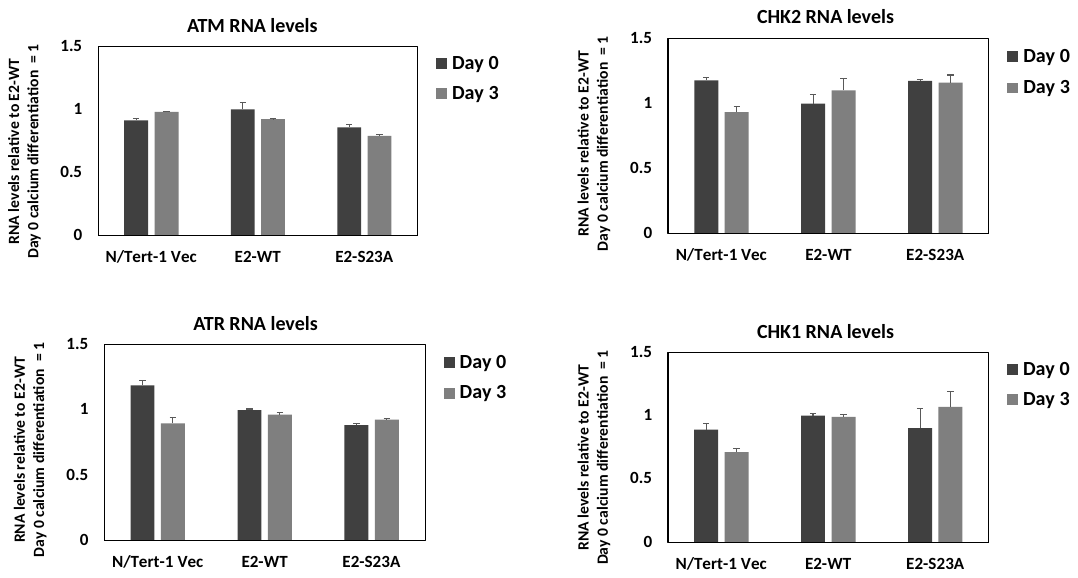
D

E

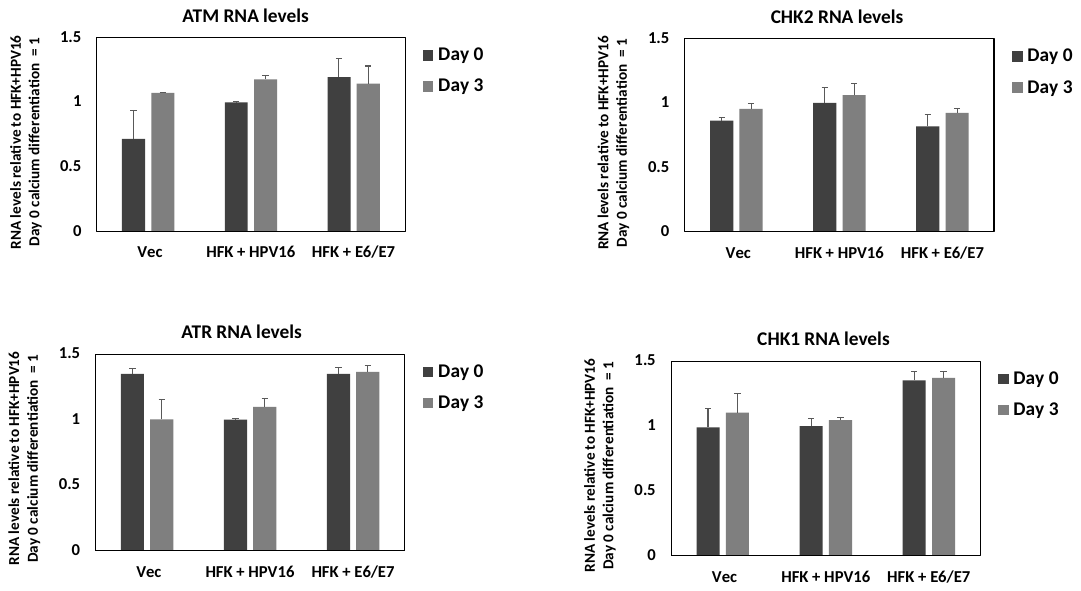

F

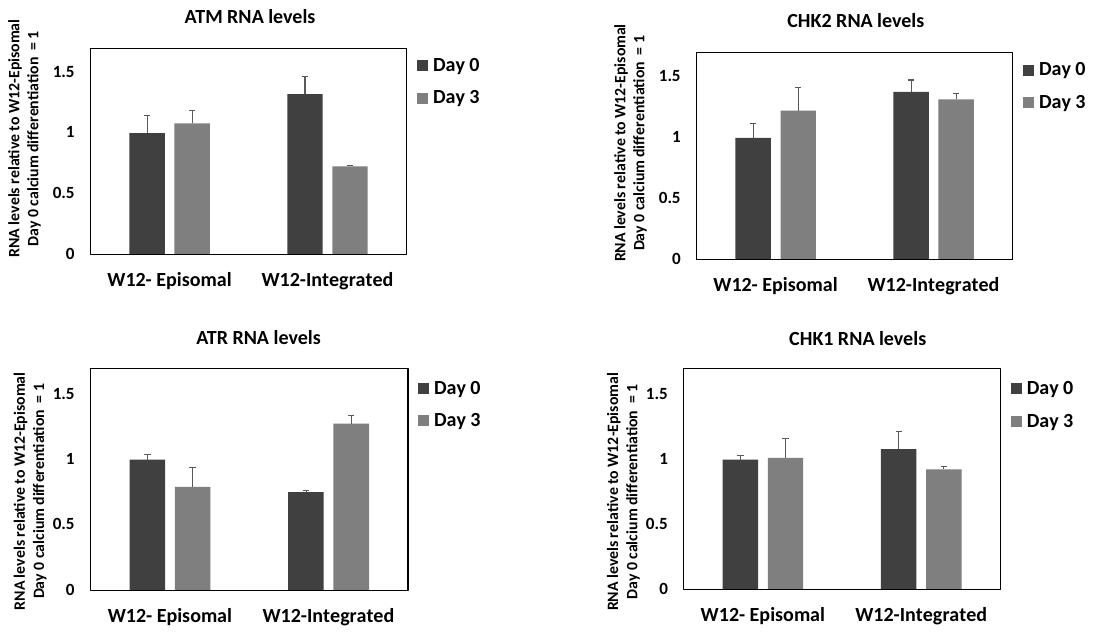

Figure S1. A. The experiment in Figure 1A was repeated and densitometry used to determine the levels of proteins relative to N/Tert-1+Vec levels equaling 1, apart from E2 where the levels in N/Tert-1+E2-WT in growing cells was set as 1. B. The western blots from the organotypic rafts in Figure 1B were repeated and densitometry used to determine the levels of proteins relative to N/Tert-1+Vec levels equaling 1, apart from E2 where the levels in HFK+HPV16 was set as 1. C. The experiment in Figure 1C was repeated and densitometry used to determine the levels of proteins relative to W12e levels equaling 1, apart from E2 where the levels in W12e at Day 3 was set as 1. D. RNA was prepared from the indicated N/Tert-1 cells from two individual experiments and RNA levels for the indicated genes are shown relative to the levels in N/Tert-1+E2 Day 0 equaling 1. E. RNA was prepared from the indicated cell lines from two individual experiments and RNA levels for the indicated genes are shown relative to HFK+HPV16 Day 0 cells equaling 1. E. RNA was prepared from the indicated W12 cells from two individual experiments and RNA levels for the indicated genes shown relative to W12e Day 0 equaling 1. * indicates a significant difference from the growing samples (S1A) or the Day 0 samples (S1B and S1C), p-value < 0.05.

Figure S2.

A

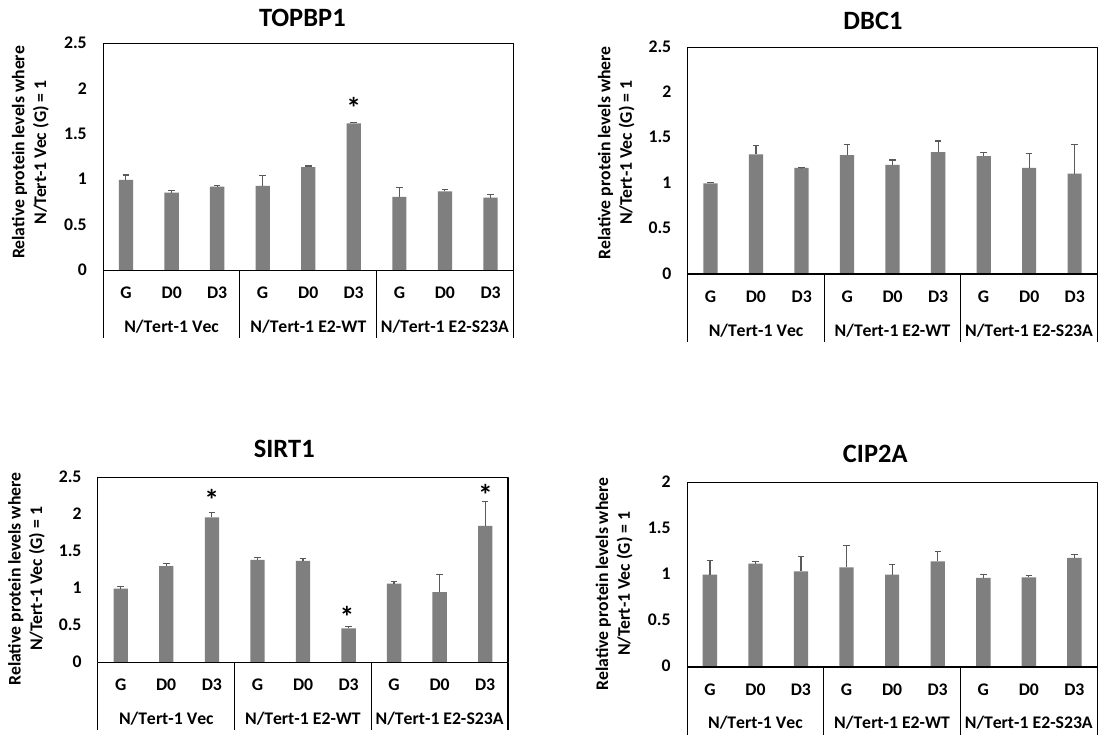

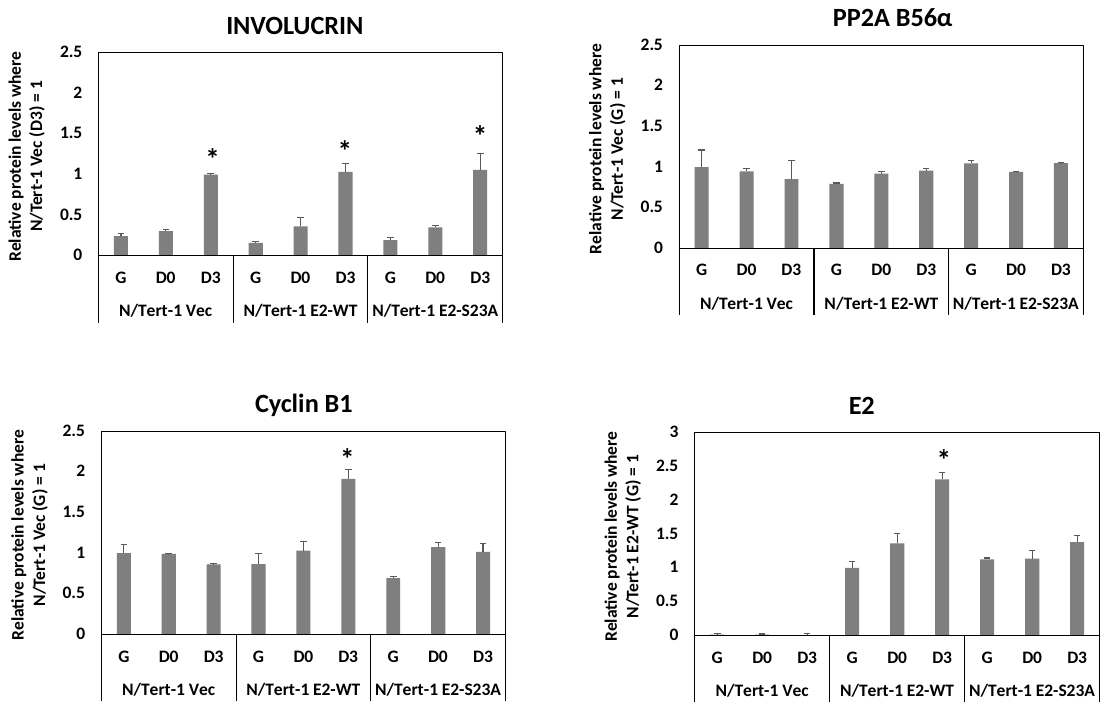

B

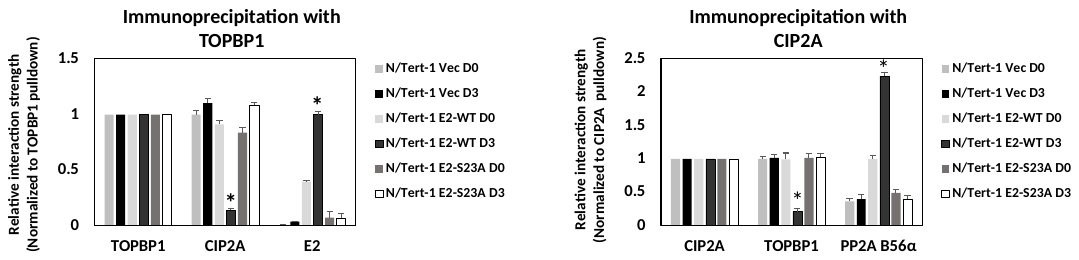

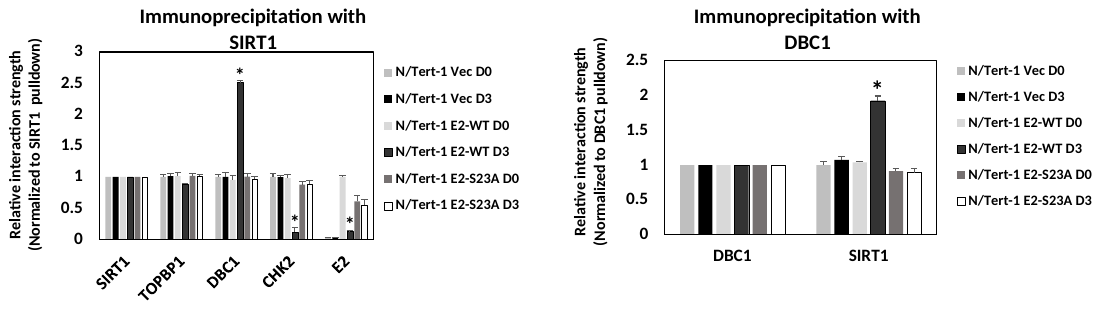

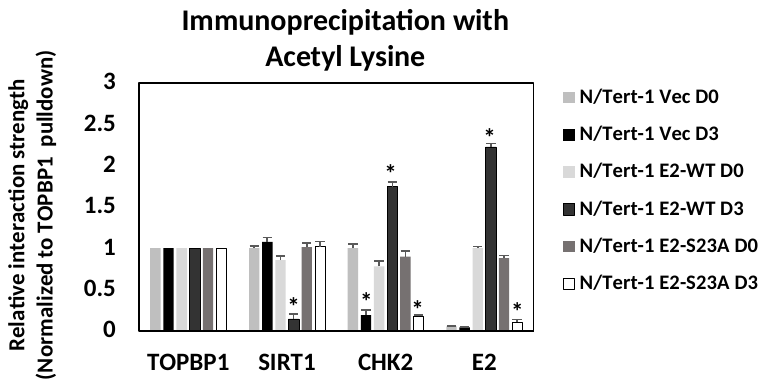

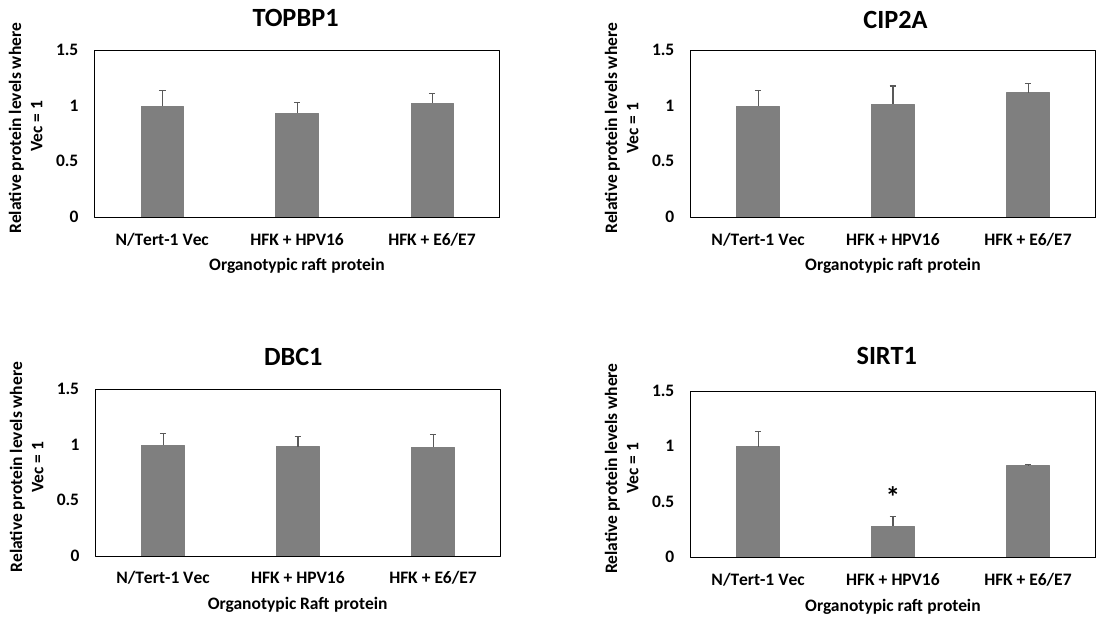
C

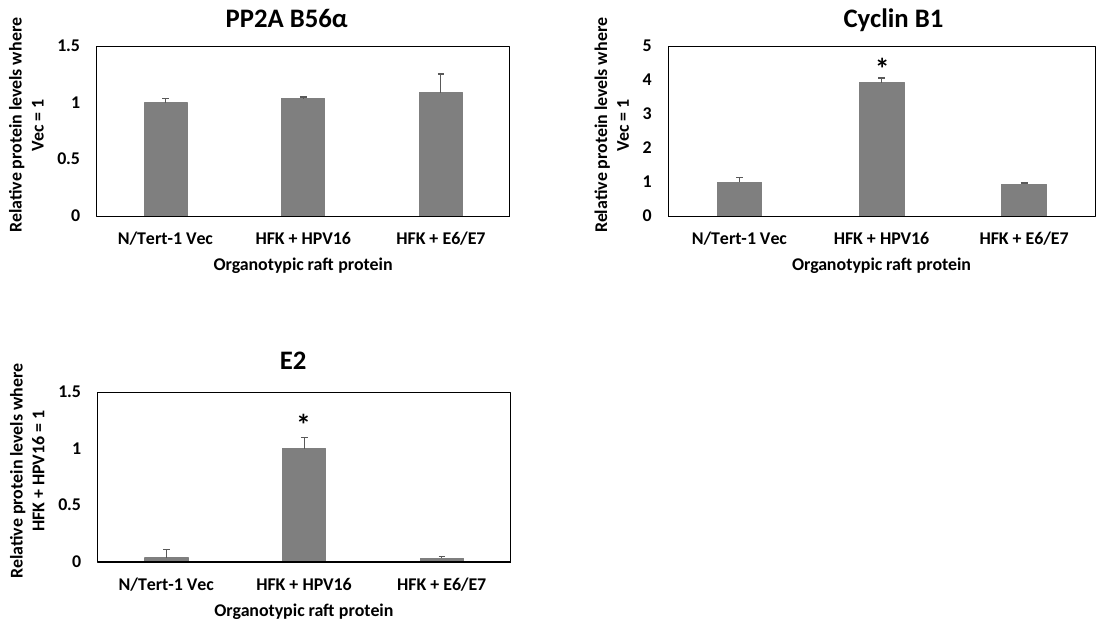

D

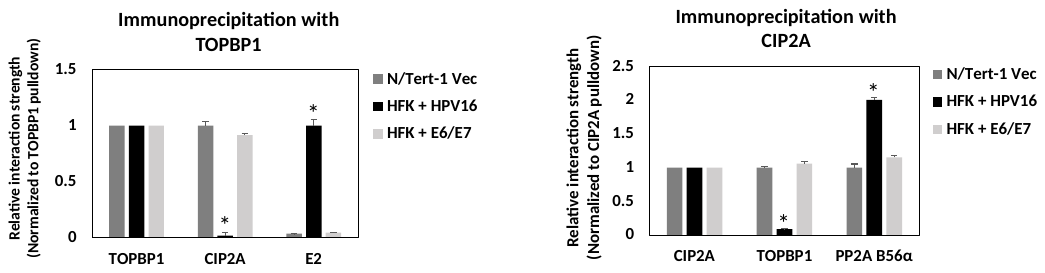

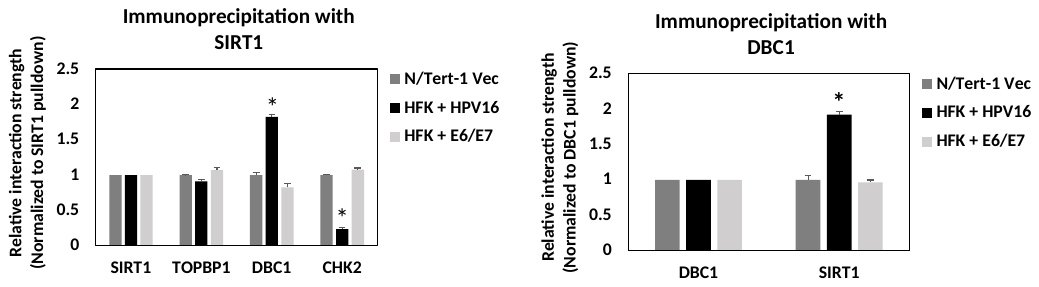

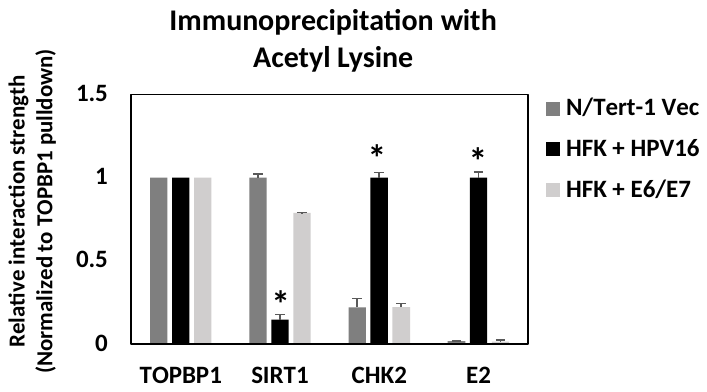

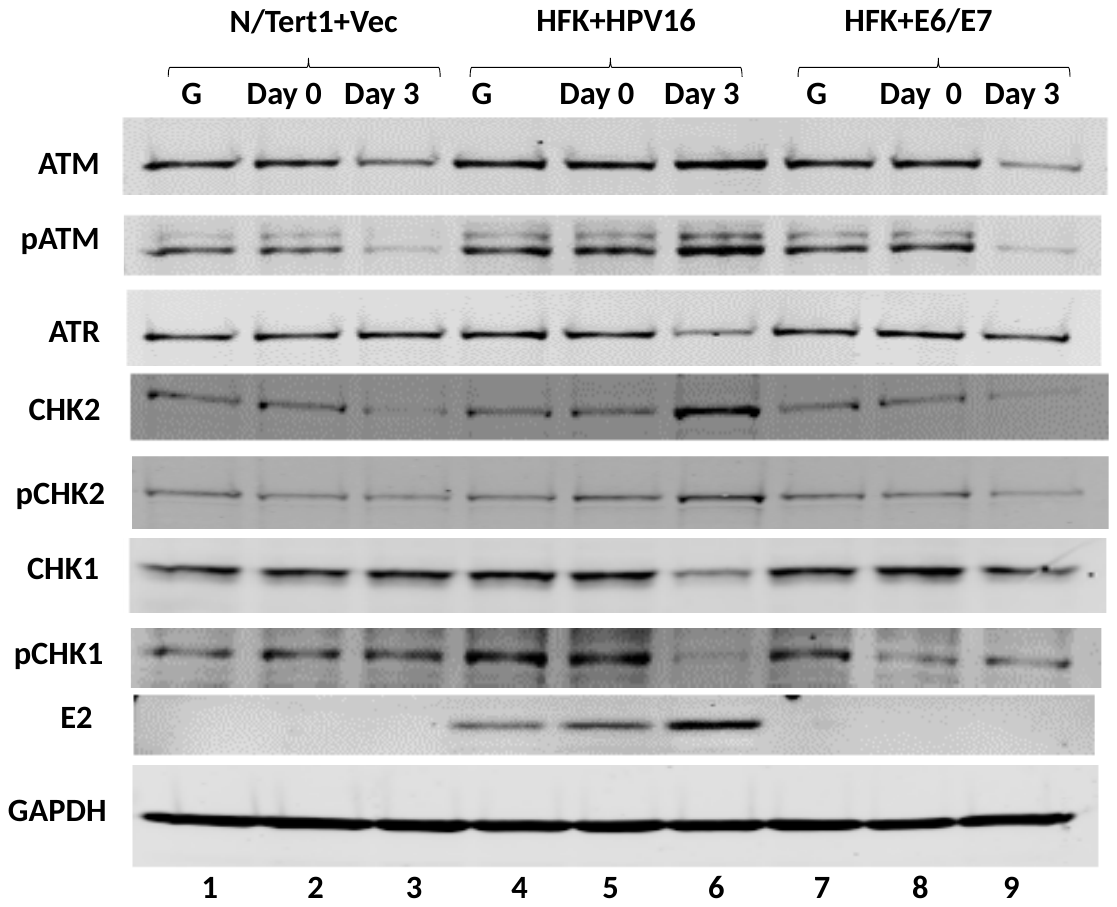
E

F

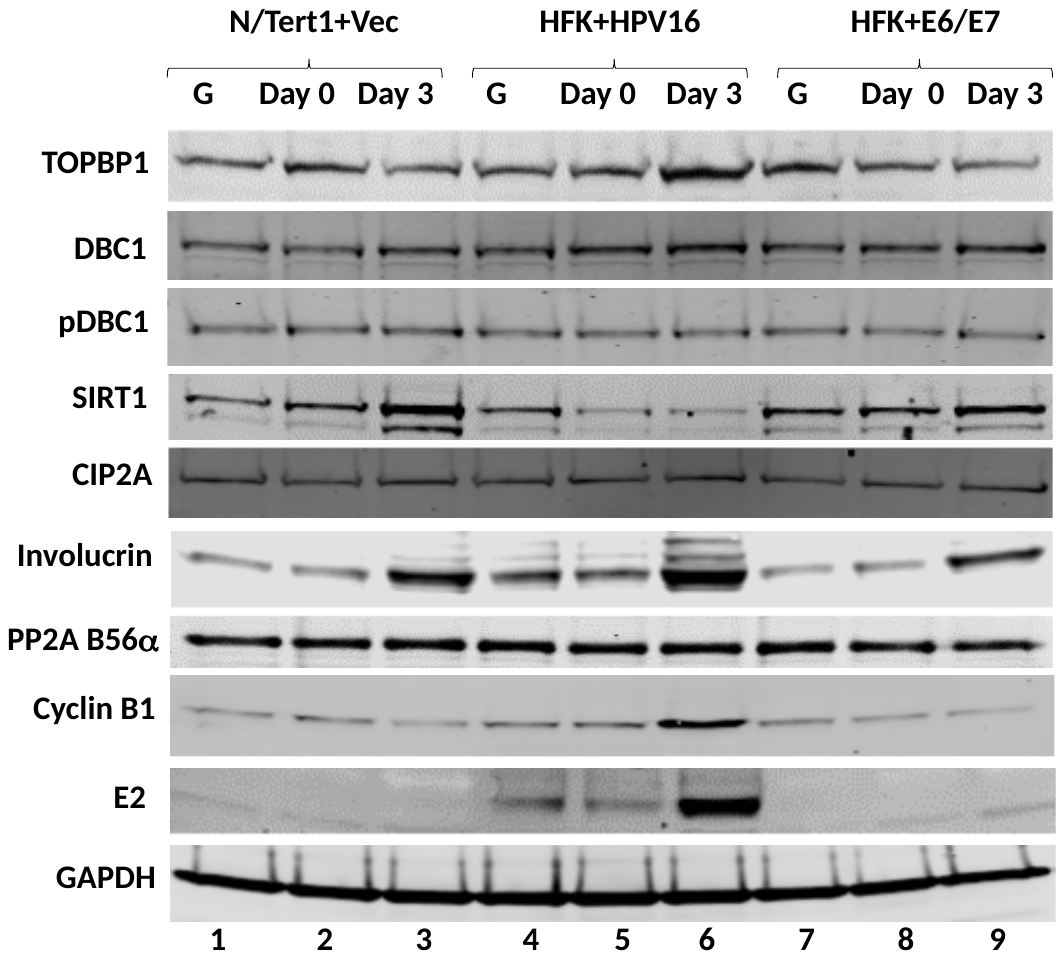

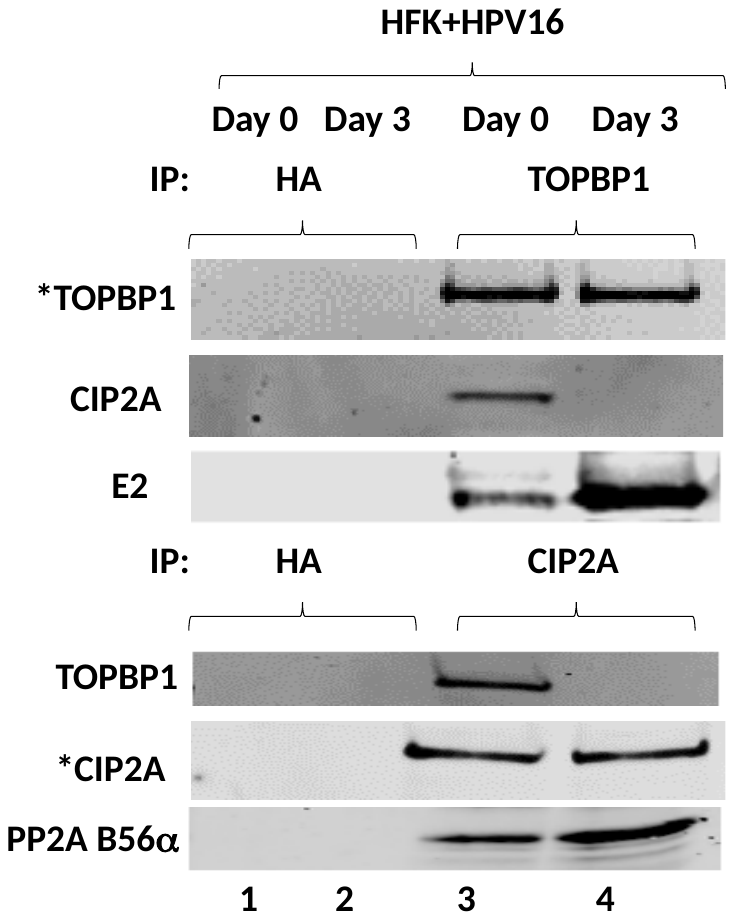
G	H	I

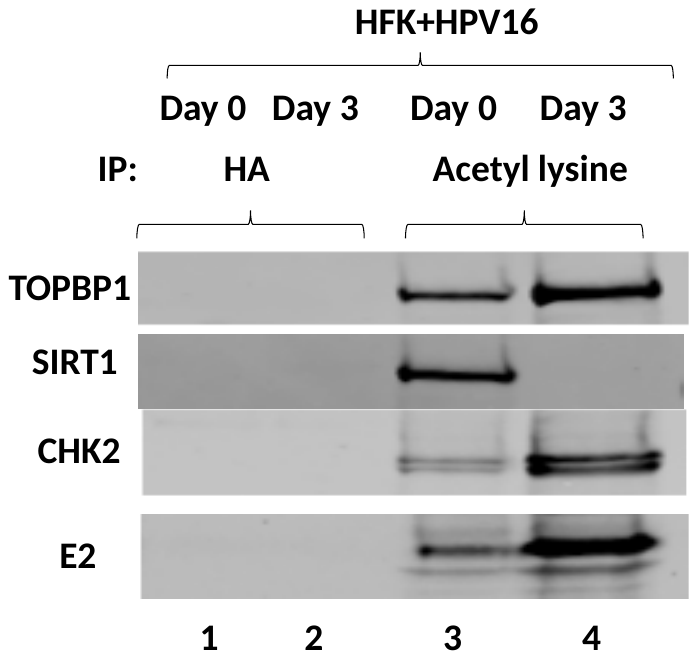

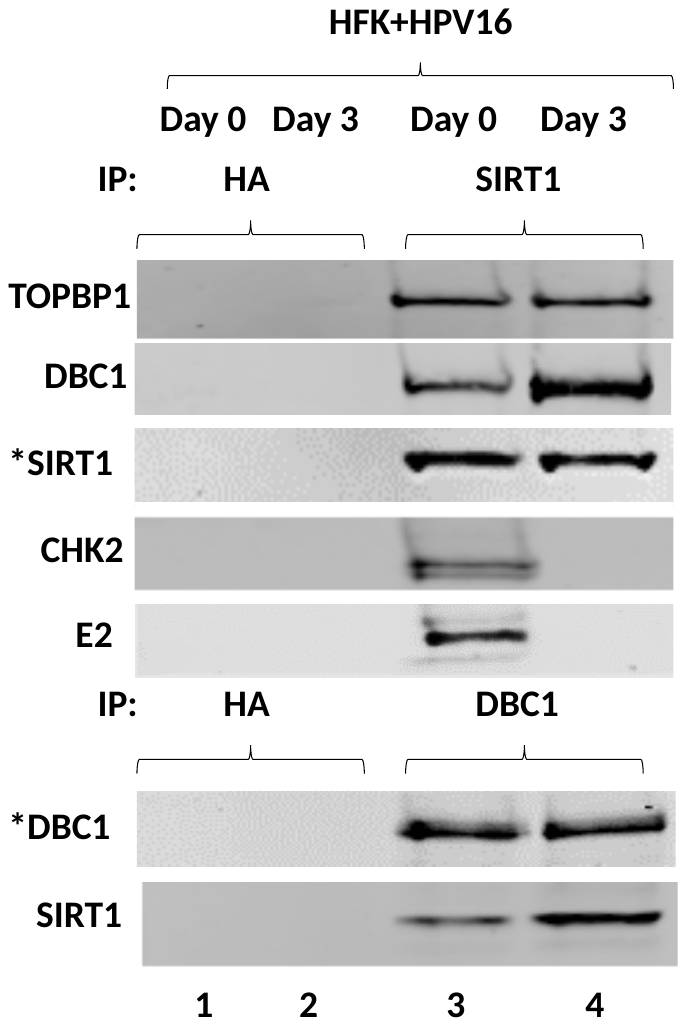

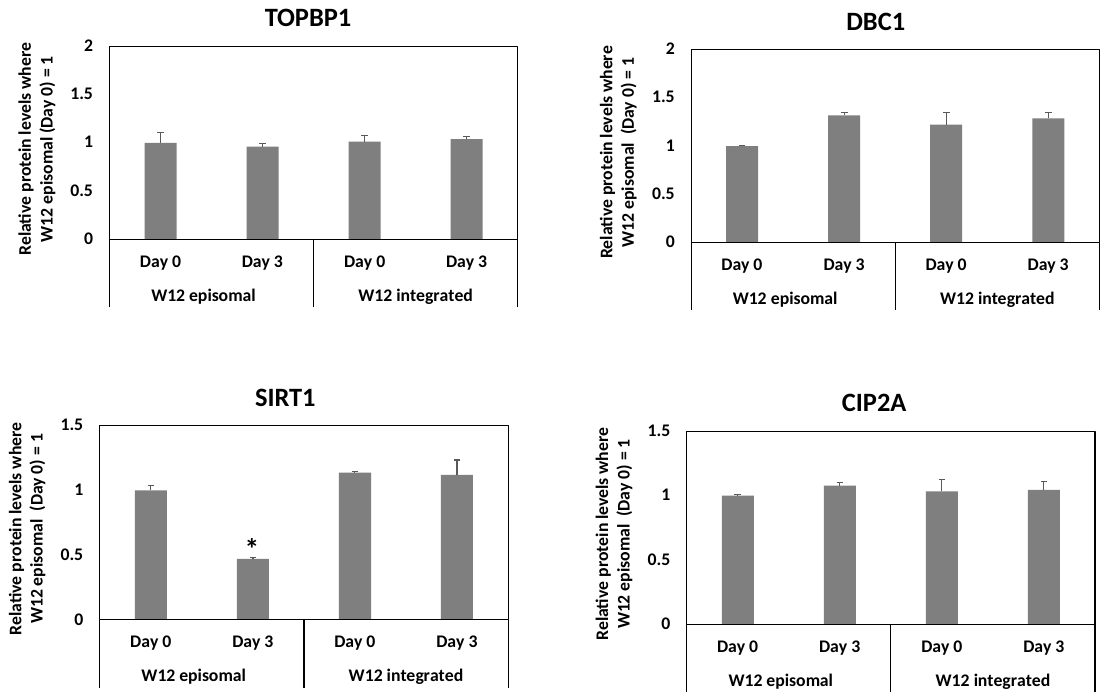
J

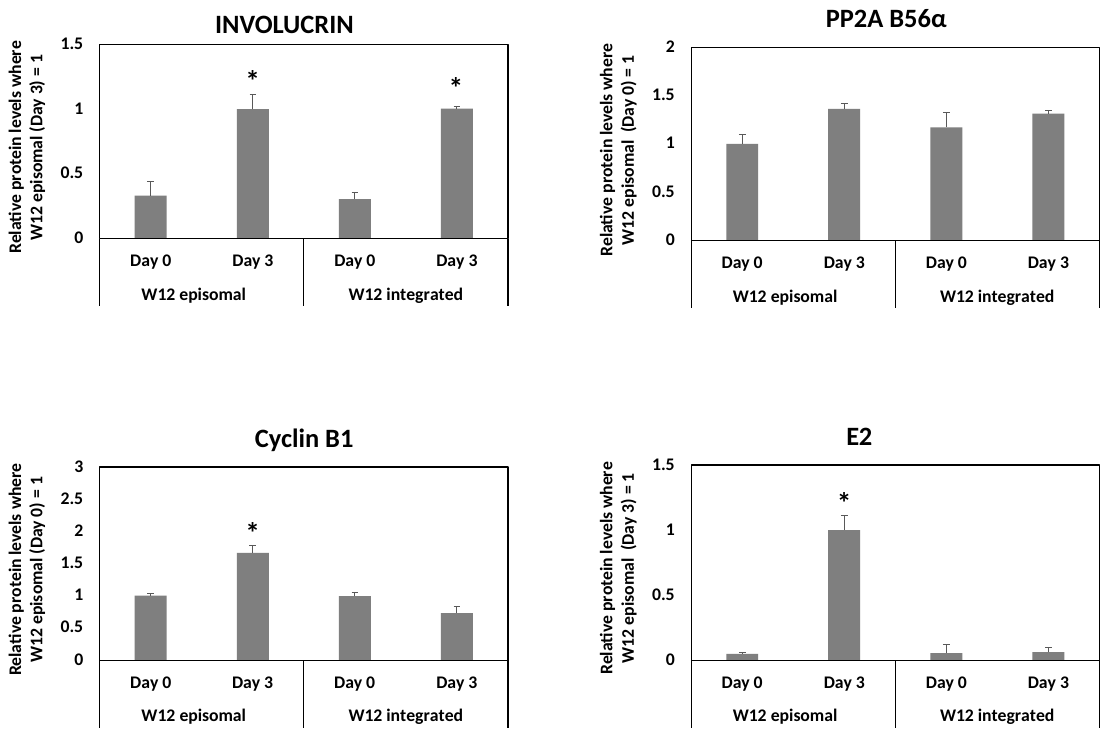

K

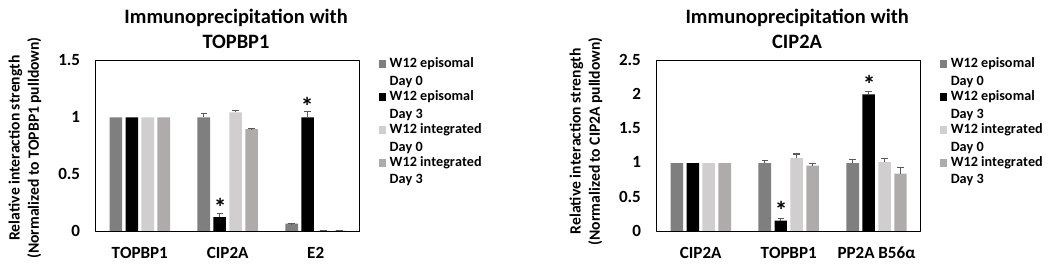

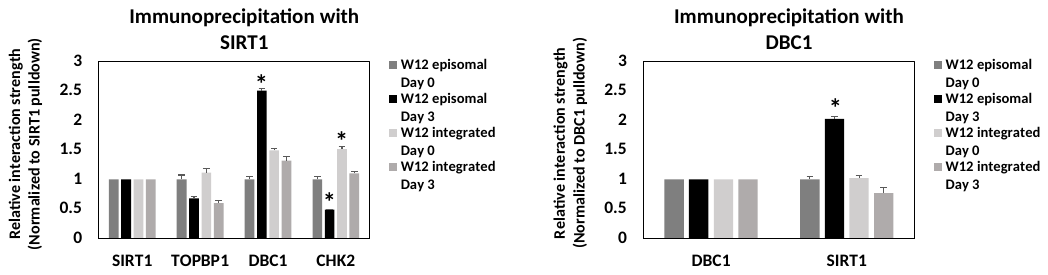

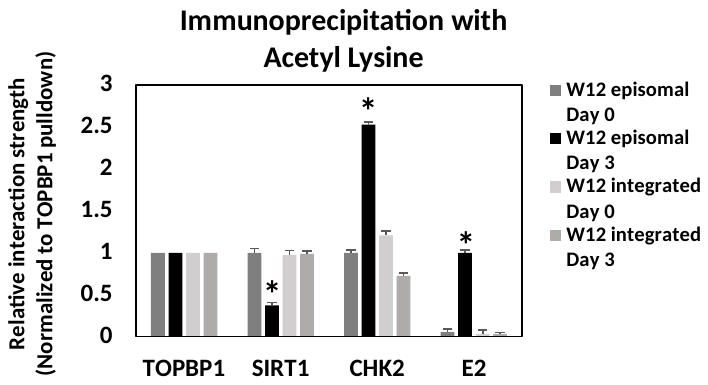

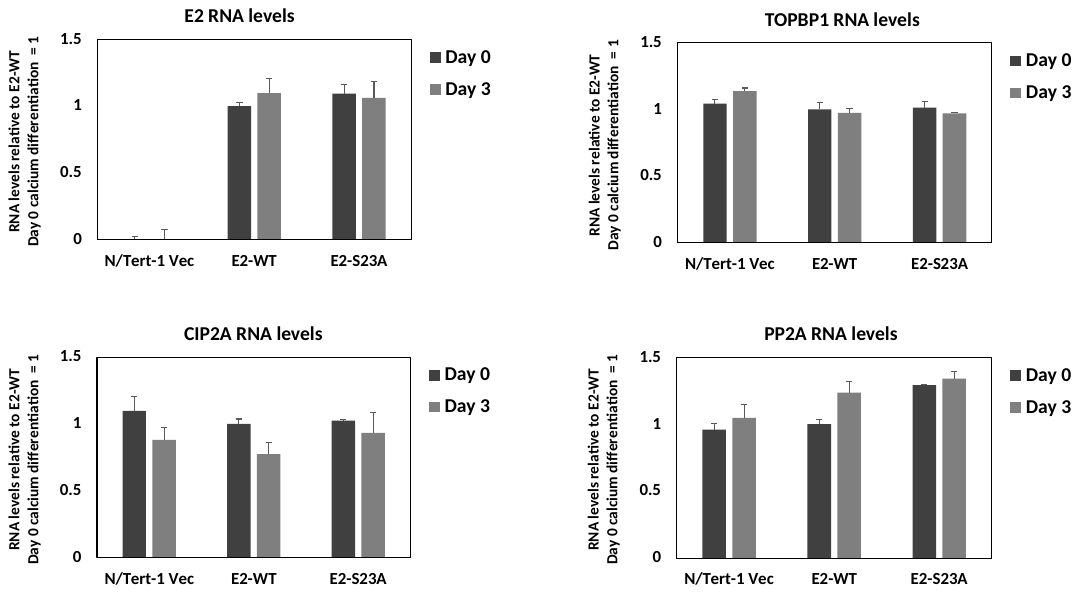
L

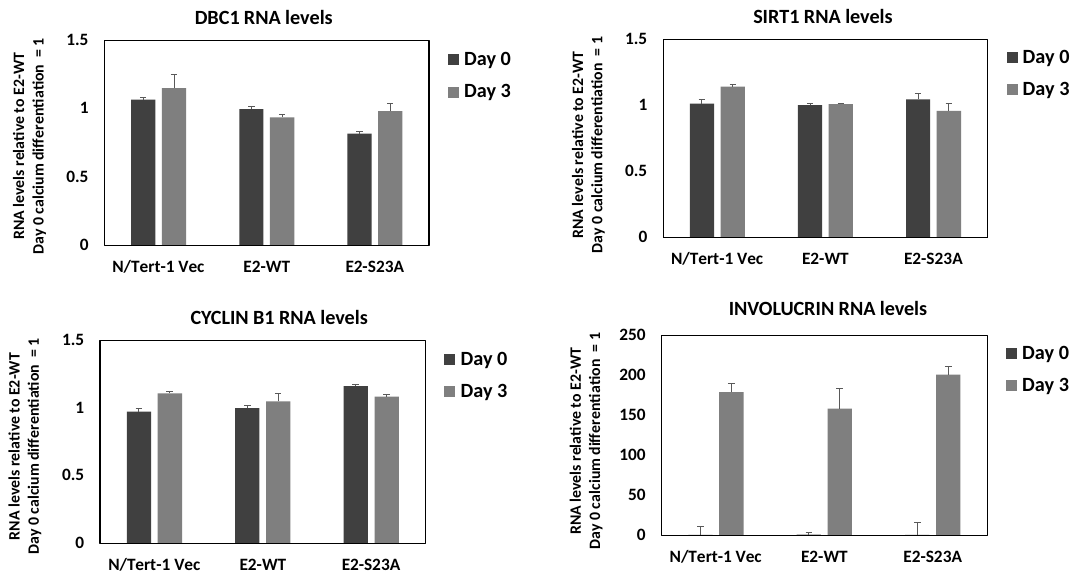

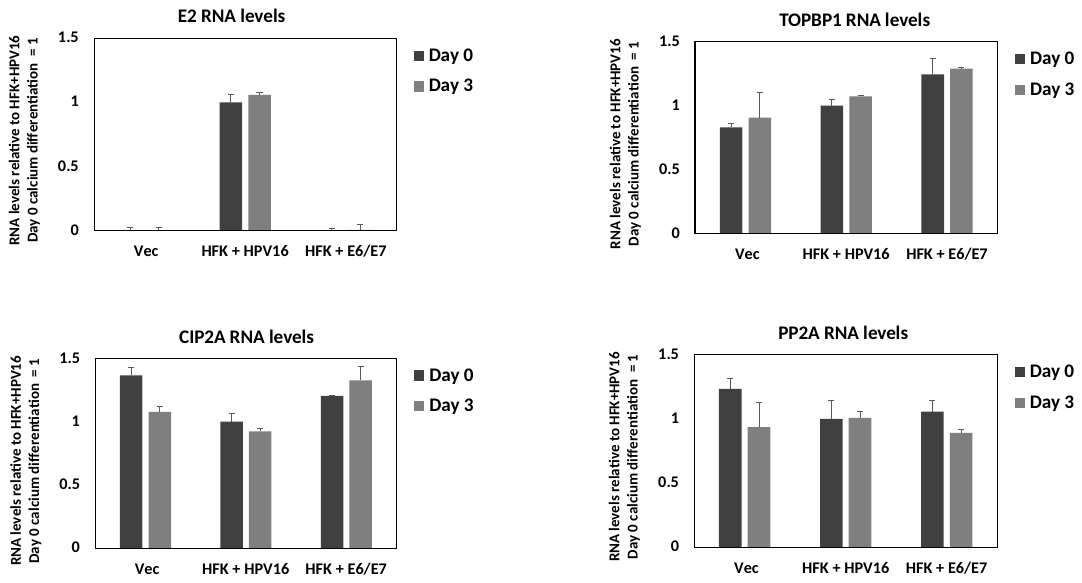
M

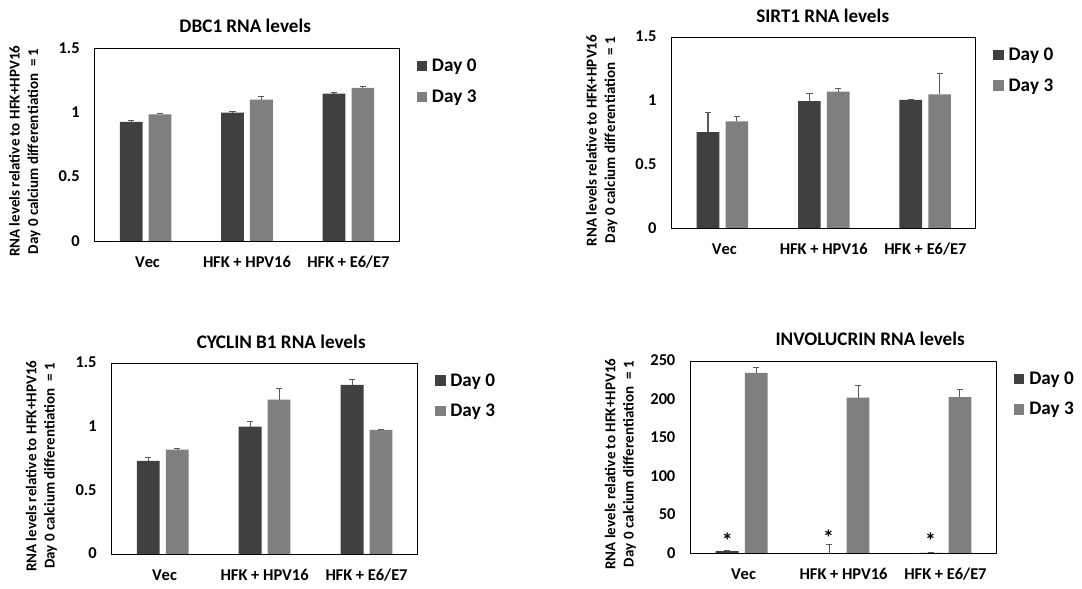

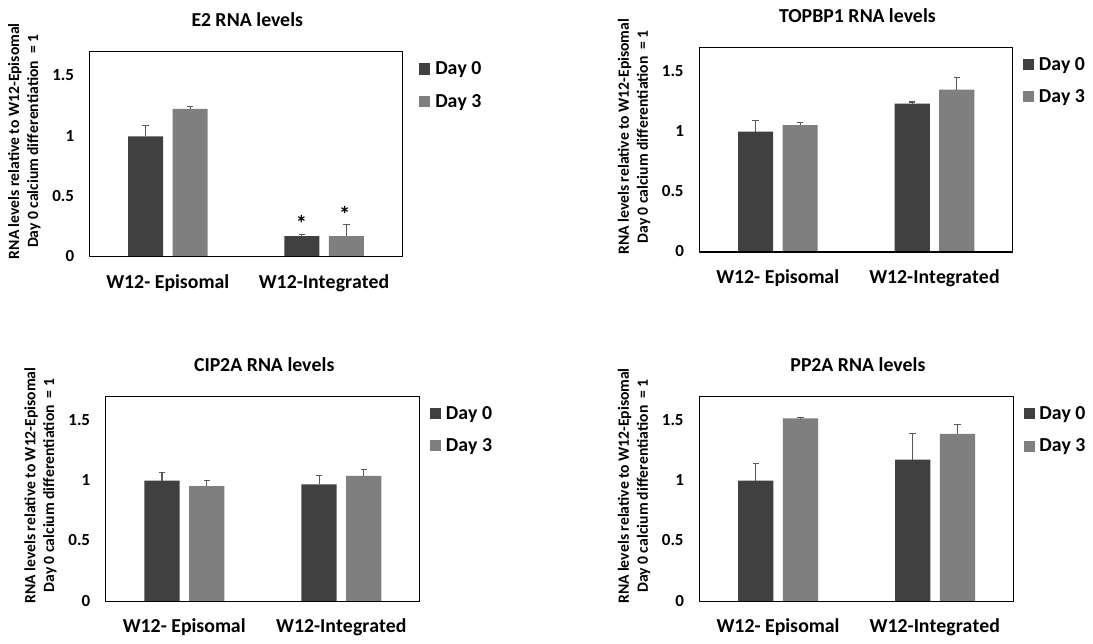
N

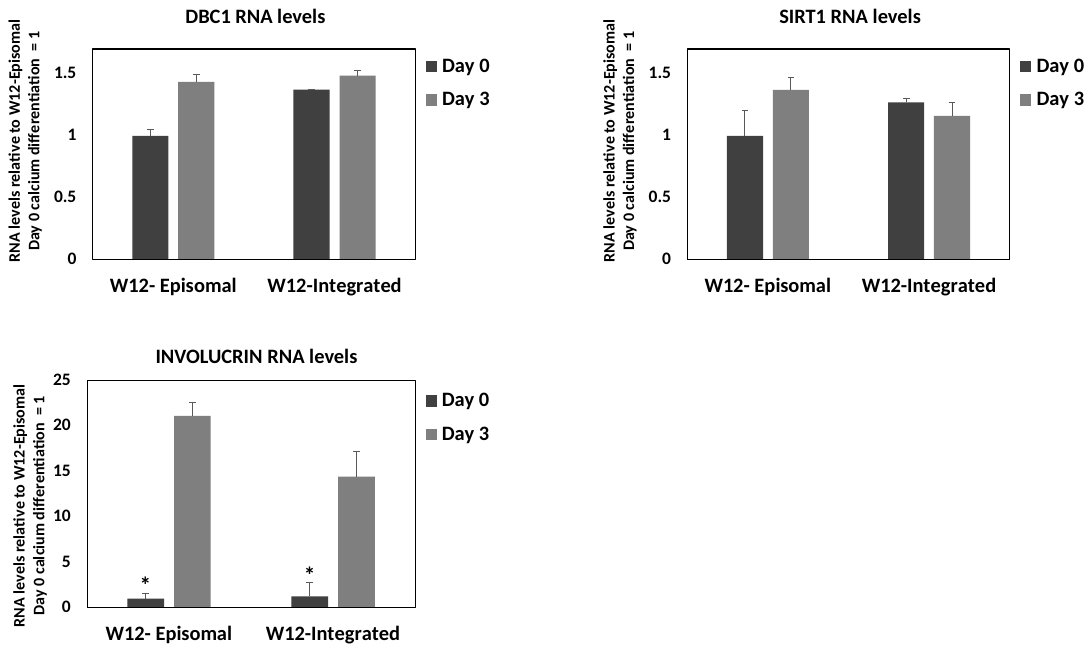

Figure S2. A. The experiment in Figure 2B was repeated and densitometry used to determine the levels of proteins relative to N/Tert-1+Vec levels in growing cells equaling 1, apart from E2 where the levels in N/Tert-1+E2-WT in growing cells was set as 1.B. The immunoprecipitation experiments described in Figures 2C-E were repeated with different cell extracts and the results quantitated by densitometry. C. The western blots from the organotypic rafts in Figure 2F were repeated and densitometry used to determine the levels of proteins relative to N/Tert-1+Vec levels equaling 1, apart from E2 where the levels in HFK+HPV16 was set as 1. D. The immunoprecipitation experiments described in Figures 2H-I were repeated extracts and the results quantitated by densitometry. E and F. The cells used for the organotypic raft proteins were differentiated using calcium and the indicated proteins detected using western blotting. G, H, and I. Immunoprecipitation experiments demonstrated the disruption of the TOPBP1-CIP2A interaction in differentiating cells (G), enhanced SIRT1-DBC1 interaction (H), and enhanced acetylation of viral and host proteins (I) in calcium differentiated HFK+HPV16, but not in HFK+E6/E7. J. The experiment in Figure 2J was repeated and the results quantitated. K. The experiments in Figure 2K-M (W12 cells) were repeated and quantitated. L. RNA was prepared from the indicated N/Tert-1 cells from two individual experiments and RNA levels for the indicated genes are shown relative to the levels in N/Tert-1+E2 Day 0 equaling 1. M. RNA was prepared from the indicated cell lines from two individual experiments and RNA levels for the indicated genes are shown relative to HFK+HPV16 Day 0 cells equaling 1. N. RNA was prepared from the indicated W12 cells from two individual experiments and RNA levels for the indicated genes shown relative to W12e Day 0 equaling 1. * Indicates a significant difference in protein levels from the growing samples (S2A) or the Day 0 samples (S2B and S2H), p-value < 0.05.

Figure S3.

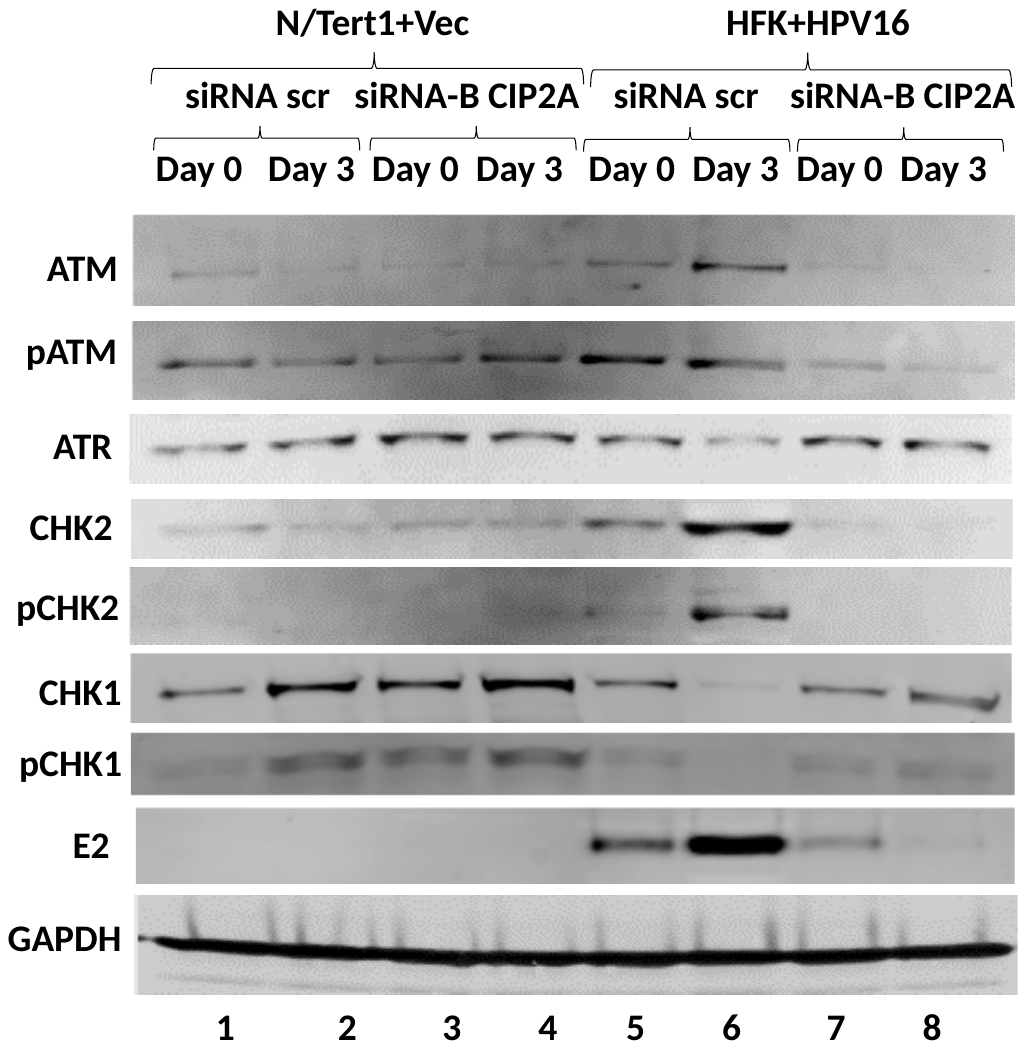
A

B

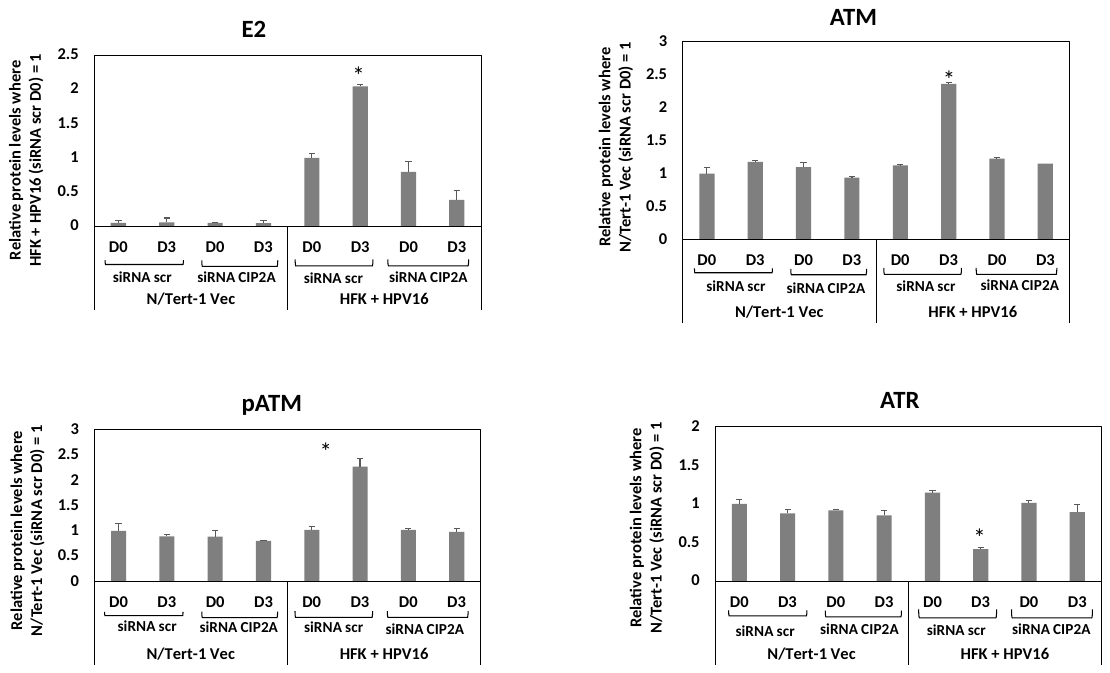

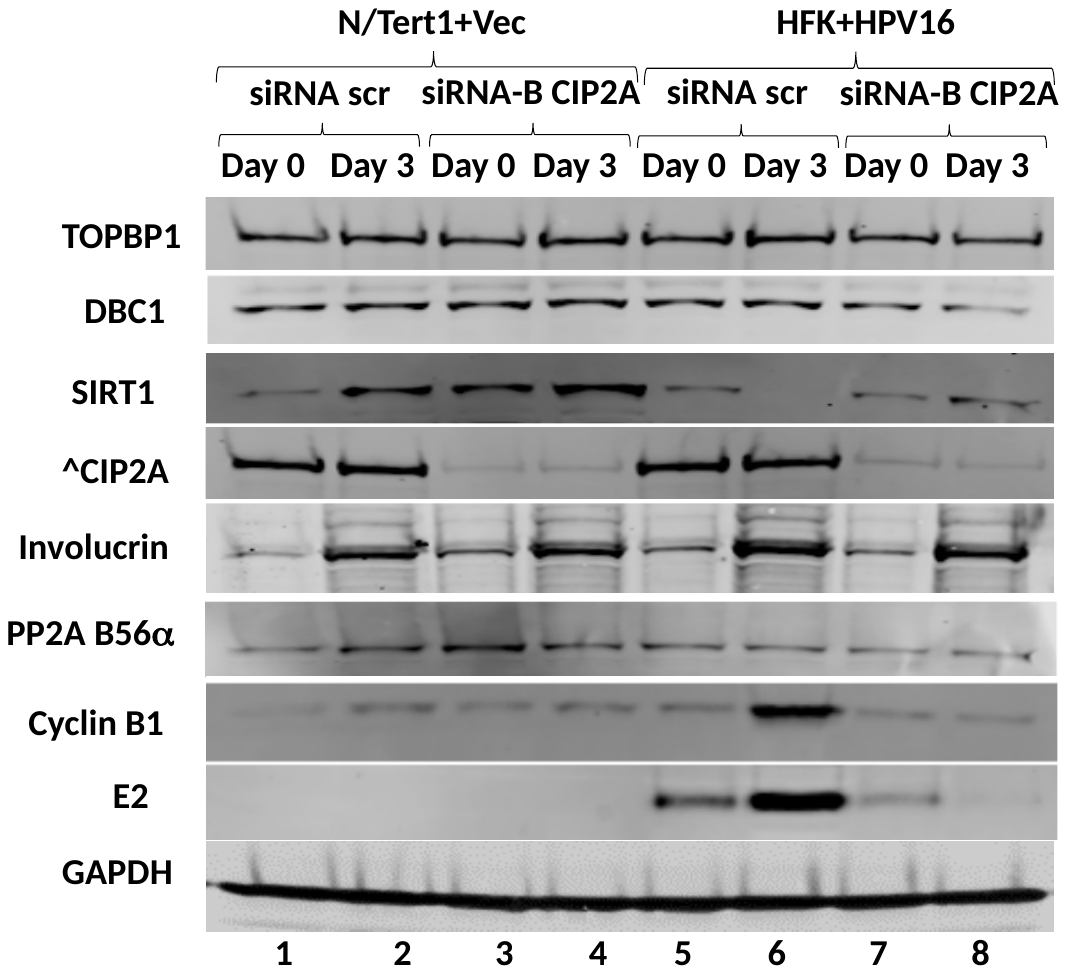

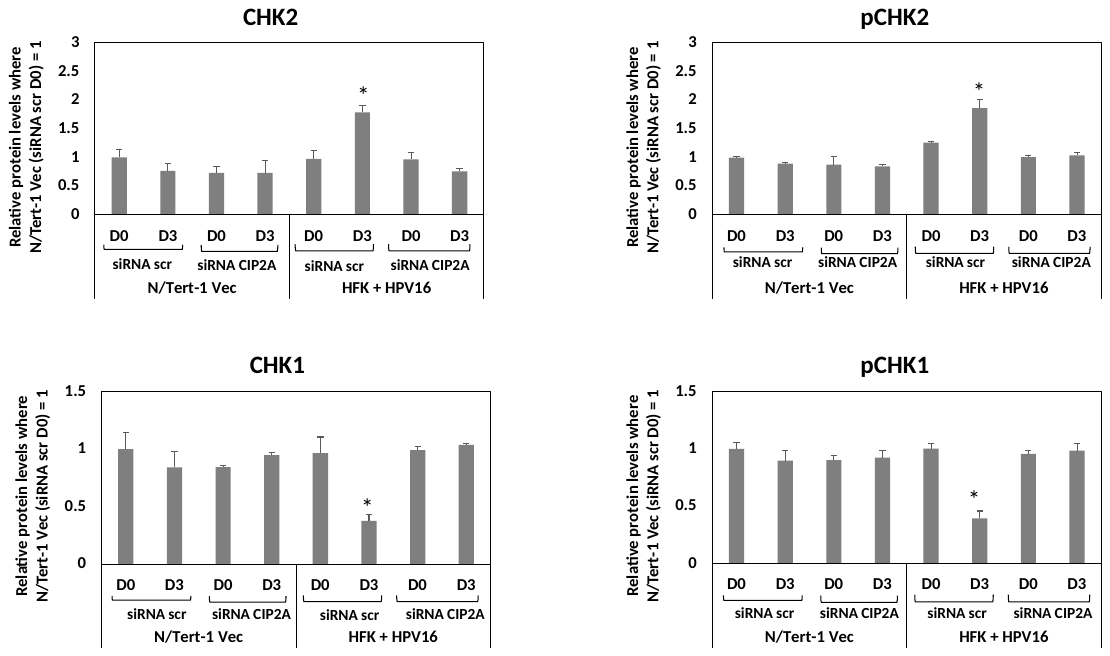
C

D

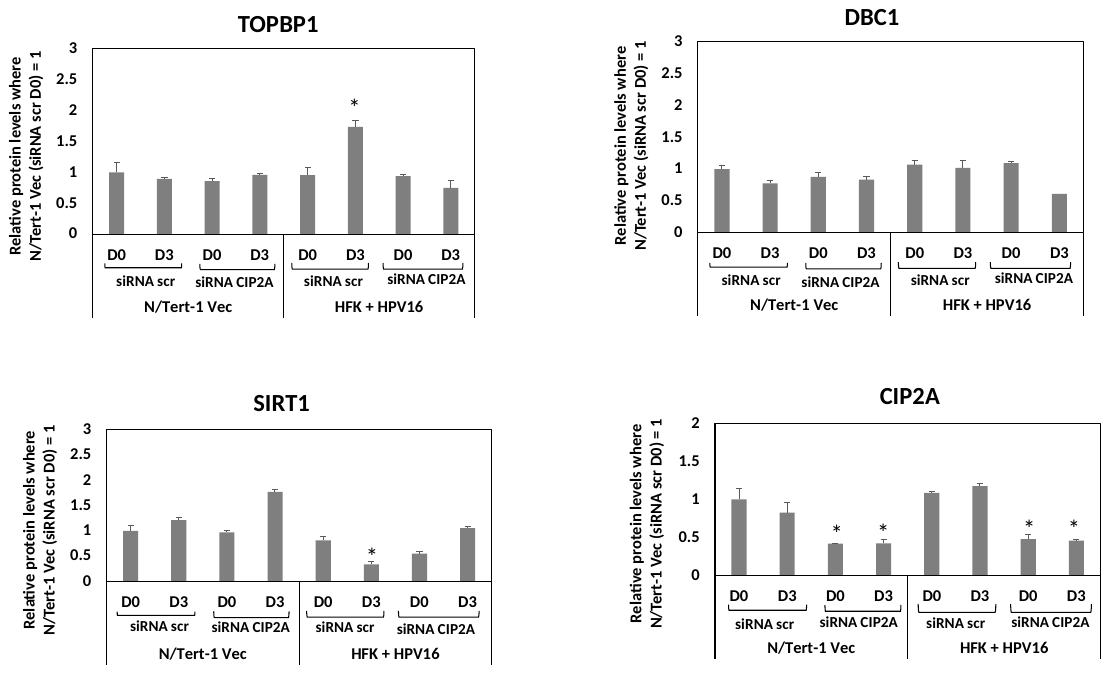

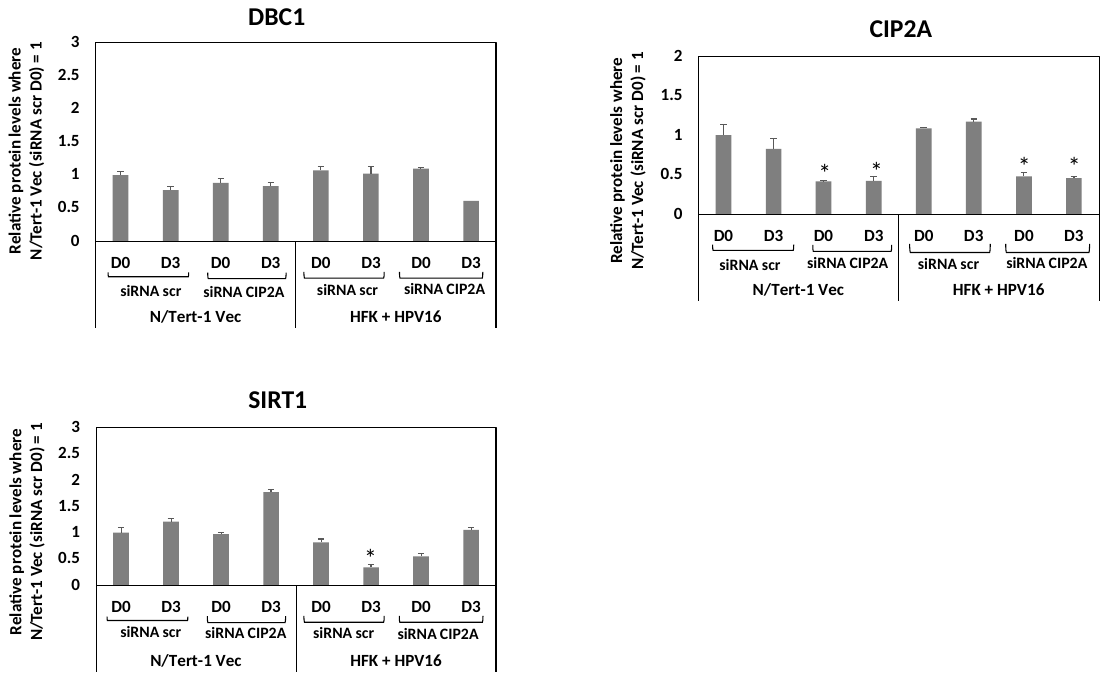

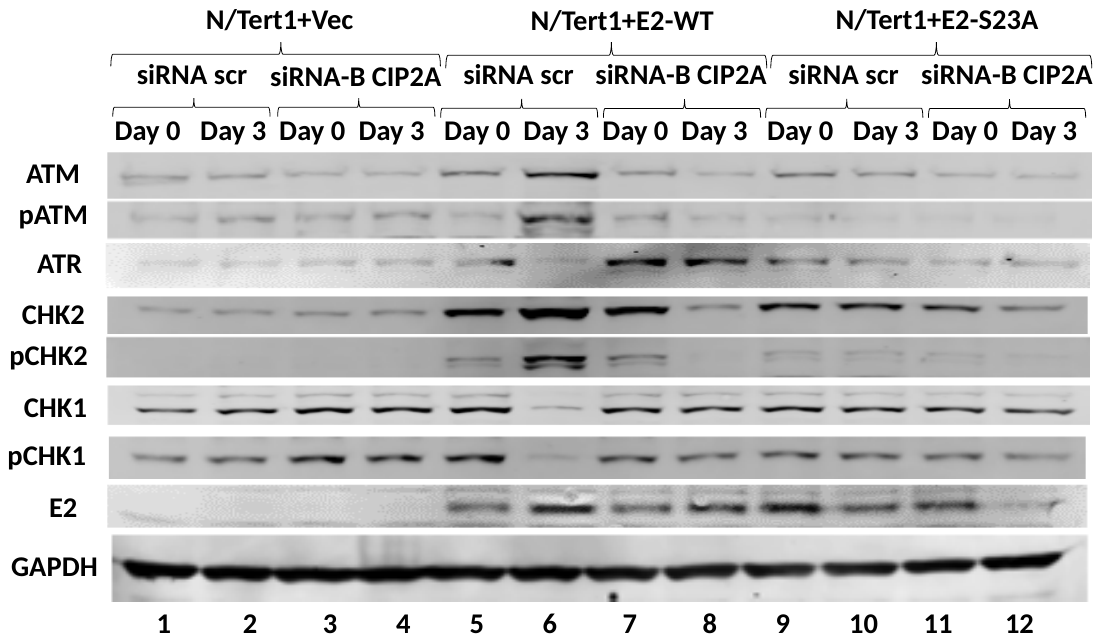
E

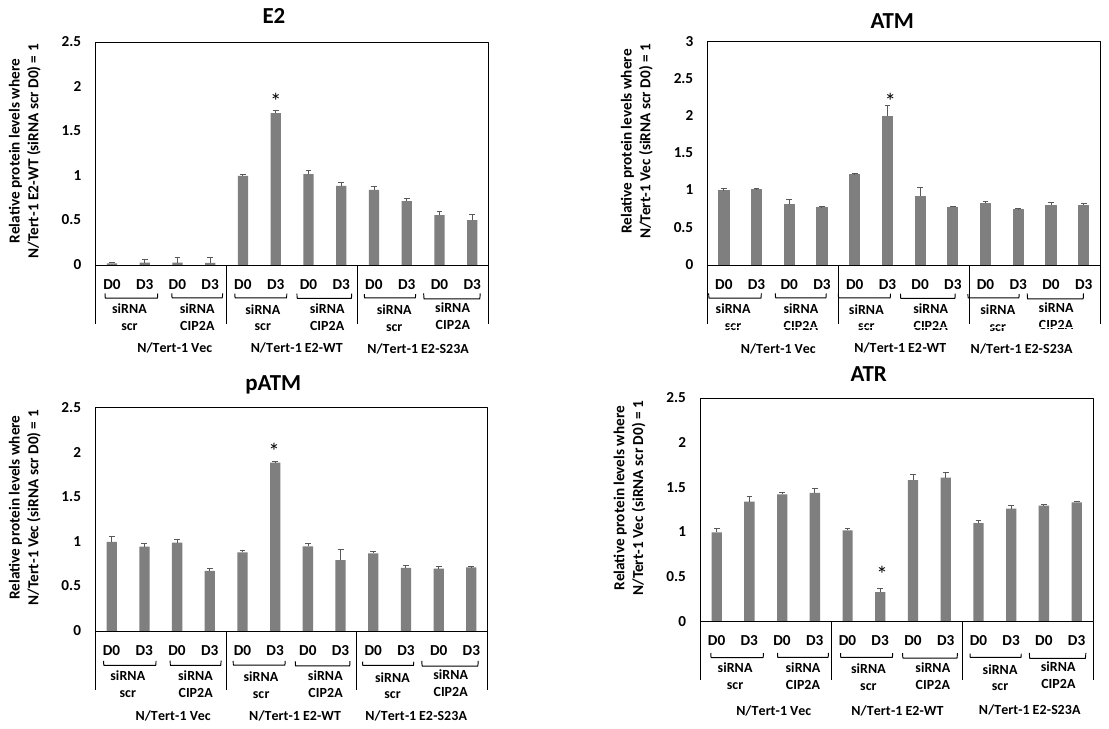
F

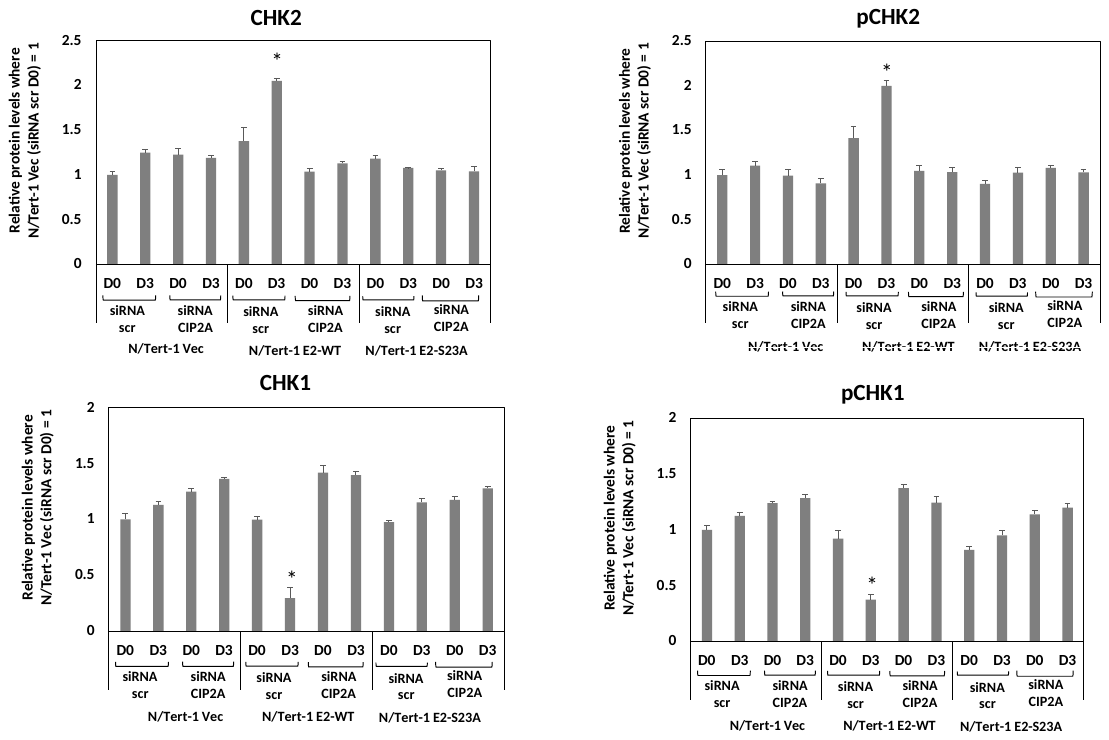

G

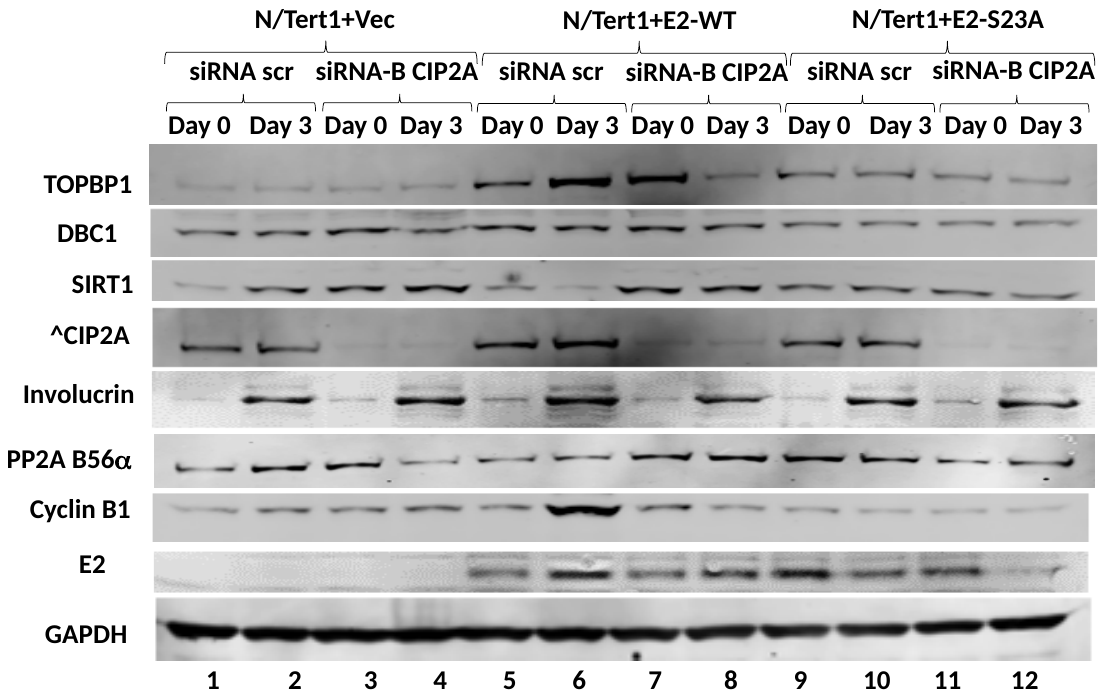

H

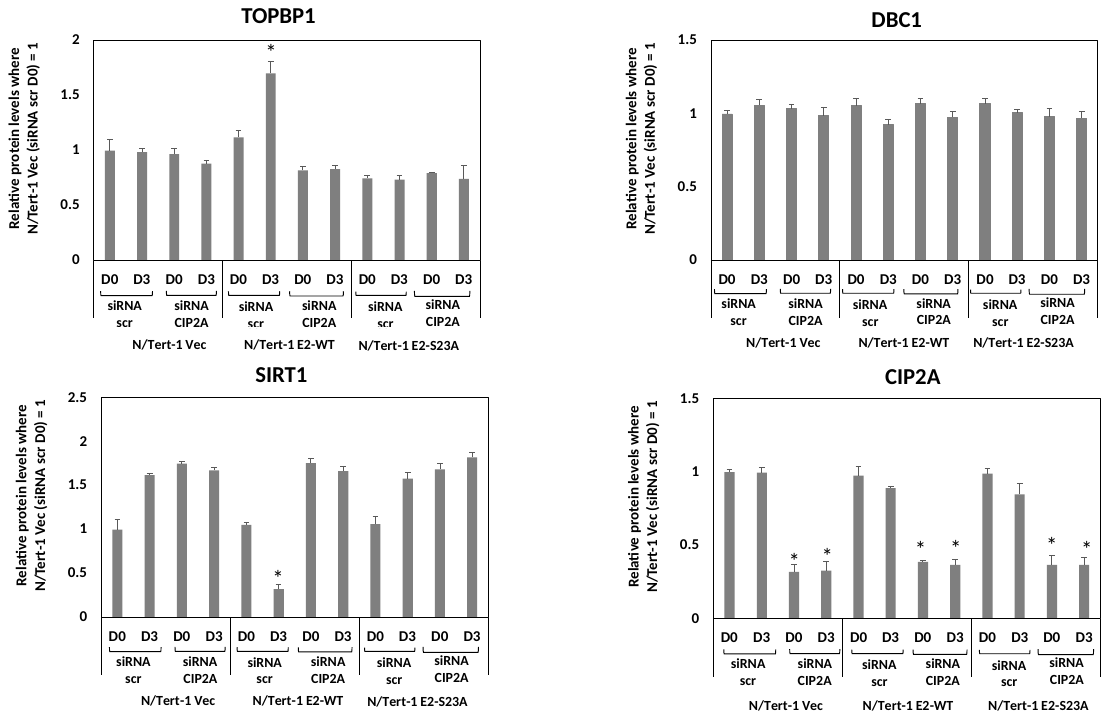

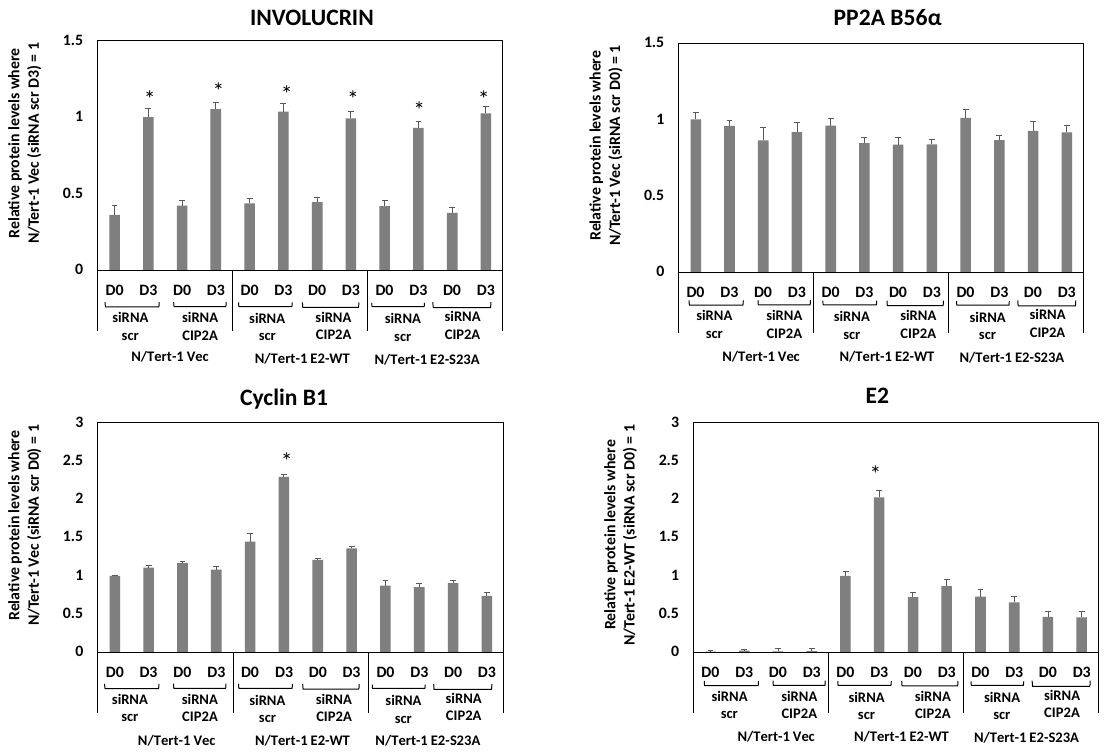

I

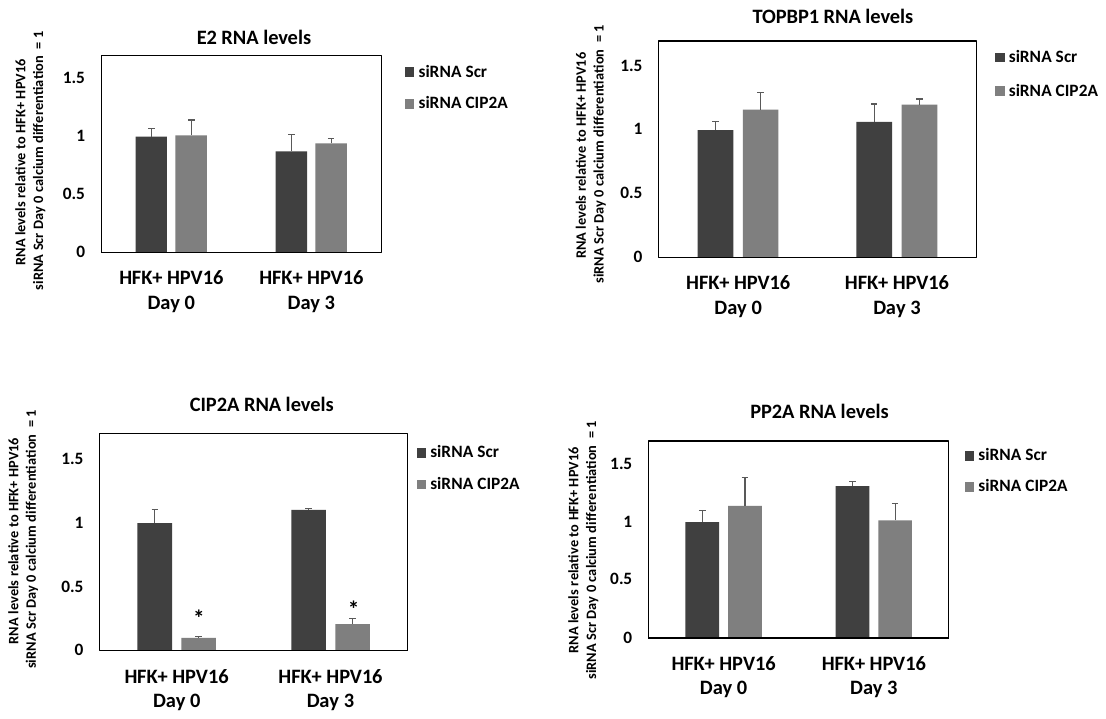

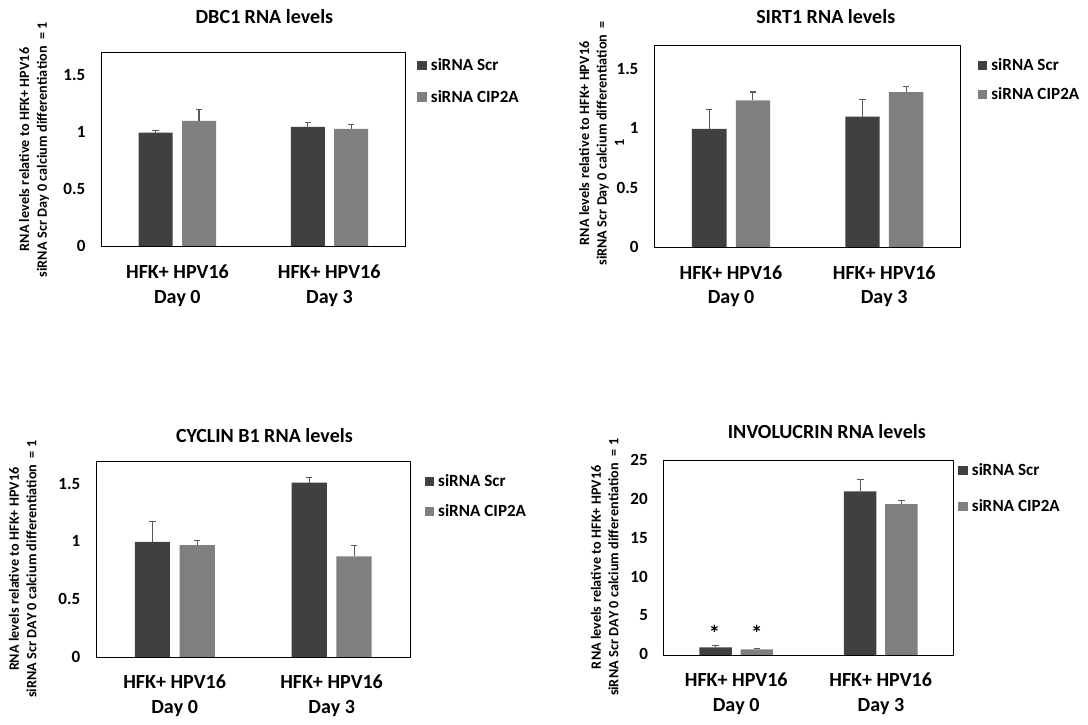

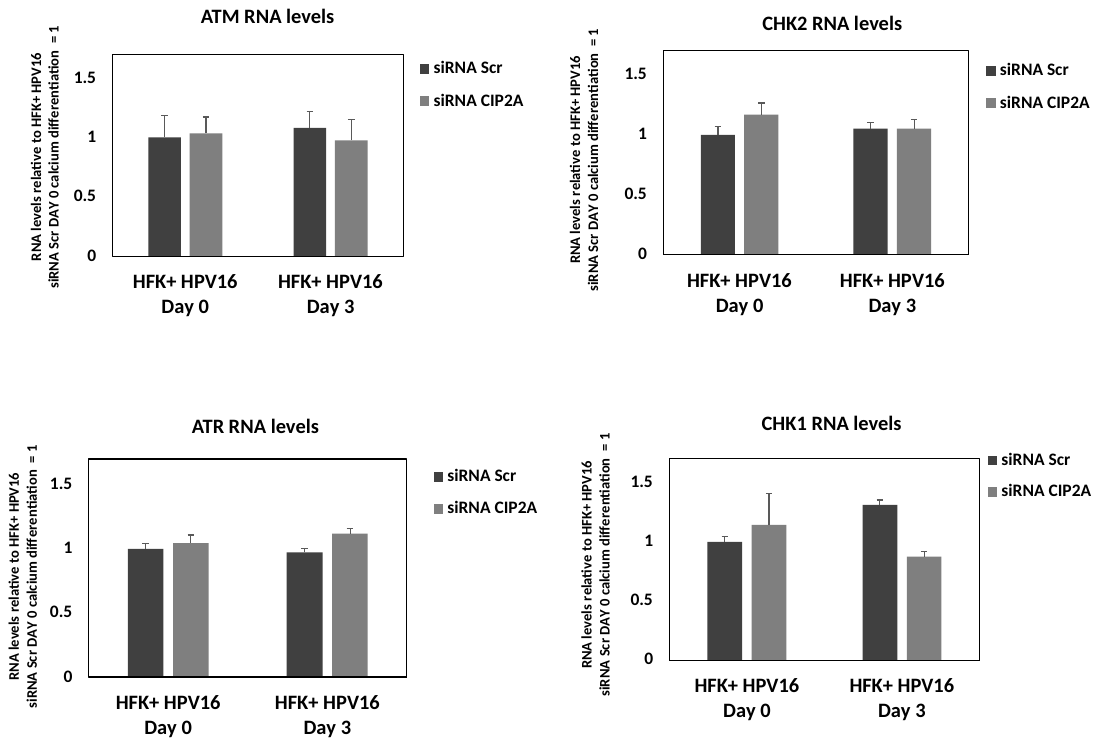

J

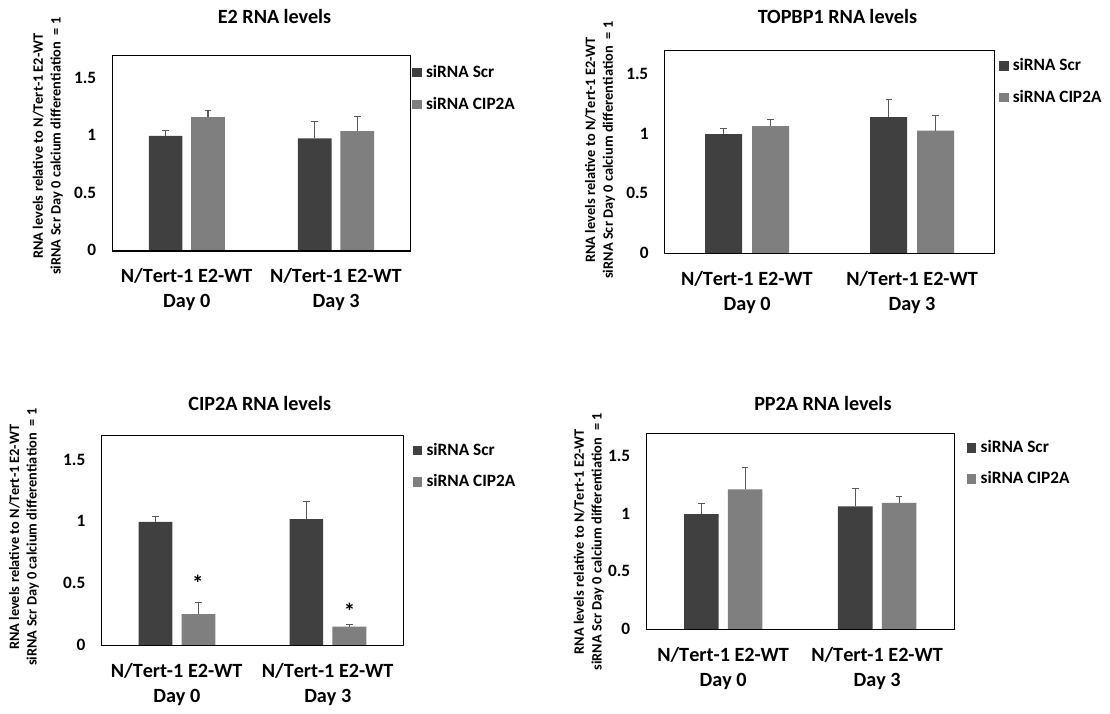

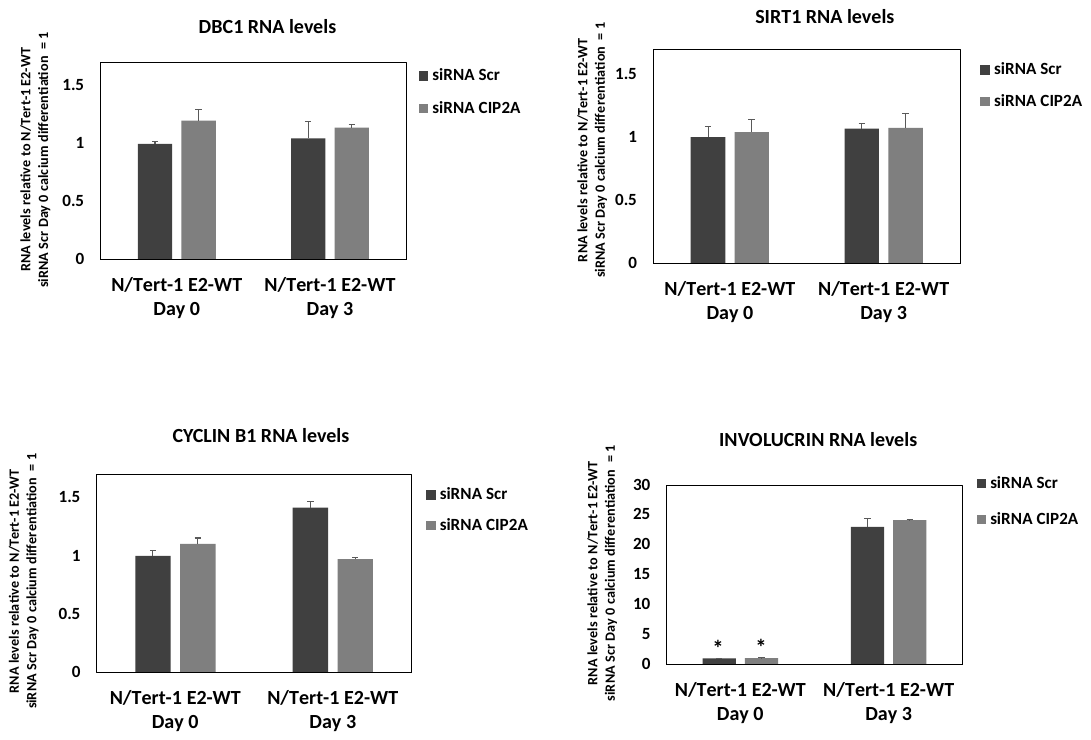

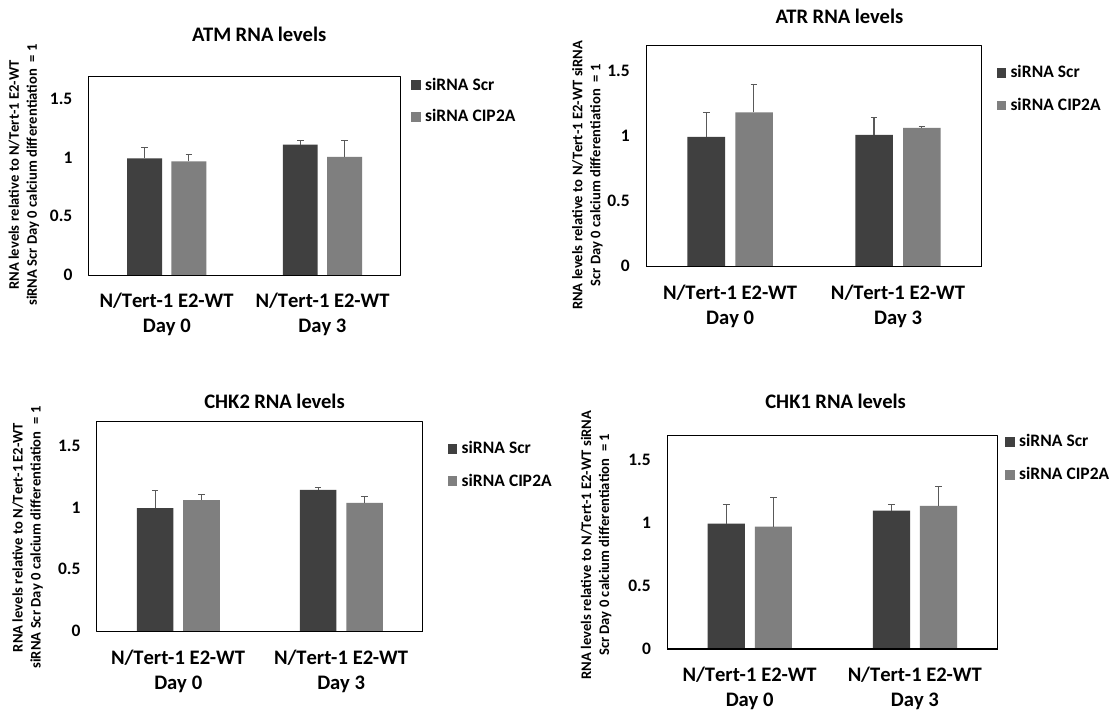

K

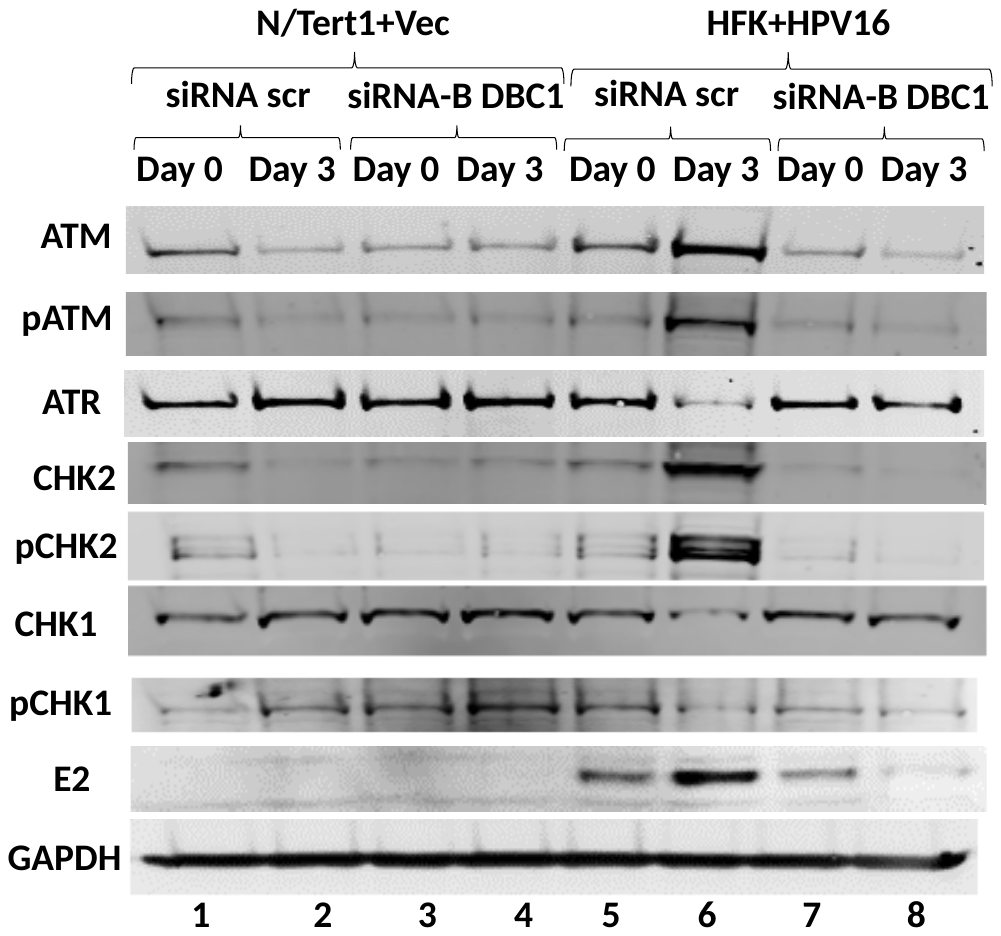

L

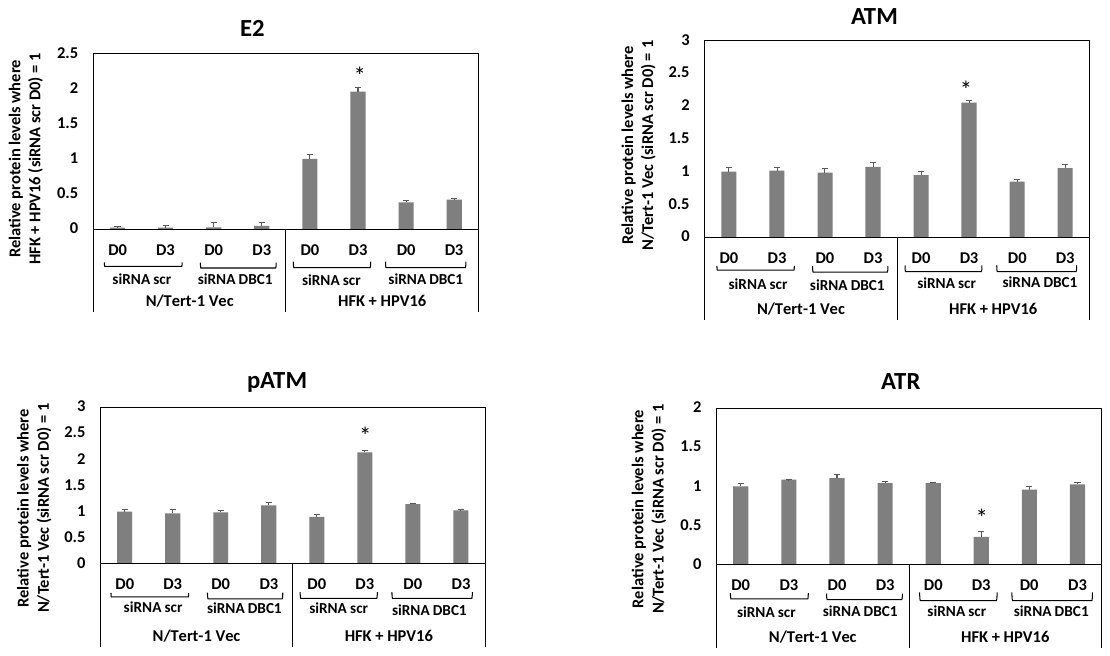

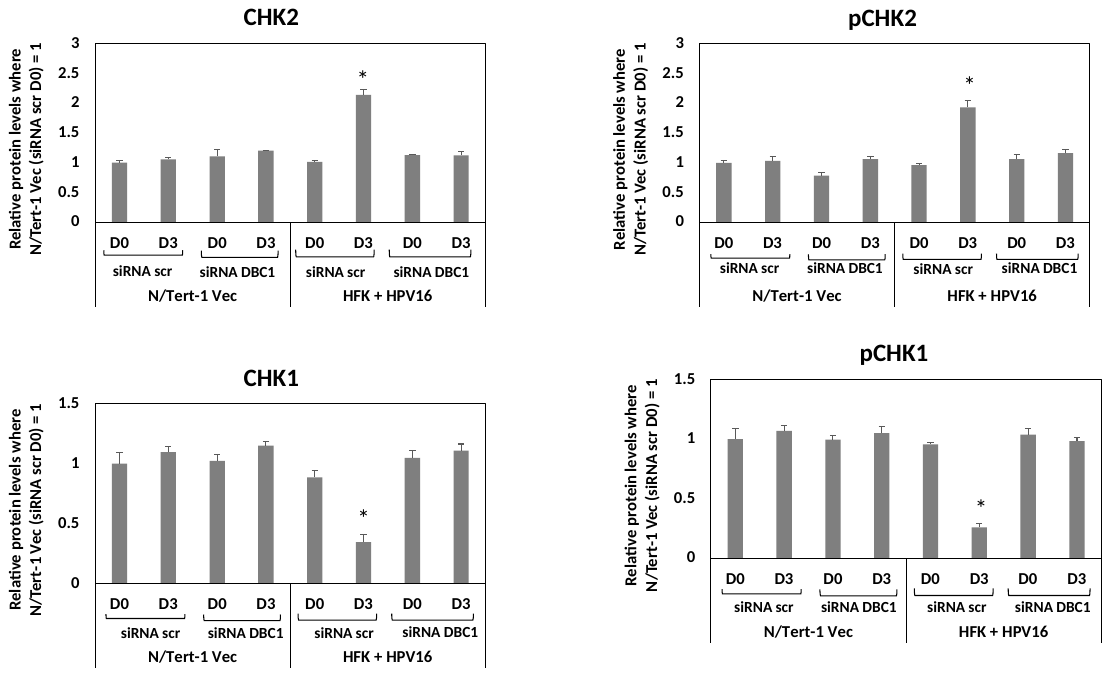

M

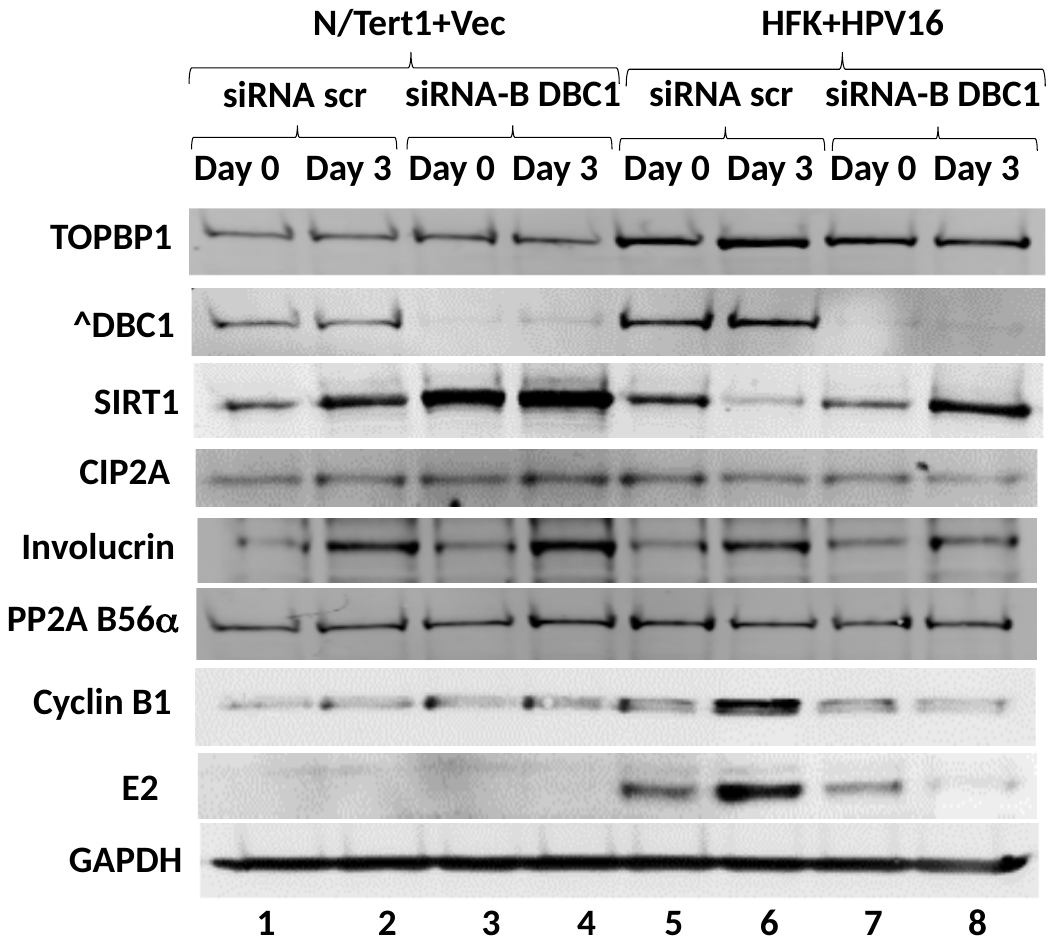

N

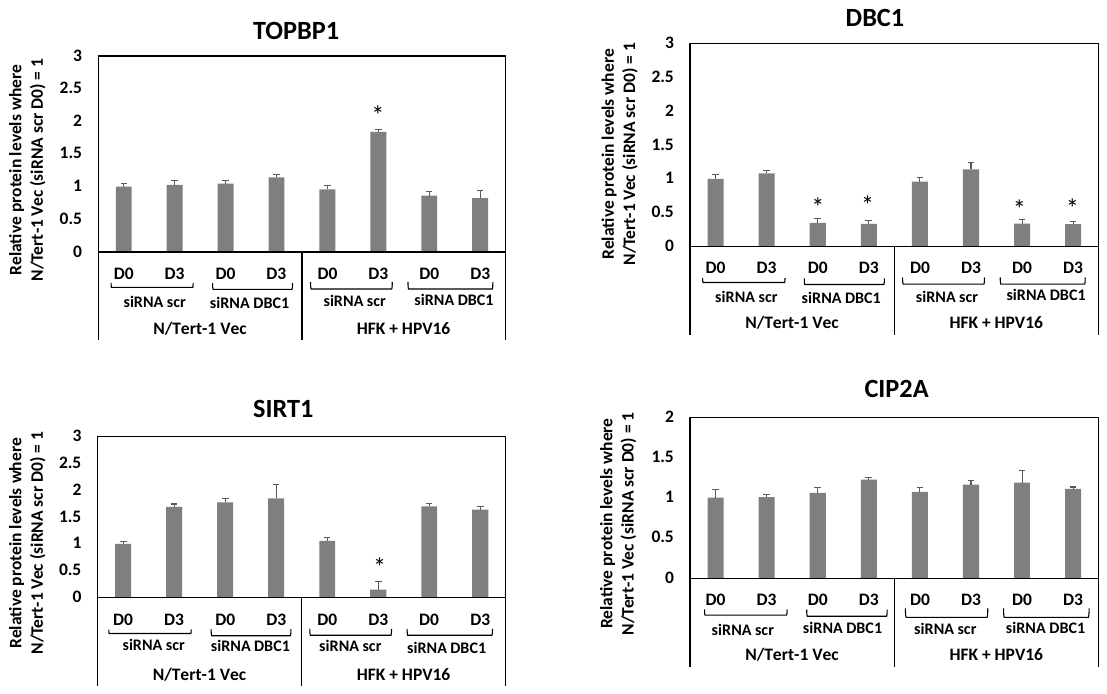

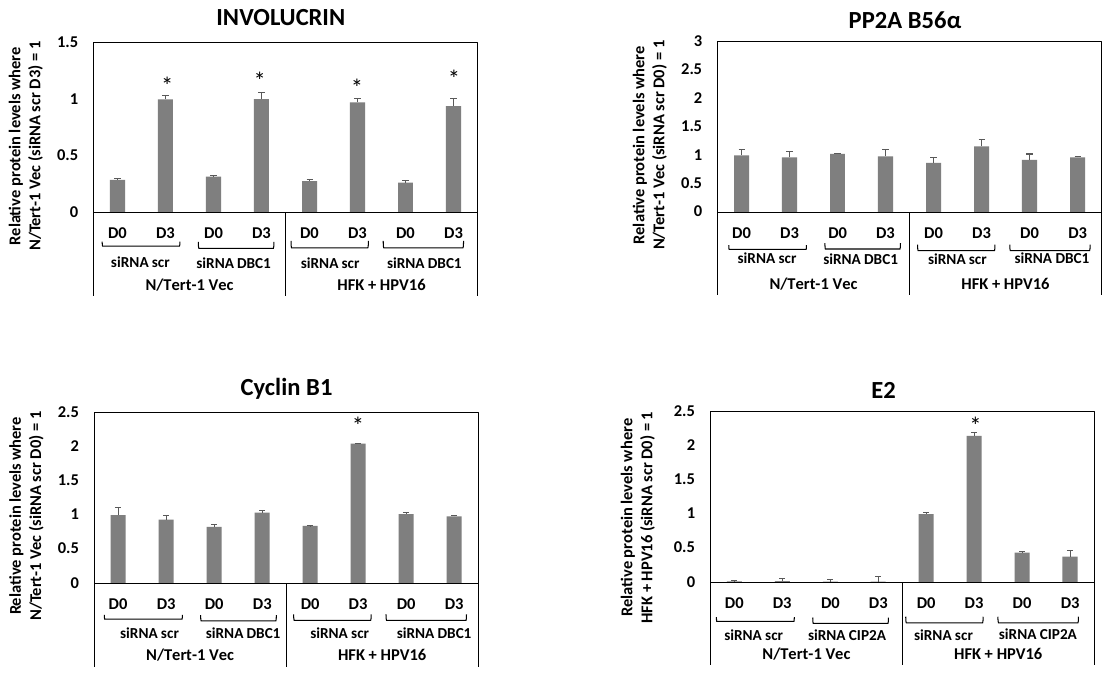

O

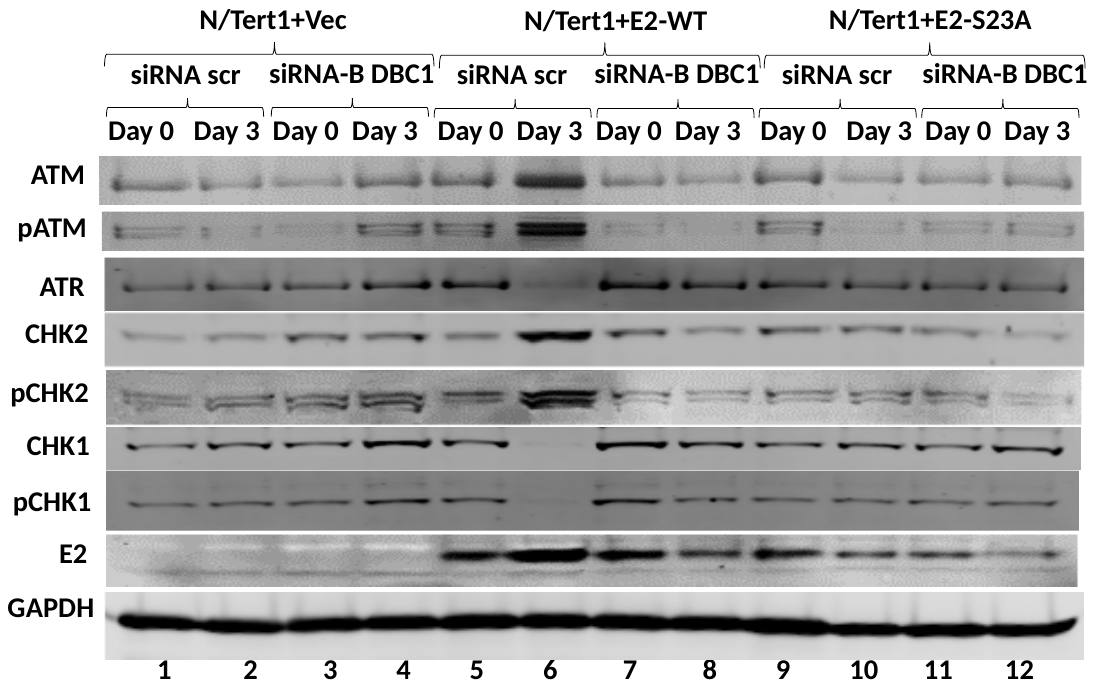

P

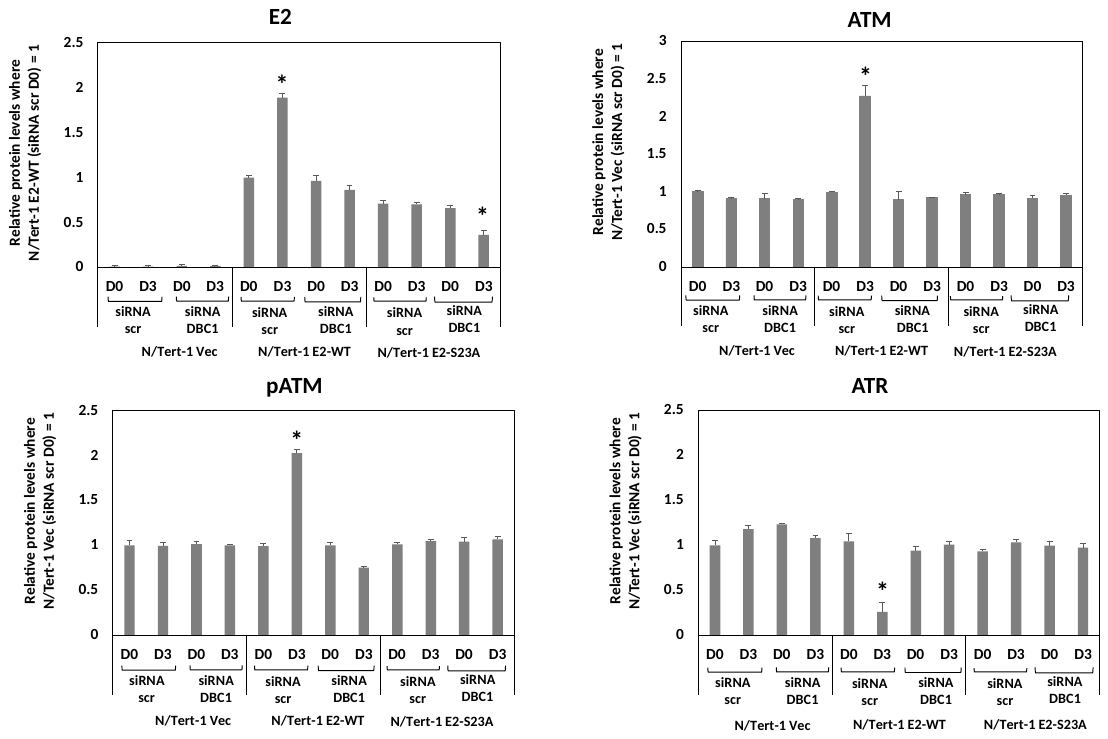

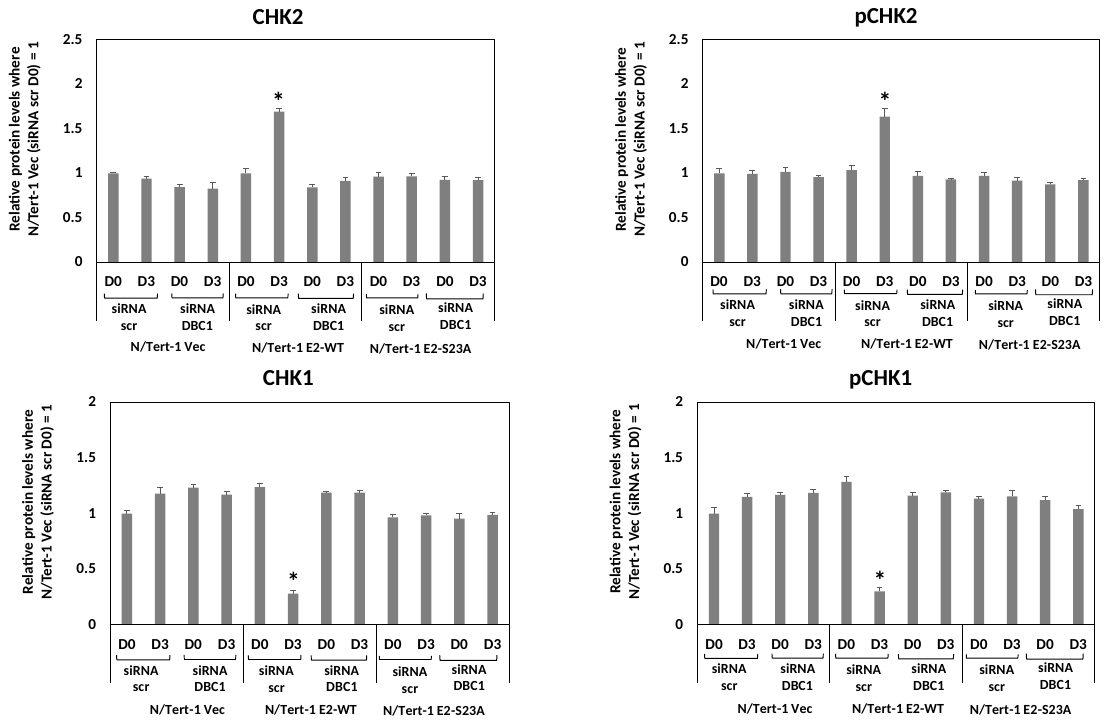

Q

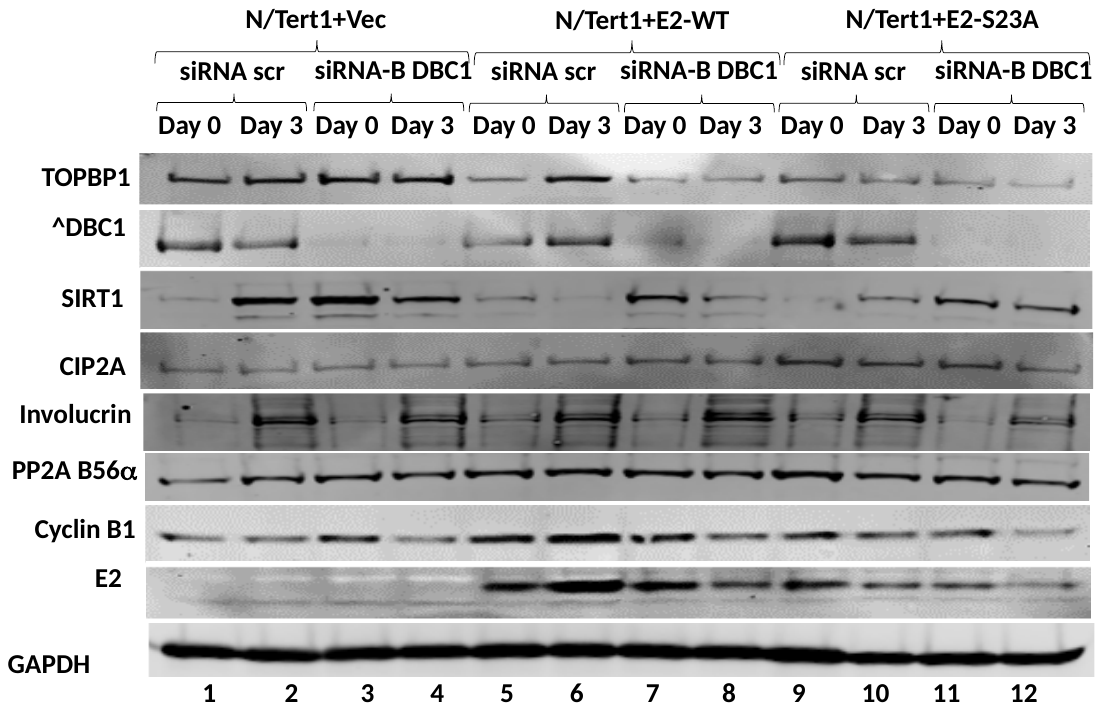

R

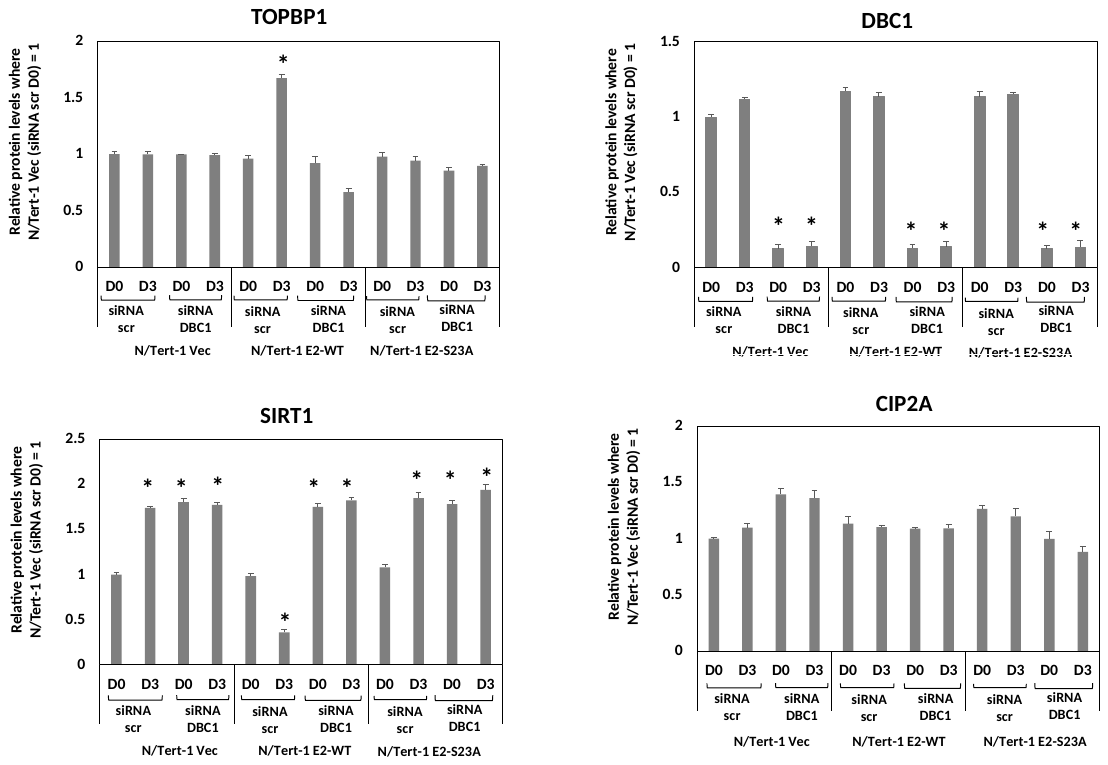

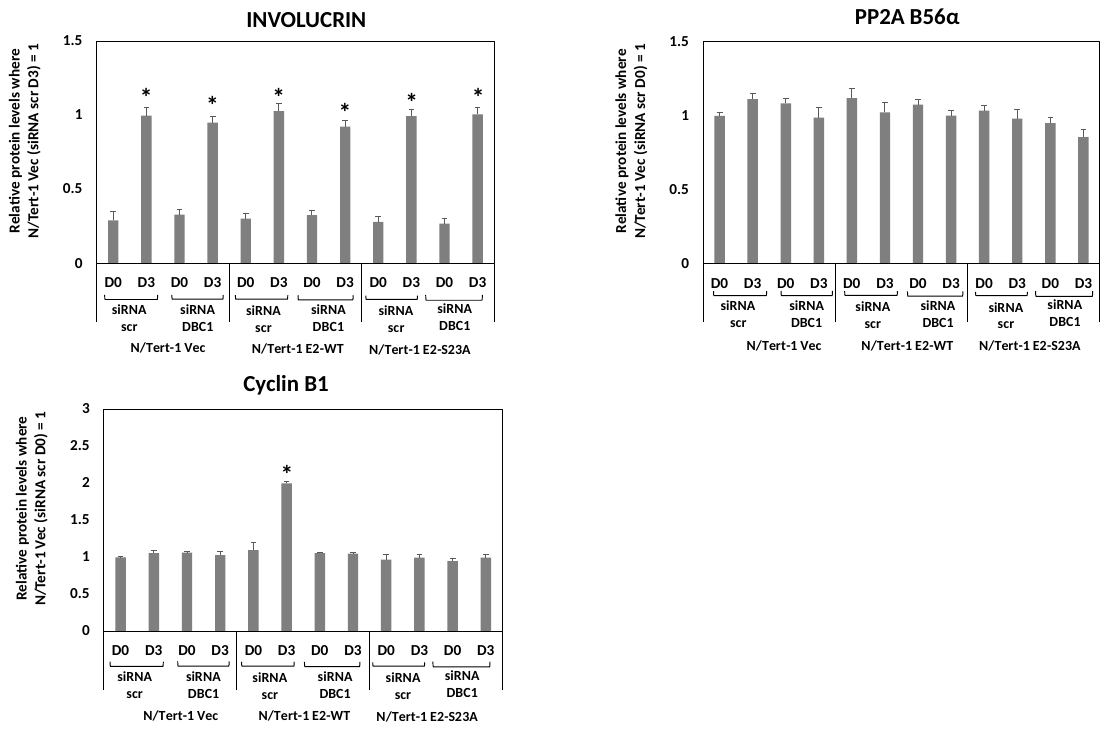

S

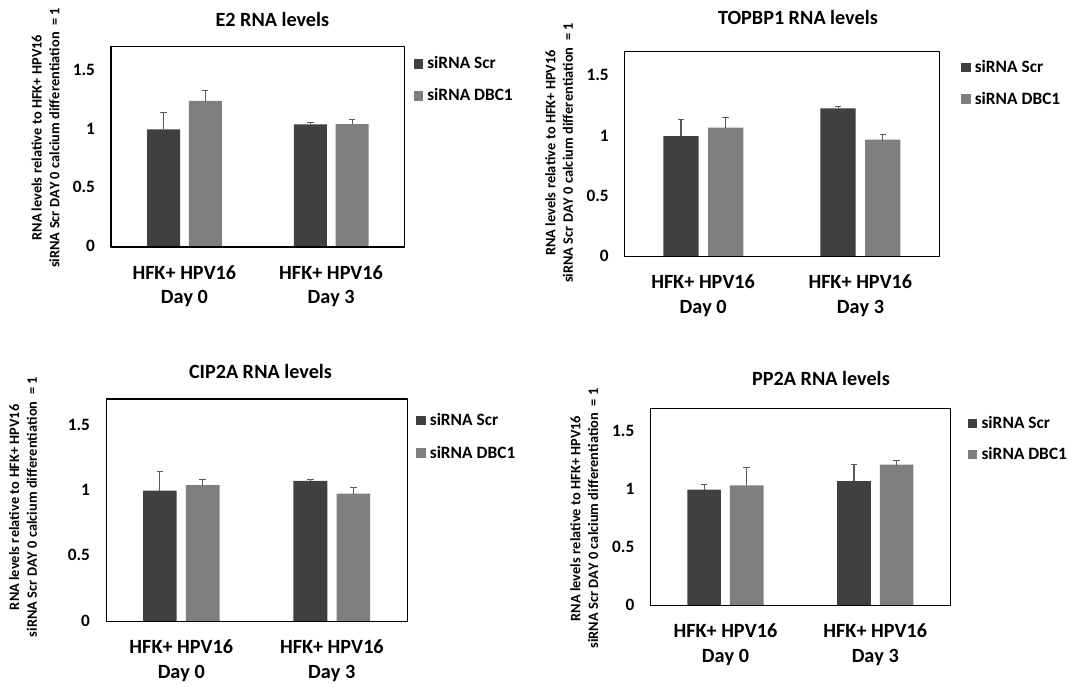

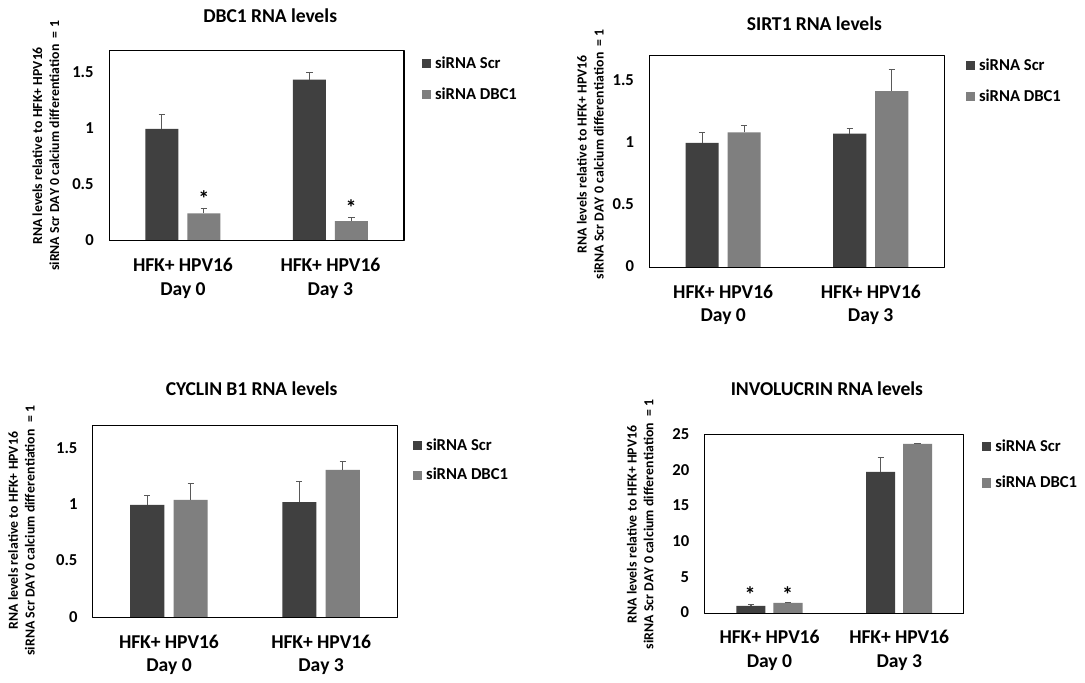

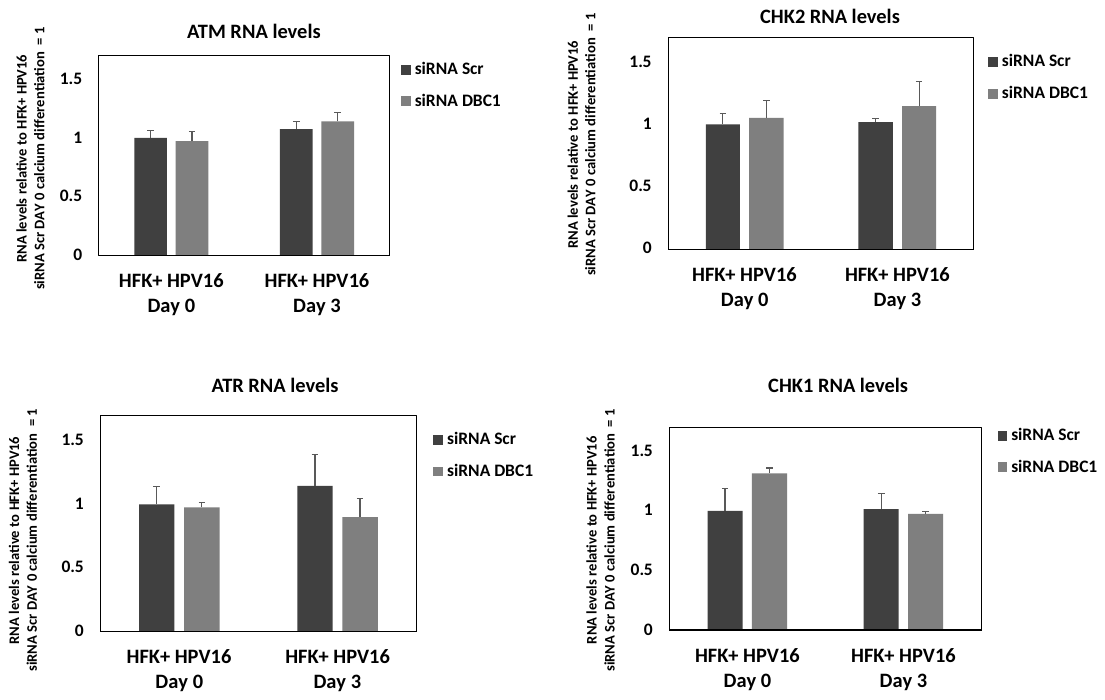

T

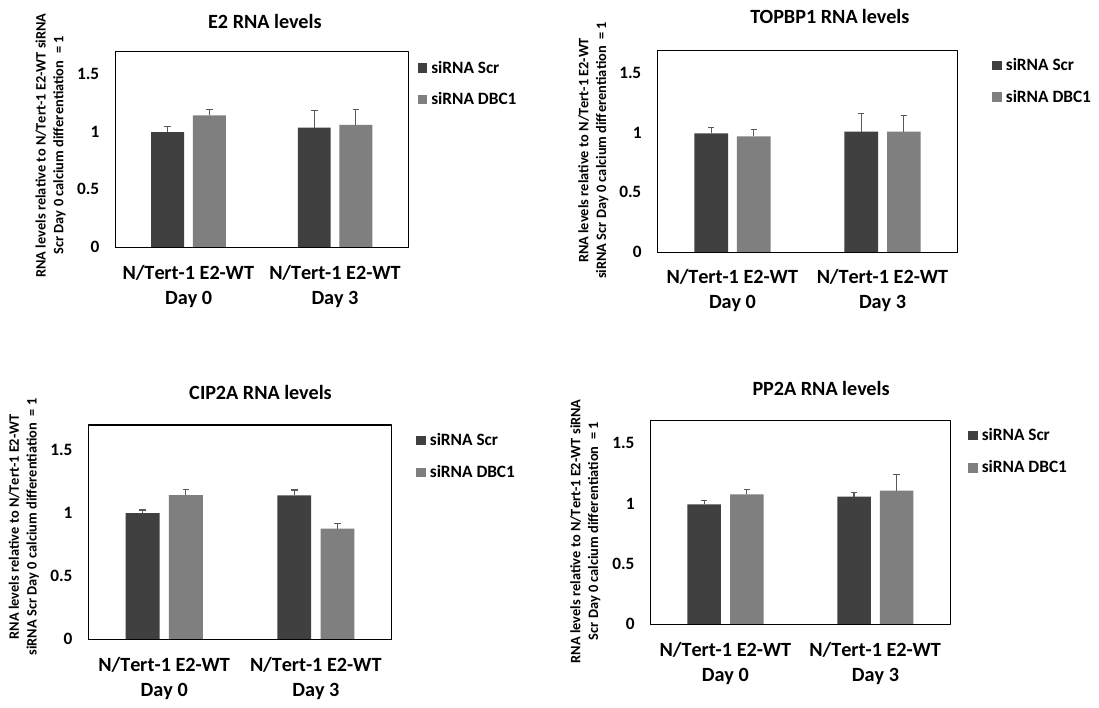

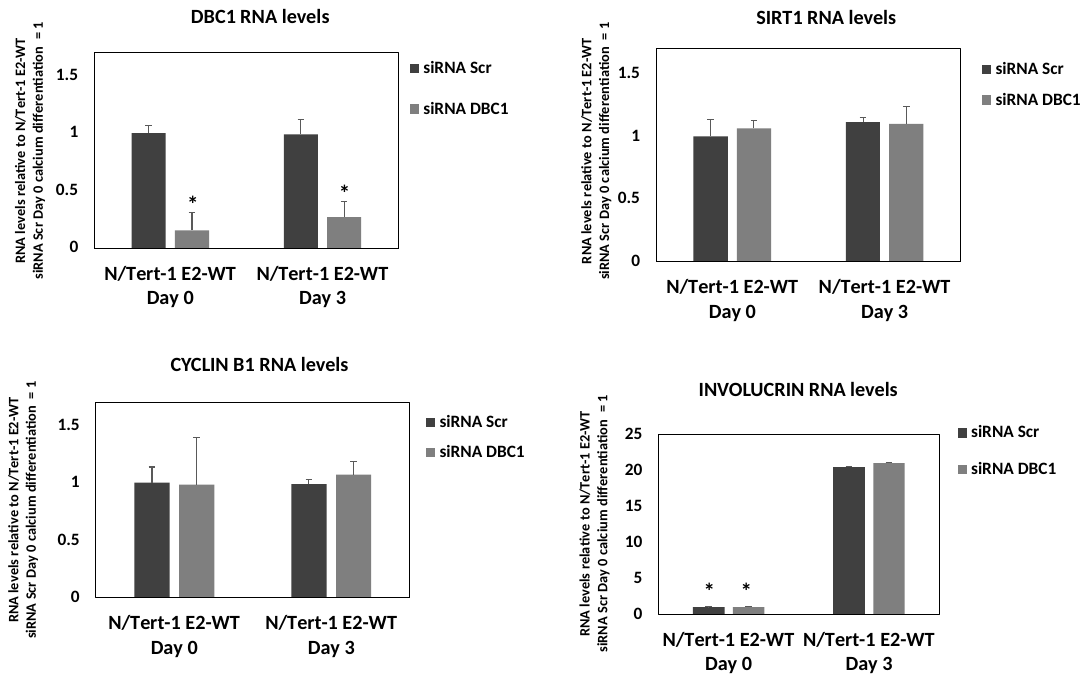

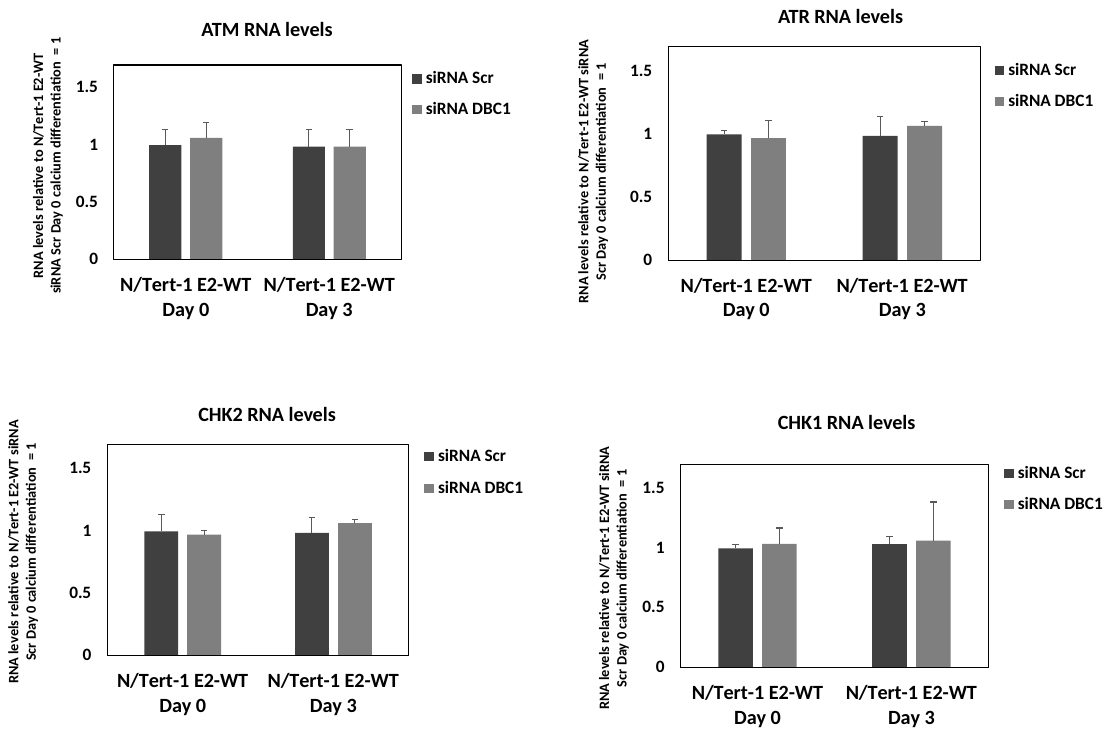

Figure S3. A. The experiment in Figure 3D was repeated with a different CIP2A siRNA, which also eliminated the “ATM up ATR down” phenotype. B. The results from Figures 3D and S3A were quantitated. C. The experiment in Figure 3E was repeated with a different CIP2A siRNA which also prevented the E2 reprogramming following differentiation. D. The results from Figures 3E and S3C were quantitated. E. The experiment in Figure 3F was repeated with a different CIP2A siRNA, which also eliminated the “ATM up ATR down” phenotype. F. The results from Figures 3F and S3E were quantitated. G. The experiment in Figure 3G was repeated with a different CIP2A siRNA which also prevented the E2 reprogramming following differentiation. H. The results from Figures 3G and S3G were quantitated. I. Following CIP2A knockdown there were no changes in RNA expression levels of proteins under study (apart from CIP2A) when compared with control siRNA cells in HFK+HPV16 cells. J. Following CIP2A knockdown there were no changes in the RNA expression levels of proteins under study (apart from CIP2A) when compared with control siRNA cells in HFK+HPV16 or N/Tert-1+E2-ET cells. K. The experiment in Figure 3H was repeated with a different DBC1 siRNA, which also eliminated the “ATM up ATR down” phenotype. L. The results from Figures 3H and S3K were quantitated. M. The experiment in Figure 3I was repeated with a different DBC1 siRNA which also prevented the E2 programming following differentiation. N. The results from Figures 3I and S3M were quantitated. O. The experiment in Figure 3J was repeated with a different DBC1 siRNA, which also eliminated the “ATM up ATR down” phenotype. P. The results from Figures 3J and S3O were quantitated. Q. The experiment in Figure 3K was repeated with a different DBC1 siRNA which also prevented the E2 programming following differentiation. R. The results from Figures 3K and S3Q were quantitated. S. Following DBC1 knockdown there were no changes in the RNA expression levels of proteins under study (apart from DBC1) when compared with control siRNA cells in HFK+HPV16 or N/Tert-1+E2-ET cells. Significant changes in the histograms are indicated by *, p-value<0.05.

Figure S4

A	B

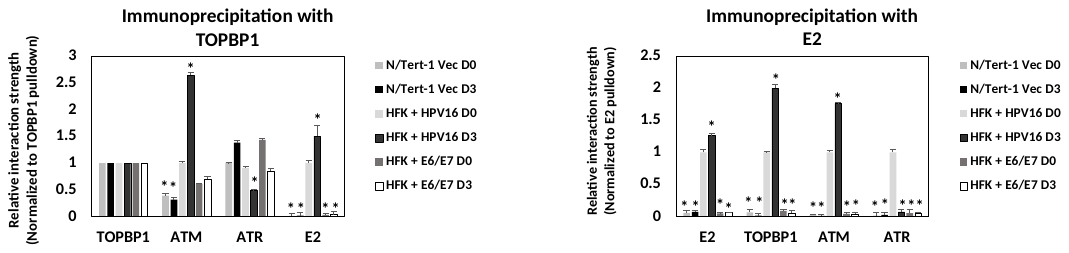

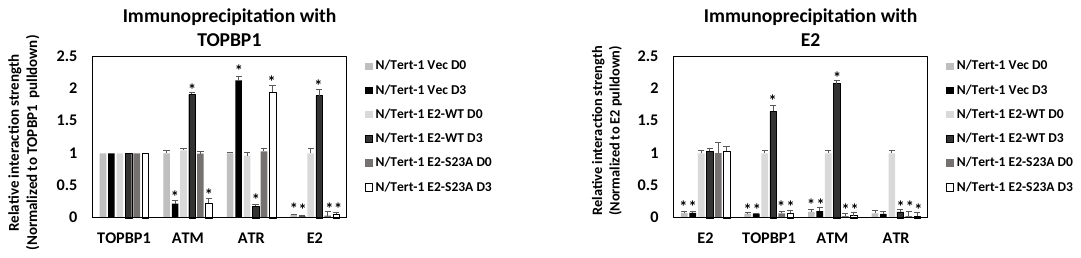

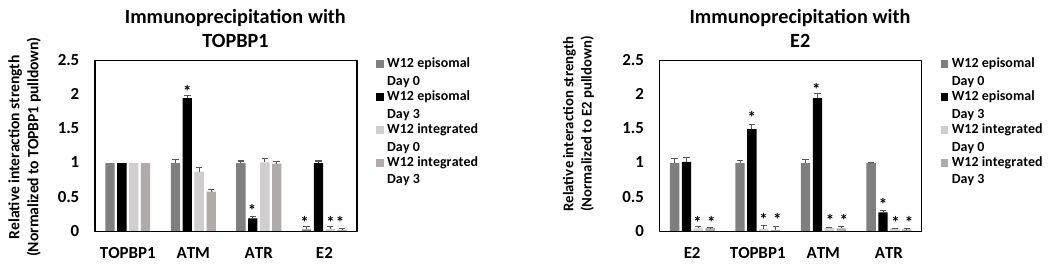

C

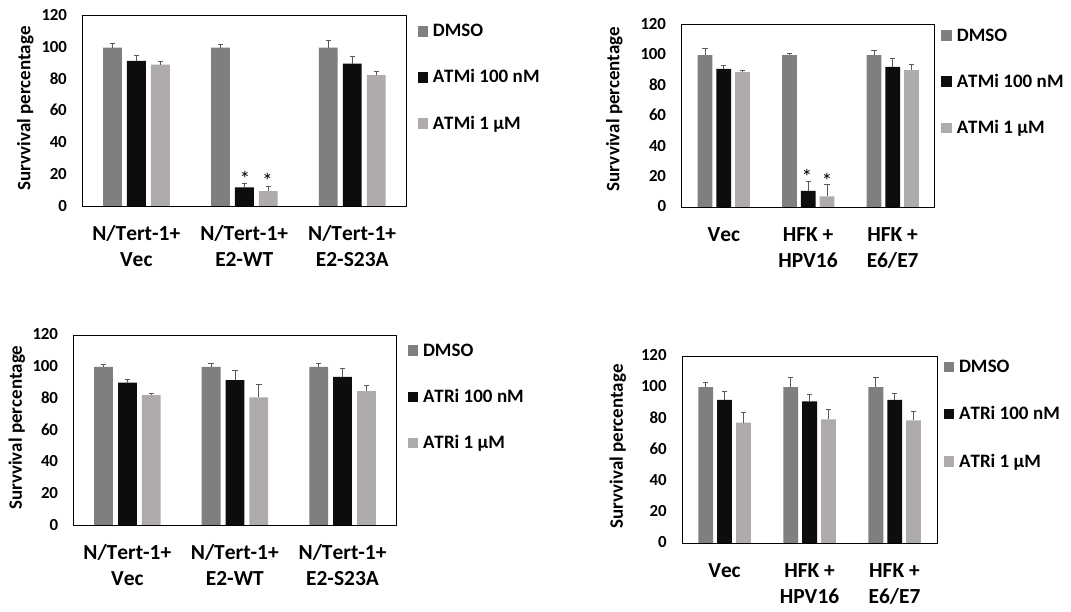

Figure S4. A. The immunoprecipitation experiments shown in Figure 4A were repeated and quantitated. B. The immunoprecipitation experiments shown in Figure 4B were repeated and quantitated. C. The experiments in Figures 4C and 4D were repeated and the results quantitated. Significant changes in the histograms are indicated by *, p-value<0.05.

Figure S5

A

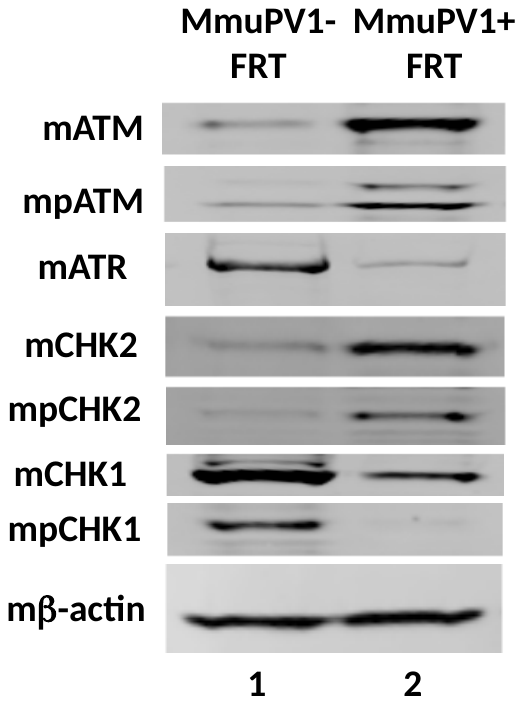

B

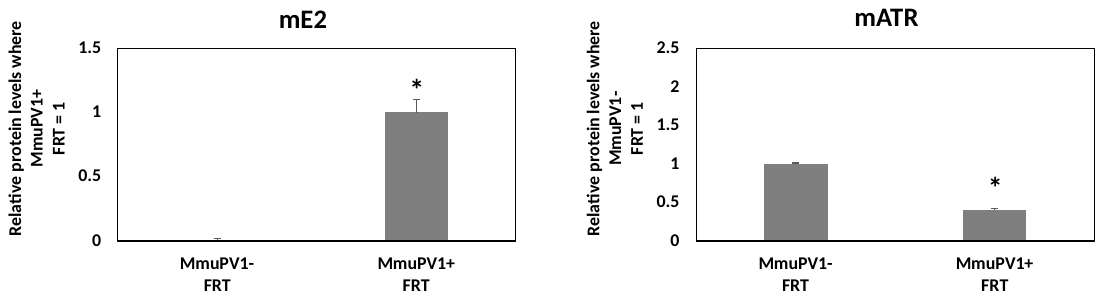

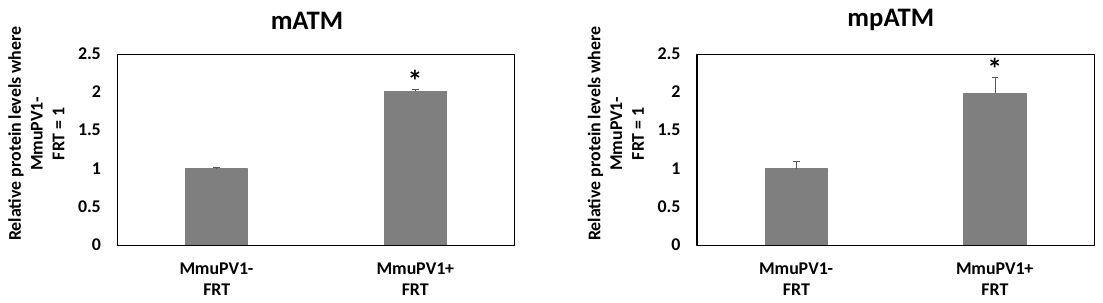

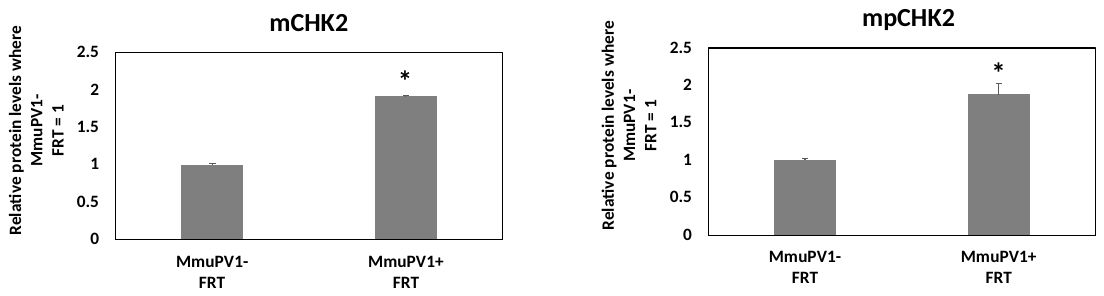

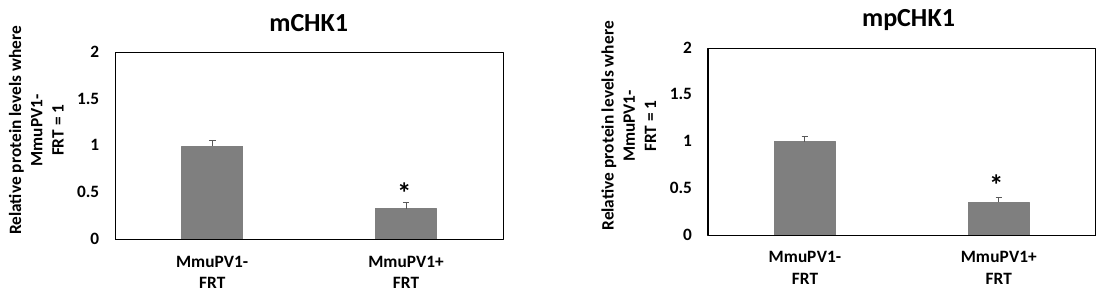

C

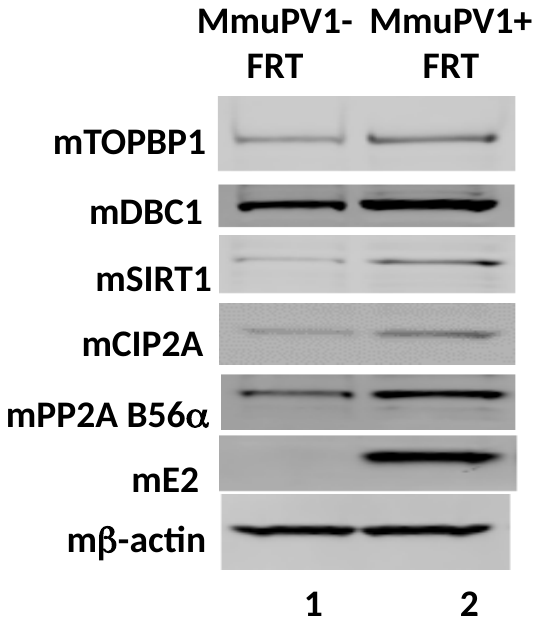

D

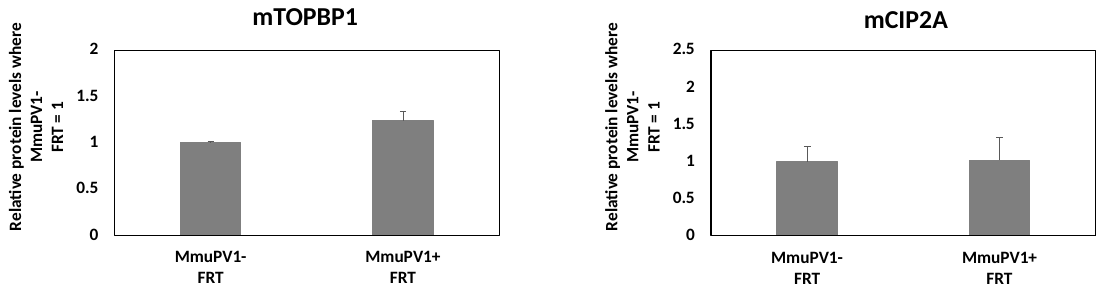

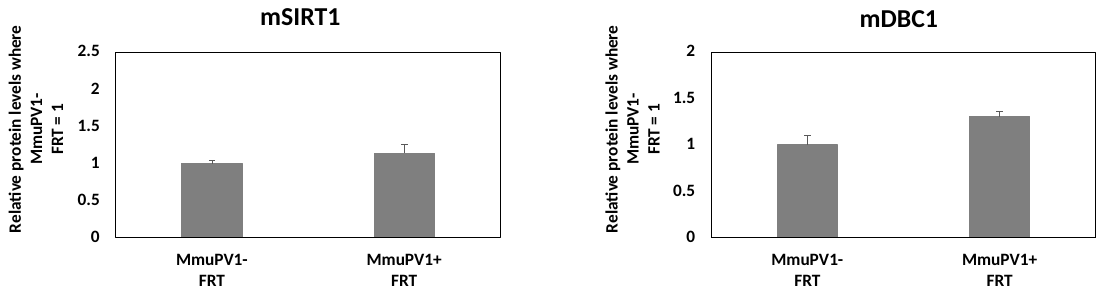

E	F	G

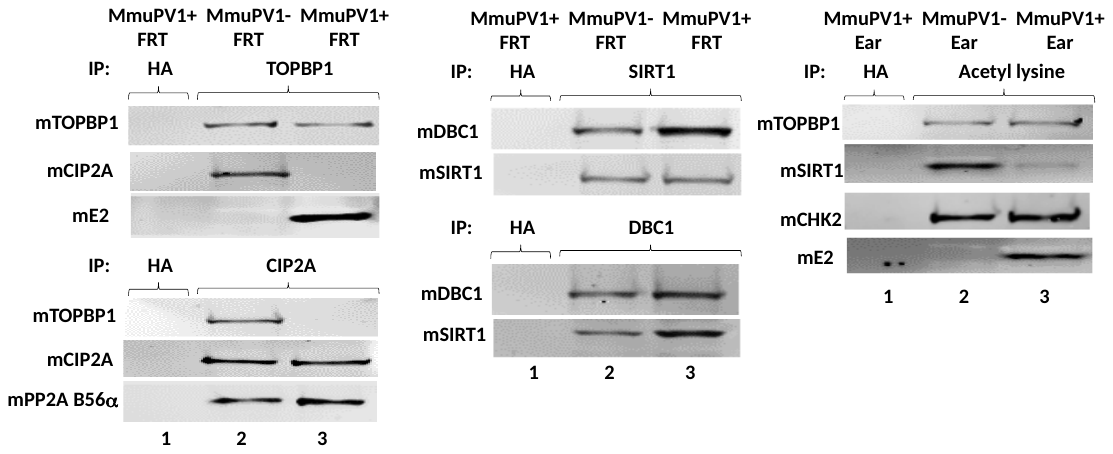

H

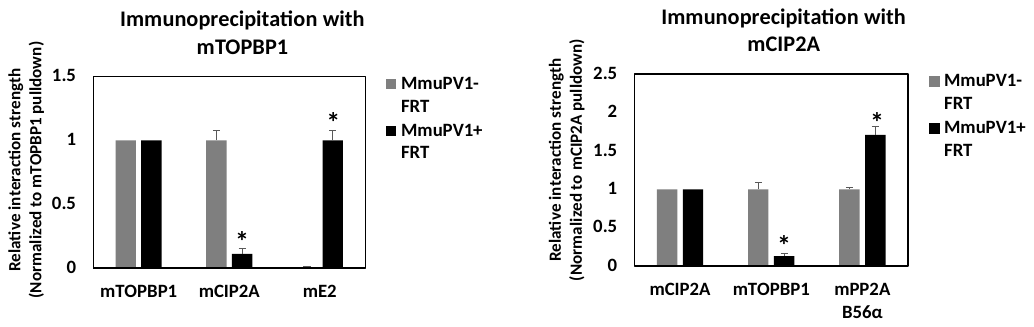

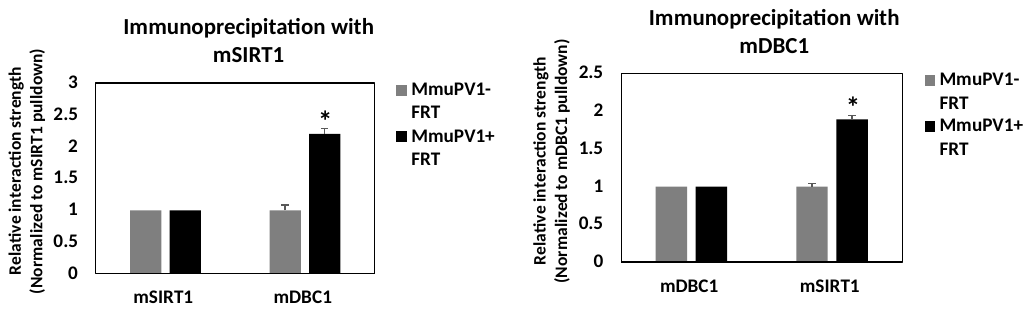

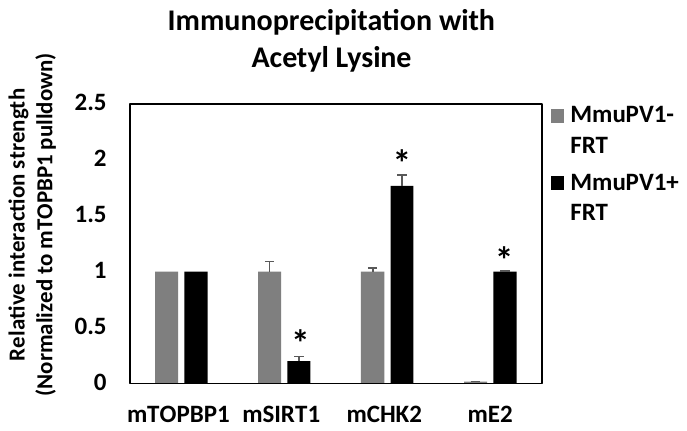

I

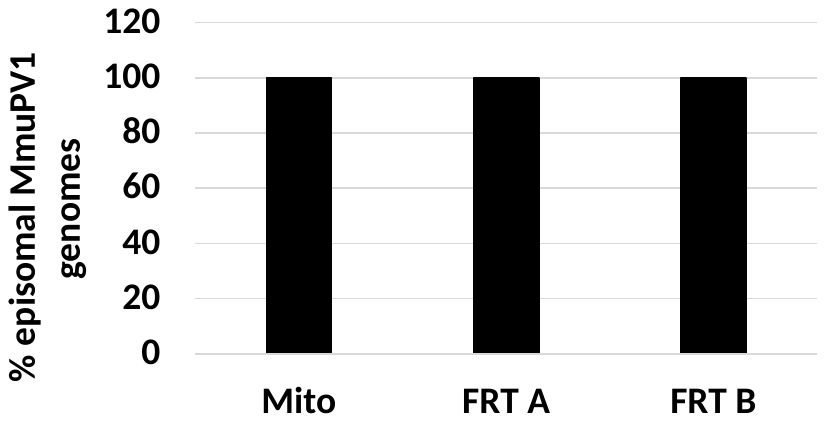

Figure S5. A. Another MmuPV1 FRT lesion (and control tissue) had protein extracted and the western blot in Figure 5C repeated. B. The results from Figures 5C and S5A were quantitated. C. Another MmuPV1 FRT lesion (and control tissue) had protein extracted and the western blot in Figure 5D repeated. D. The results from Figures 5D and S5C were quantitated. E., F., and G. The immunoprecipitations shown in Figures 5E-G were repeated with additional MmuPV1 and control FRT extracts. H. The immunoprecipitations in Figures 5E-G and S5-E-G were quantitated. Significant changes in the histograms are indicated by *, p-value<0.05. I. DNA was rescued from two FRTA MmuPV1 lesions and digested with TV exonuclease. GAPDH signal is diminished by 5-6 Ct values following TV treatment. There was zero degradation in mitochondrial (Mito) DNA, which is a circular 16kbp (episomal) molecule, and zero degradation of MmuPV1 DNA as determined by zero reduction in Ct signal following TV exonuclease treatment. The results represent two independent assays. There are zero error bars as there was 100% MmuPV1 episomal genomes in all assays.

Figure S6

A

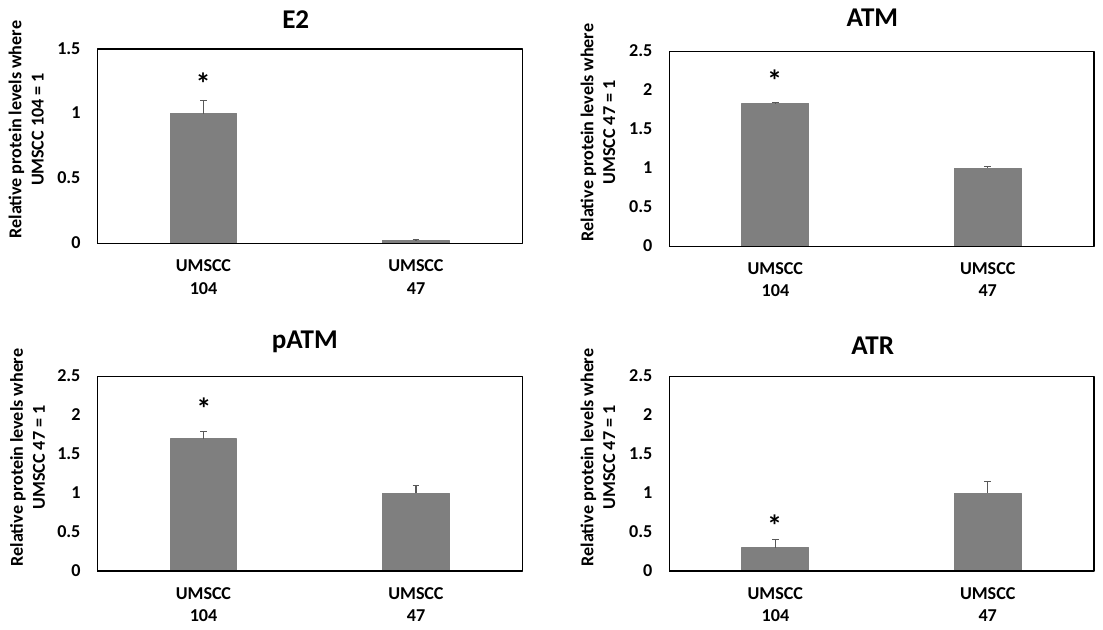

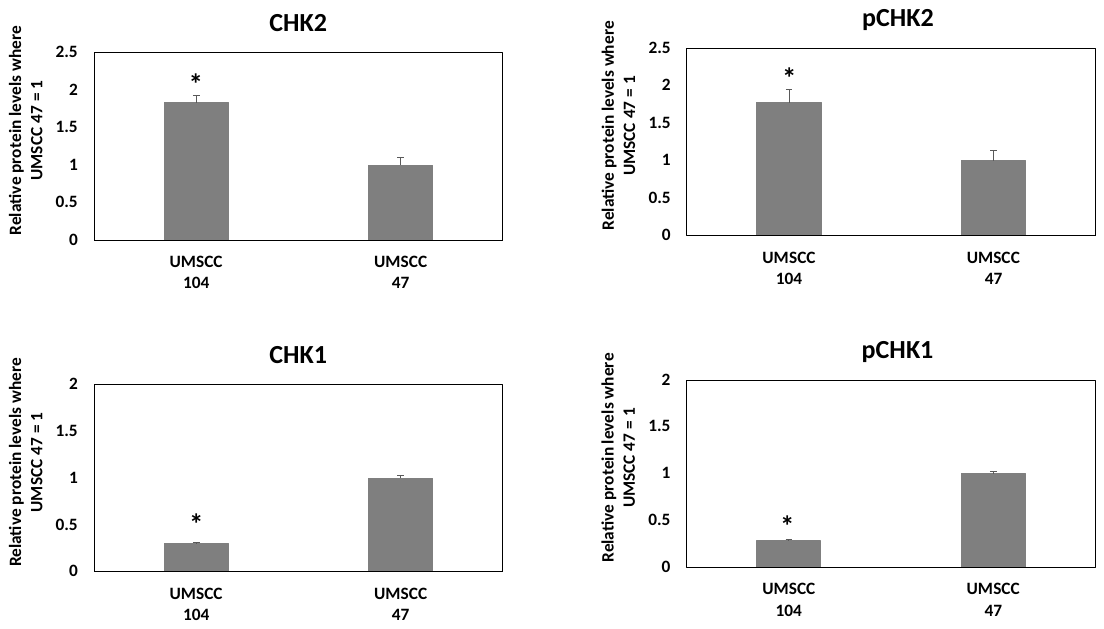

B

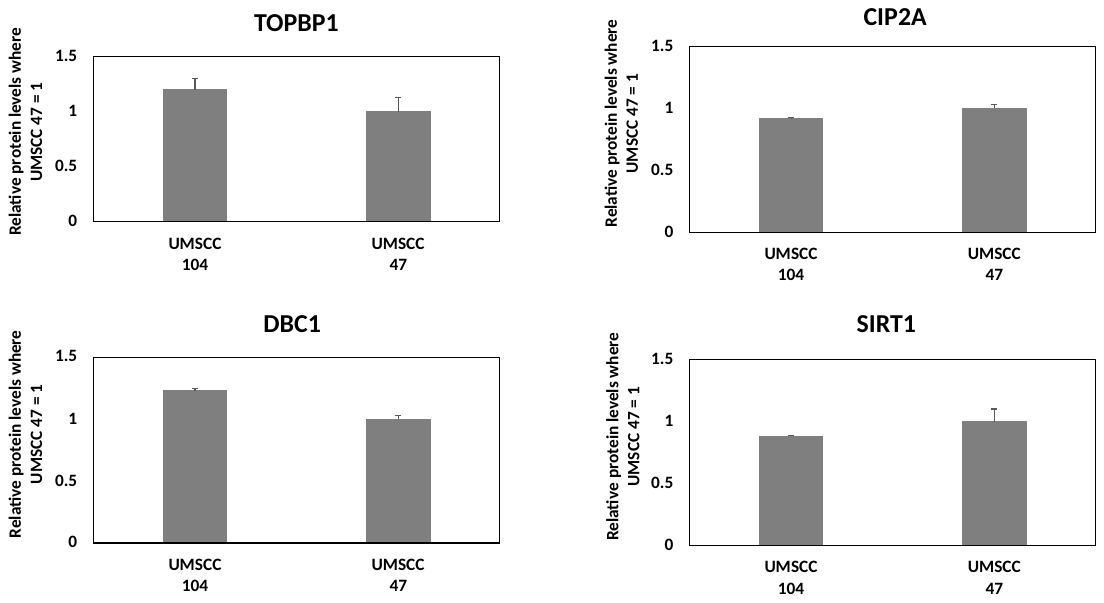

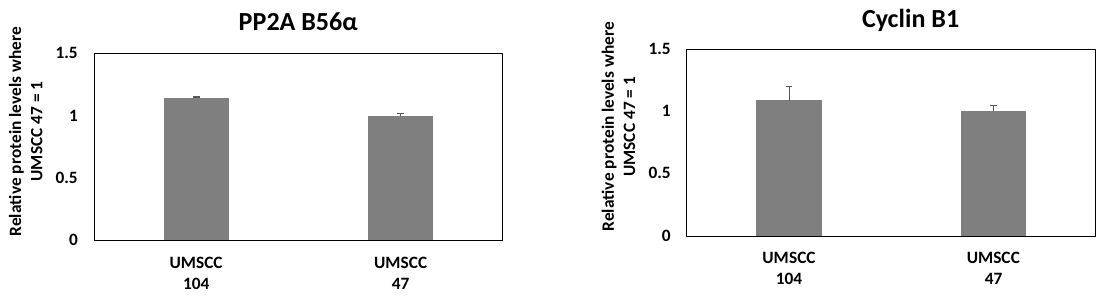

C

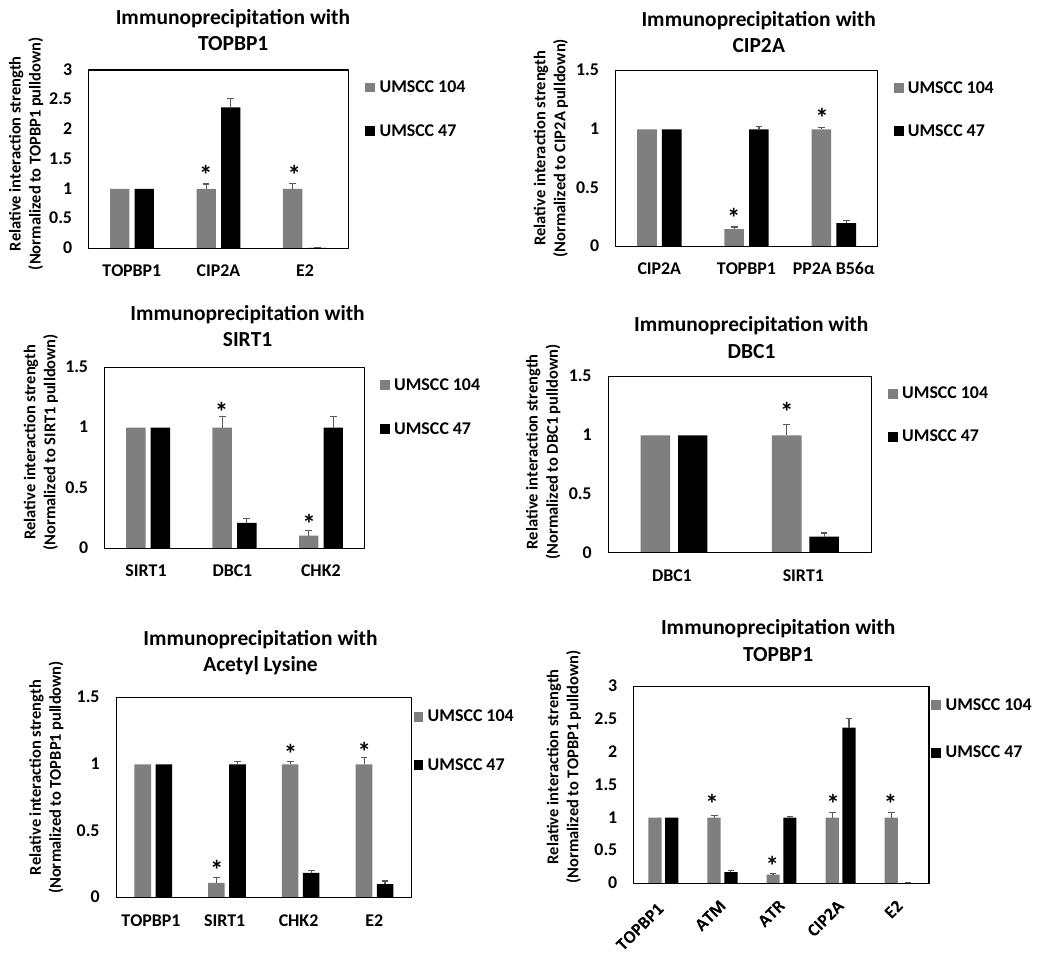

D

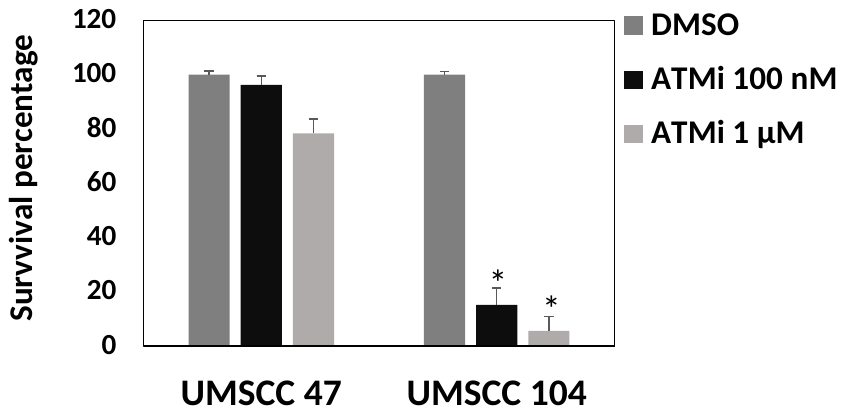

E

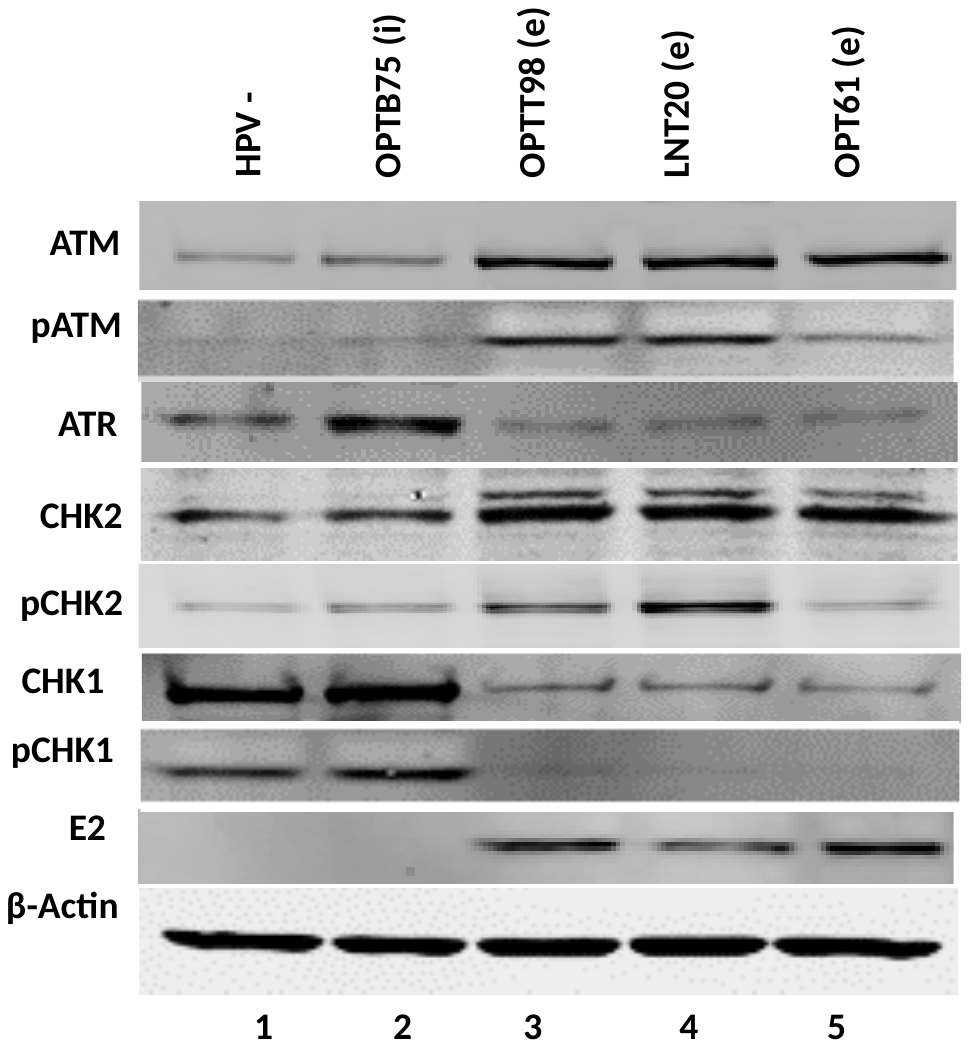

F

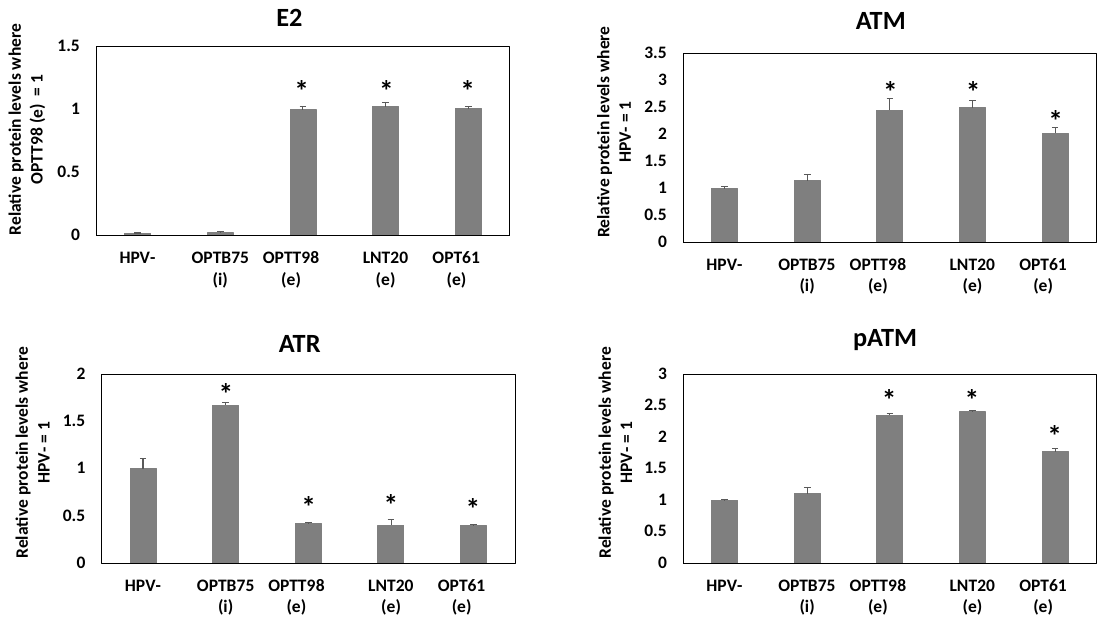

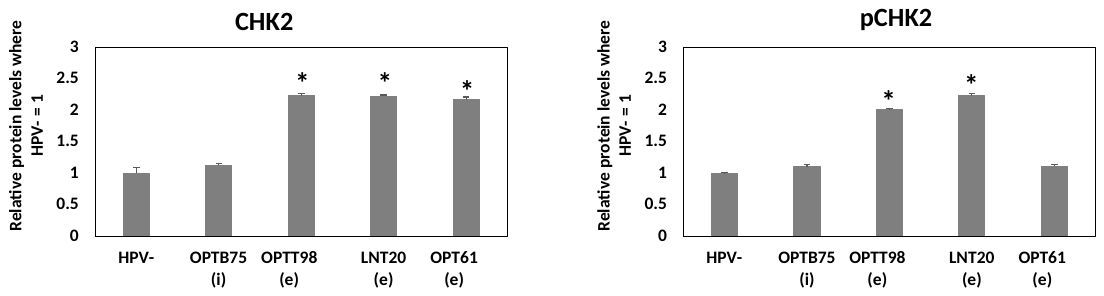

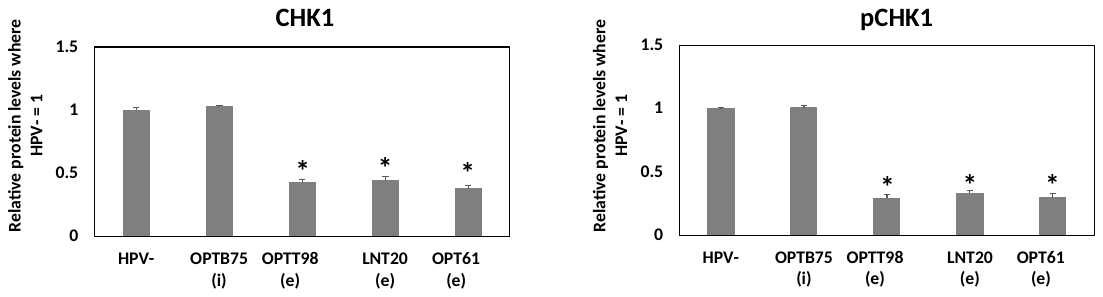

G

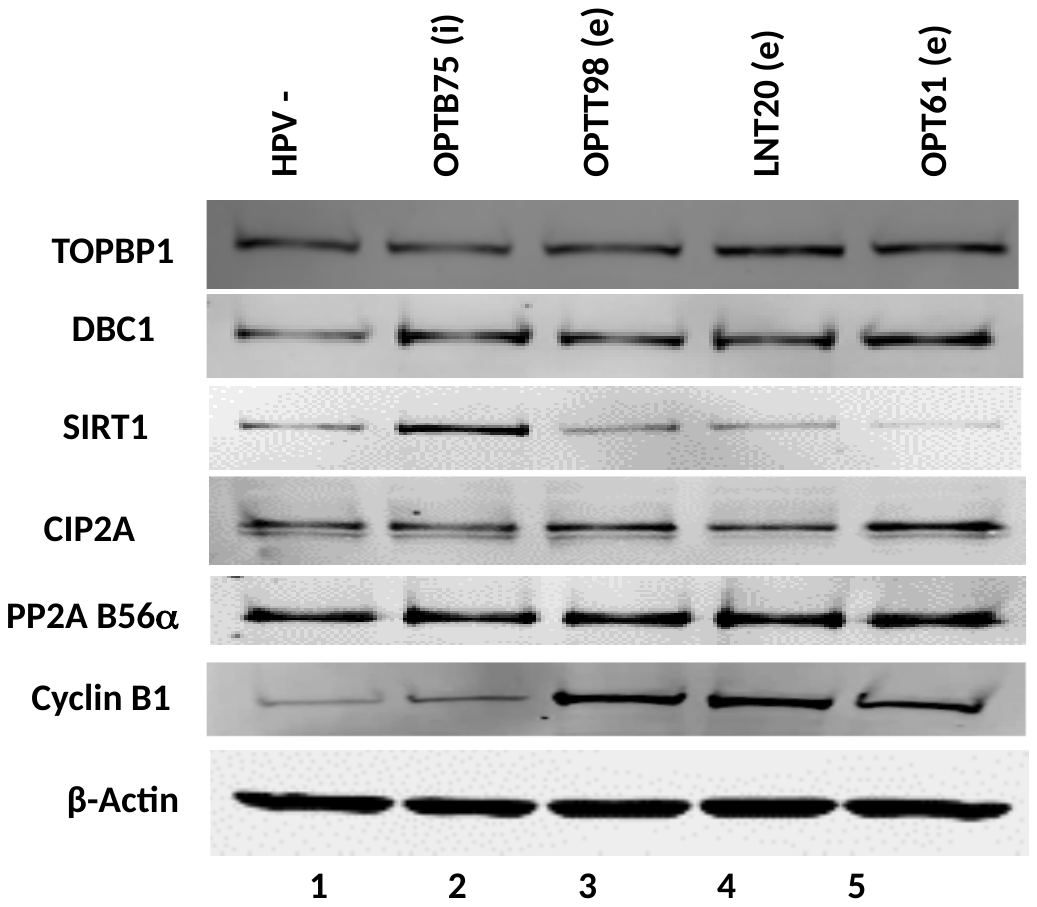

H

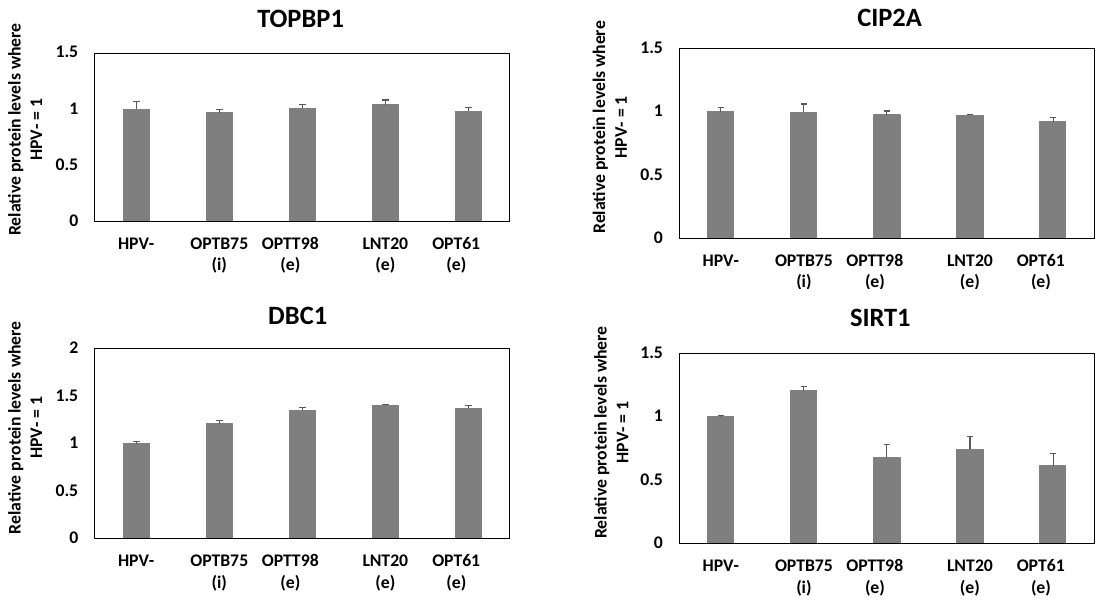

I

Figure S6. A. The experiment in Figure 6A was repeated and densitometry used to determine the levels of proteins indicated relative to UMSCC47 equaling 1, apart from E2 where the UMSCC-104 levels equaled 1. B. The experiment in Figure 6B was repeated and densitometry used to determine the levels of proteins indicated relative to UMSCC47 equaling 1. C-F. The experiments in Figures 6C-F were repeated and the results quantitated. D. The experiment in Figures 6H was repeated and the results quantitated. D. Protein extracts were prepared from the indicated HPV16 positive oropharyngeal patient derived xenografts (HPV16+OPC PDX), lanes 2-5, and from an HPV negative OPC PDX, lane 1. i means the viral genome is integrated (OPTB75, lane 2), while e indicates there is an episomal genome. Western blotting for the indicated proteins was carried out. F. The experiment in Figure S6E was repeated and quantitated. G. The OPC PDX extracts were blotted for the indicated proteins. H. The experiment in Figure S6G was repeated and quantitated. I. The indicated immunoprecipitations were carried out with the OPC PDX extracts. Significant changes in the histograms are indicated by *, p-value<0.05.
